# Frequency Spectra and the Color of Cellular Noise

**DOI:** 10.1101/2020.09.15.292664

**Authors:** Ankit Gupta, Mustafa Khammash

**Affiliations:** Department of Biosystems Science and Engineering, ETH Zürich, Mattenstrasse 26, 4058 Basel, Switzerland

**Keywords:** Power Spectral Density, Frequency Spectrum, Stochastic Reaction Networks, Time Lapse Microscopy, Systems Biology, Synthetic Biology

## Abstract

The invention of the Fourier integral in the 19th century laid the foundation for modern spectral analysis methods. By decomposing a (time) signal into its essential frequency components, these methods uncovered deep insights into the signal and its generating process, precipitating tremendous inventions and discoveries in many fields of engineering, technology, and physical science. In systems and synthetic biology, however, the impact of frequency methods has been far more limited despite their huge promise. This is in large part due to the difficulties encountered in connecting the underlying stochastic reaction network in the living cell, whose dynamics is typically modelled as a continuous-time Markov chain (CTMC), to the frequency content of the observed, distinctively noisy single-cell trajectories. Here we draw on stochastic process theory to develop a spectral theory and computational methodologies tailored specifically to the computation and analysis of frequency spectra of noisy cellular networks. Specifically, we develop a generic method to obtain accurate Padé approximations of the spectrum from a handful of trajectory simulations. Furthermore, for linear networks, we present a novel decomposition result that expresses the frequency spectrum in terms of its sources. Our results provide new conceptual and practical methods for the analysis and design of noisy cellular networks based on their output frequency spectra. We illustrate this through diverse case studies in which we show that the single-cell frequency spectrum facilitates topology discrimination, synthetic oscillator optimization, cybergenetic controller design, systematic investigation of stochastic entrainment, and even parameter inference from single-cell trajectory data.

## 1 Introduction

Modern microscopy and the advent of a wide array of fluorescent proteins has afforded scientists the unprecedented ability to monitor the dynamics of living biological cells [1]. The rapid pace of development in imaging technology coupled with advanced image processing techniques has made it viable to obtain high-resolution time-lapse live-cell data for a multitude of cell-types and biological processes [2]. Recent innovations in microfluidics make it possible to quantitatively measure single-cell dynamics for long periods of time over multiple generations [3]. These trends underscore the need for developing theoretical and computational tools that are specifically geared towards quantitatively extracting information about intracellular networks from live single-cell imaging data. One of the main reasons why the development of such tools is mathematically challenging is that the dynamics of single-cells is inherently noisy due to randomness in molecular interactions that constitute intracellular processes, and hence single-cell dynamics must be described with stochastic models that are more difficult to analyse than their deterministic counterparts [4]. These stochastic models usually represent the reaction dynamics as a continuous-time Markov chain (CTMC) and the existing methods for analysing them have mostly focussed on solving the Chemical Master Equation (CME) that governs the evolution of the probability distribution of the random state [5]. While these methods have been successfully applied in several significant biological studies [6, 7], they typically do not account for temporal correlations in time-traces of living cells, but rather they are designed to connect network models to flow-cytometry data [8] where temporal correlations are anyway lost due to discarding of the measured cells. Temporal correlations are a feature of single-cell trajectories that contain valuable information about the underlying network, and in order to access this information we need computational methods that can efficiently deduce the temporal correlation profile from a given stochastic reaction network model.

### Box 1

#### Frequency domain analysis of stochastic signals

Consider a reaction network, comprising species **X**_1_*, …*, **X**_*d*_ whose copy-number dynamics is described by an ergodic *continuous-time Markov chain* (CTMC) (*X*(*t*))_*t≥*0_ with stationary distribution *π*. Our goal is to estimate the PSD which measures the strengths of oscillatory components of various frequencies in the output signal (*X_n_*(*t*))_*t≥*0_ tracking the copy-number trajectory for species **X**_*n*_. We first subtract the stationary mean 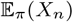 and construct the mean zero signal as 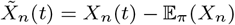 and then the time-averaged signal power *P*(*X_n_*) is equal to the stationary variance Var_*π*_(*X_n_*), i.e.

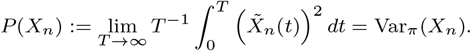

The *power spectral density (PSD)* for the output signal is given by

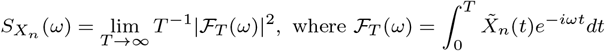

is the one-sided Fourier Transform, *ω* is the frequency and 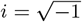. This PSD is related to the *autocovariance function*

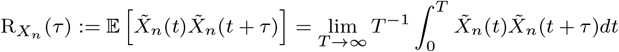

by the well-known *Wiener-Khintchine Theorem* [9] that shows that the PSD can be expressed as the two-sided Fourier Transform of the autocovariance function

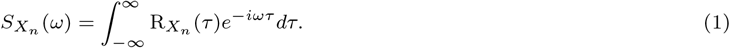

The interpretation of the PSD curve is given below. The location *ω*_max_ of its global maximum is considered to be the oscillatory frequency of the output signal.

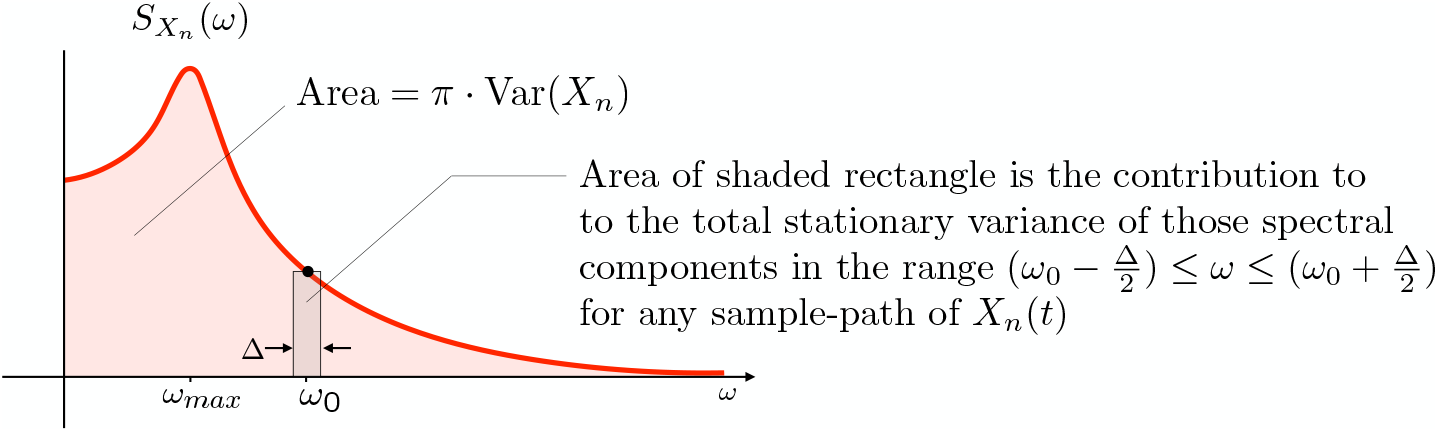

Commonly the PSD is estimated by first sampling a discrete time-series from a simulated CTMC trajectory at steady-state, and then taking its *Discrete Fourier Transform (DFT)* to estimate 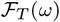 which then yields the PSD. This nonparametric procedure for PSD estimation is often called the *periodogram* method and it has known drawbacks due to estimator bias and inconsistency that often manifests in a high variance of the PSD estimator. The reliability of the estimator can be improved by ensemble averaging, windowing or artificial smoothing [10], but the underlying problems that compromise the accuracy of the PSD estimate still remain.

As is well-known in engineering and physics communities among many others, frequency-domain analysis is a powerful way to analyse random signals and systematically study temporal correlations. In particular, a signal’s *power spectral density (PSD)* measures the power content at each frequency, and it is related to the signal’s temporal *autocovariance function* via the Fourier Transform (see Box 1). The PSD of a single-cell trajectory is intimately related to the underlying network’s architecture and parametrisation *within the observed cell* [11]. There exist many studies that have successfully unravelled this relationship and discovered mechanistic principles for specific examples of reaction networks. For example, in [12] the role of feedback-induced delay in generating stochastic oscillations is explored and in [13] a stochastic amplification mechanism for oscillations is found. Notably, the exact PSD for linear reaction networks was derived in [14] and this was used to show how in gene expression networks post-translational modification reaction reduces the noise by serving as a low-pass filter.

Other works in this direction have relied on approximating the CTMC with a *stochastic differential equation* (SDE) such as the Linear Noise Approximation (LNA) [15] or the chemical Langevin equation (CLE) [16]. With these SDE-based approaches the protein PSD for gene-regulatory networks was investigated in [17, 18, 19], the relationship between input and output PSD for a single-input single-output system was computed in [20], the single-cell PSD for a general biomolecular network in the vicinity of a deterministic Hopf bifurcation was determined in [21] and corrections to the LNA-based PSD estimates were systematically derived in [22]. Even though SDE approximations make the problem of computing the PSD analytically tractable, their accuracy is severely compromised if any of the species are in low copy-numbers [23], as is the case for many synthetic networks where low copy-numbers are desired in order to reduce metabolic load on the host cell [24]. Moreover, even when the species copy-numbers are uniformly large, the accuracy of SDE approximations can only be guaranteed over finite time-intervals [25], and hence the PSD, which is estimated at steady-state, could have an error. In order to address these issues, we need PSD estimation methods that work reliably with CTMC models, especially in the low copy-number regime, without requiring any dynamical approximations. The aim of this paper is to develop such a method.

In a recent paper [26], the analytical relationship between the PSDs of the output species and its time-dependent production rate was derived for CTMC models of certain reaction networks including birth-death and simple gene-expression. While this analysis enables investigation of the dynamics of the protein creation process from experimentally measured protein time-traces, it does not extend to nonlinear networks, such as gene expression networks with transcriptional feedback, for which some analytical results exist for simplified models [27].

A recurring theme in the existing literature is that typically the autocovariance function is well-approximated by the sum of a few exponential functions [26, 20, 18], and consequently the PSD is a rational function of a special form. This low dimensional feature can be theoretically explained by appealing to the compactness of the resolvent operator [28] associated with the CTMC, which we as prove, is connected to the PSD. Exploiting this connection we develop the two-point Padé approximation [29] technique for estimating the PSD for a general nonlinear stochastic reaction network. This method, which we refer to as *Padé PSD*, computes the PSD expression based on certain stationary expectations. We design efficient Monte Carlo estimators to estimate the required expectations by generating a handful of simulations of an augmented CTMC, constructed by adding certain state-components and reactions to the original CTMC. We show how this augmented CTMC construction not only facilitates PSD estimation but also its empirical validation.

Our PSD estimation approach is *semi-analytic*, in the sense that analytical expressions for the PSD are found by first estimating certain quantities with simulation. Such approaches have become increasingly popular in recent years, as they provide viable solutions to nonlinear problems which are otherwise analytically intractable [30]. Analytical expressions for the PSD are known in the special case of linear reaction networks [14], where all reaction propensity functions are affine functions of the state variables. We show how this expression can be alternatively derived via the resolvent connection and we also generalise this result to allow for arbitrary time-varying inputs. This generalisation yields a novel PSD decomposition result that is similar to what was found in previous SDE-based studies [20] and it extends the recent results in [26].

Given a stochastic reaction network model, commonly the single-cell PSD is estimated with nonparametric methods by first simulating a trajectory, and then sampling it at finitely-many timepoints to obtain a discrete time-series whose PSD can be straightforwardly computed with the Discrete Fourier Transform (DFT) [31]. Either one can apply the DFT directly to the time-series to estimate the PSD or one can first estimate the autocovariance function and then compute its DFT (see Box 1 for more details). While the latter approach is computationally very expensive due to the autocovariance function computation, the former approach yields an inconsistent estimator for the PSD, which implies that the estimator variance does not vanish, even as the time-series length tends to infinity. To mitigate this inconsistency issue, PSDs from several independent trajectories are averaged, at the cost of significant computational burden as trajectory simulations are time-consuming. More importantly, the averaged PSD may still not be accurate because it is based on discrete sampling of continuous signals that can cause the problem of *aliasing* which distorts the estimated PSD by introducing frequency components corresponding to the sampling operation (see Chapter 1 in [10]). As shown by the Nyquist’s Sampling Theorem [32] we can mitigate this aliasing effect by choosing the time-step parameter that is smaller than half of the reciprocal of the maximum frequency represented in the signal. However for stochastic dynamics this criterion is unusable as the range of frequencies in the signal is very wide and picking a very small time-step can lead to computational intractability. These issues motivated us to devise Padé PSD that is not based on discrete-sampling and provides a parametric approach for estimating the PSD that rather than relying on only the output signal, uses full information contained in the stochastic model of the dynamics.

We illustrate our results with applications of relevance to both systems and synthetic biology. Using our PSD decomposition result for linear networks, we demonstrate how PSDs enable differentiation between two fundamental types of adapting circuit topologies, viz. Incoherent Feedforward (IFF) and Negative Feedback (NFB) [33], in the presence of dynamical intrinsic noise. We also present an example where the phenomenon of single-cell entrainment is examined in the stochastic setting using our PSD decomposition result. Employing Padé PSD we illustrate how the performance of certain synthetic circuits, with noisy dynamics, can be optimised. Specifically we examine the problem of optimising the oscillation strength of a well-known synthetic oscillator (called the *repressilator* [34]) and the problem of reducing single-cell oscillations which can arise when an intracellular network is controlled with the *antithetic integral feedback* (AIF) controller [35] that has the important property of ensuring robust perfect adaptation despite randomness in the dynamics and other environmental uncertainties. Lastly we present a simple example to highlight how our Padé PSD method can facilitate parameter inference from experimentally measured single-cell trajectories, by providing providing clean and accurate estimations of the PSD. Interestingly, a parameter is identified in this example without the explicit knowledge of the proportionality constant that relates the measured signal to the copy-number of the output species.

## 2 The resolvent representation of the PSD

In this section, we describe the CTMC model for a reaction network and define the resolvent operator associated with it. We then connect this operator to the PSD. This connection shall be exploited in later sections to develop our analytical and computational PSD results.

### 2.1 The Stochastic Model

Consider a reaction network with *d* species, called **X**_1_, … ,**X**_*d*_, and *K* reactions. In the classical stochastic reaction network model, the dynamics is described as a *continuous-time Markov chain* (CTMC) [5] whose states represent the copy numbers of the *d* network species. If the state is *x* = (*x*_1_, … ,*x_d_*) and reaction *k* fires, then the state is displaced by the integer stoichiometric vector ζ_*k*_. The rate of firing for reaction *k* at state *x* is governed by the propensity function λ_*k*_(*x*). Under the mass-action hypothesis [5]

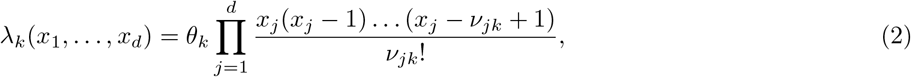

where *θ_k_* is the rate constant and ν_*jk*_ is the number of molecules of **X**_*j*_ consumed by the k-th reaction. Formally, the CTMC (*X*(*t*))_*t≥*0_ representing the reaction kinetics can be defined by its generator 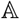, which is an operator that specifies the rate of change of the probability distribution of the process (see Chapter 4 in [36]). It is defined by

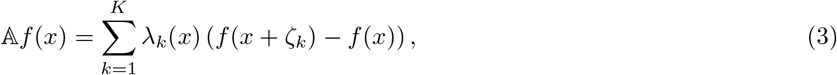

for any real-valued bounded function *f* on the state-space which consists of all accessible states in the *d*-dimensional nonnegative integer lattice.

For each state *x*, let *p*(*t, x*) be the probability that the CTMC (*X(t)*)_*t≥*0_ is in state *x* at time *t*. Then these probabilities evolve according to a system of ordinary differential equations, called the Chemical Master Equation (CME) [5], which is typically unsolvable. Hence its solutions are often estimated with Monte Carlo simulations of the CTMC, using methods such as Gillespie’s *stochastic simulation algorithm* (SSA) [37]. If the CME has a unique, globally attracting fixed point π then the CTMC is called *ergodic* with π as the stationary distribution. If the convergence of *p(t)* to π is exponentially fast in t, then the CTMC is called exponentially ergodic and the exponential rate of convergence is called the *mixing strength* of the CTMC. We shall work under the assumption of exponential ergodicity which is computationally verifiable using techniques in [38] and [39], wherein, it is also demonstrated that this assumption is satisfied by networks typically encountered in systems and synthetic biology. It is important to note that for an ergodic network, all stochastic trajectories, despite being different, have the same PSD.

### 2.2 The Resolvent Operator and its connection to the PSD

Let (*X(t)*)_*t*≥0_ be a CTMC with generator 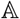. For such a Markov process, we define the transition semigroup 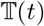 as the operator which maps any real-valued function g on the state space, to the function specified by the conditional expectation

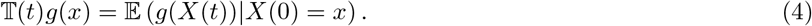

We now define the resolvent operator which plays a central role in the development of our method for PSD estimation. For any complex number *s*, the resolvent operator maps the function *g* to the Laplace transform of the map 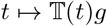

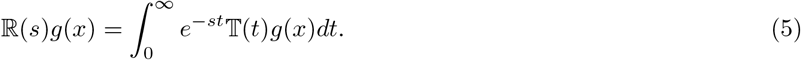

It can be shown that the map *s* ↦ ℝ(*s*)*g*(*x*) is complex-analytic.

Assuming that the observed single-cell trajectory (*X_n_*(*t*))_*t≥*0_ is the copy-number dynamics of the output species **X**_*n*_, we now establish a relation between the PSD 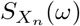 (see Box 1) and the resolvent operator. Let 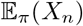 denote the stationary expectation of the copy-number of species **X**_*n*_ and let *f* be the function

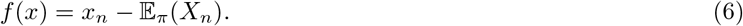

Defining

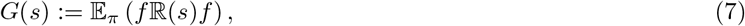

the PSD 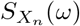 is given by

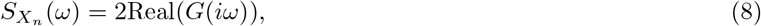

where 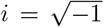. This relation is proved in Section S2.2 of the Supplement. In this result we view the function *x* ↦ *f*(*x*)ℝ(*s*)*f*(*x*) as a random variable on the probability space whose sample-space is the state-space of the CTMC and the probability distribution is given by the stationary distribution π. The expectation of this random variable is denoted by G(s) and in the PSD estimation method we develop, we first estimate G(s) and then obtain the PSD using (8).

The eigen-decomposition of the resolvent operator allows us to express *G*(*s*) as an infinite sum

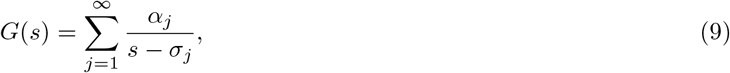

where σ_1_, σ_2_, … are the non-zero eigenvalues of 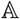, arranged in descending order of their real parts (which are negative due to ergodicity). Each coefficient α*j* captures the power in the signal corresponding to eigenmode σ*j*, and their sum is equal to the total signal power which is also the stationary variance Var_*π*_(X_*n*_) of the output species copy-number

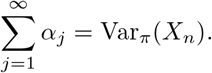

Relation (9) is equivalent to the following representation of the autocovariance function

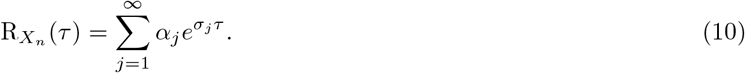

In the case of linear networks, *G*(*s*) can be exactly computed and (8) yields an analytical expression for the PSD which is already known in the literature [14]. However for linear networks stimulated by external inputs it is not known how the output PSD is related to the PSDs of the input signals. We derive this relation by exploiting the resolvent connection and this yields a novel and practically useful PSD decomposition result (see Section 3). For general nonlinear networks, we apply the theory of Padé approximations to find an accurate rational function representation of *G*(*s*) (see Section 4) which is then used to estimate the PSD (8).

## 3 A PSD decomposition result for linear networks

In this section we present a novel PSD decomposition result for linear networks, that extends a similar result recently reported in [26]. A reaction network is called linear if all its propensity functions are affine functions of the state variables. Under mass-action kinetics, linear networks are necessarily unimolecular, i.e. all reactions have at most one reactant and are of the form ∅ → ∗ or **X**_*j*_ → ∗, where ∗ represents any linear combination of species. Assuming *d* species and *K* reactions, for linear networks we can express the vector of propensity functions λ(*x*) = (λ_1_(*x*), … , λ_*K*_(*x*)) as an affine map on the state-space

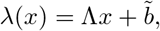

where Λ is some *K* × *d* matrix and 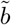 is a *K* × 1 vector. Letting *S* be the *d* × *K* matrix whose columns are the stoichiometric vectors ζ_1_, … , ζ_*K*_ for the reactions. We define

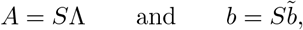

and under the assumption of ergodicity, the *d* × *d* matrix *A* is Hurwitz-stable, i.e. all its eigenvalues have strictly negative real parts. It can be easily shown (see [38] for e.g.) that the dynamics of the expected state 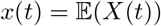 is given by

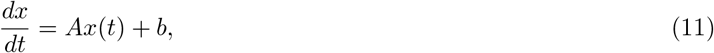

and as *t* → ∞, *x*(*t*) converges to 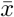 which is the state expectation under the stationary distribution π

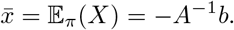

Moreover the stationary covariance matrix Σ for the state can be computed by solving the following Lyapunov equation

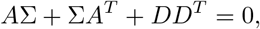

where *D* is the positive semidefinite matrix satisfying 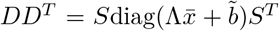. In this setting, we can show that the resolvent operator maps the class of affine functions to itself, and this allows us to apply formula (8) to prove (see the Supplement, Section S2.3) that the PSD is given by

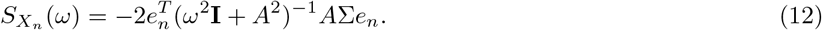

where **I** is the *d* × *d* identity matrix and *e_n_* denotes its *n*-th column. This expression is equivalent to the PSD formula for linear networks proved in [14] using Gardiner’s *regression theorem* [40].

Now consider the situation where such a linear network is being driven by external signals. These signals could be generated by different sources, e.g. upstream interconnected networks, environmental stimuli, or by engineered inputs introduced to probe the dynamics (see Figure 1). A fundamentally important question is to understand how the internal noise and each of these inputs (deterministic or stochastic) conspire to make up the full power spectrum of an output of interest. Indeed it would be of considerable conceptual and practical significance to be able to decompose the output power spectrum in a way that allows the quantification of the specific contributions to the spectrum of the internal noise and of each of the external inputs. Although approximate decompositions of this sort have been reported in specific example networks modeled by CLEs [20, 19], to the best of our knowledge no spectral decomposition results exist for general biochemical networks modeled by CLE, nor for those modeled by discrete stochastic CTMC models.

**Figure 1:**
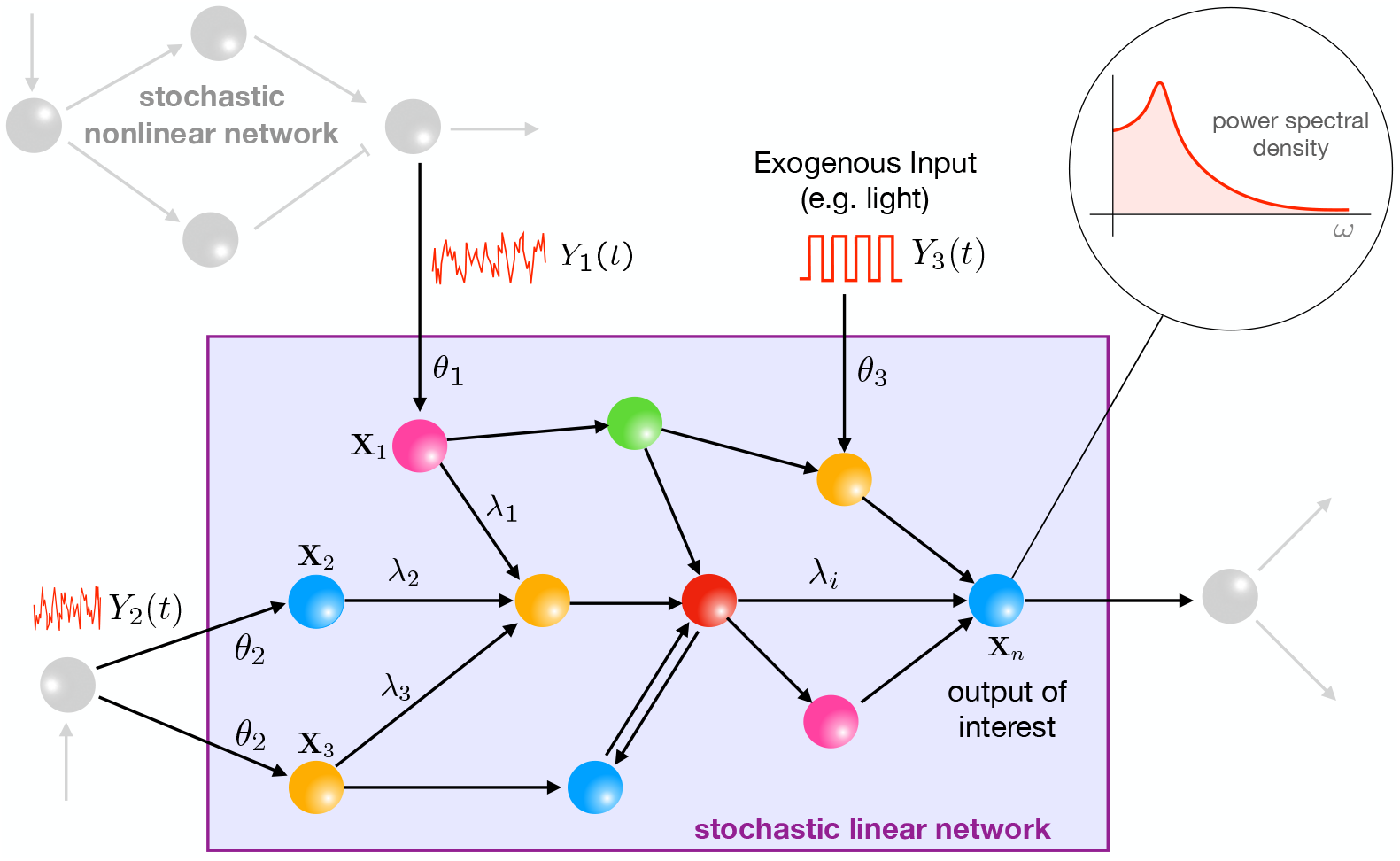
The setting of the PSD decomposition result: A stochastic reaction network with linear propensity functions embedded in the intracellular milieu and receiving stimulation from several upstream networks. Theorem 3.1 provides an analytical decomposition for the output PSD 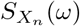 in terms of the PSDs of all the stimulating signals.

We consider *m* independent time-varying signals (*Y*_1_(*t*))_*t≥*0_, … , (*Y_m_*(t))_*t≥*0_. We assume that these signals stimulate through *m* zeroth-order reactions of the form

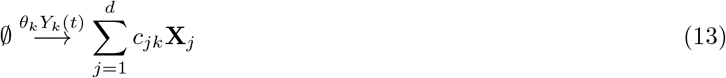

for *k* = 1, … , *m*. Each reaction follows mass-action kinetics and for reaction *k*, *θ_k_* is a positive constant and *c_k_* = (*c*_1*k*_, … , *c_dk_*) is the vector representing the number of molecules of each species **X**_1_, … , **X**_*d*_ created by this reaction. We shall assume that process (*Y*(*t*))_*t≥*0_, which includes all the stimulating signals, is an exponentially ergodic Markov process with stationary expectation 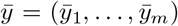. Let 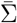 be the stationary variance-covariance matrix for the process (*X*(*t*))_*t≥*0_ *when each stimulating signal is deterministic and fixed to its stationary mean at all times*, i.e. 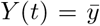 for all t ≥ 0. We now present our main result for linear networks which provides an analytic relationship between the PSD 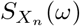 of our output species **X**_*n*_ and the of PSDs 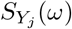 for j = 1, … , *m*.

### Theorem 3.1 (PSD Decomposition)

*Consider a linear reaction network comprising species* **X**_1_, … , **X**_*d*_, *stimulated by independent time-varying signals (Y_1_(t))_*t≥*0_, … , (Y_*m*_(t))_*t≥*0_, through zeroth-order reactions of the form* (13). *We assume that each Y_j_ is an exponentially ergodic Markov process with PSD* 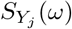. *The PSD of the output species* **X***_n_ is given by*

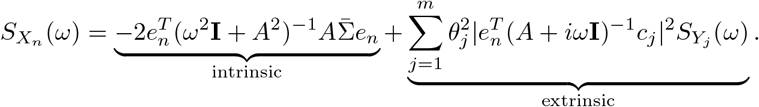

The proof of this result is provided in Section S2.3 in the Supplement and it shows that the output spectrum is the sum of the intrinsic contribution and the external contributions from all stimulating signals. The external contribution due to signal *Y_j_* is modulated by the frequency dependent gain 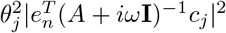.

## 4 Padé PSD: A spectrum estimation method for nonlinear networks

In this section we develop our framework, called *Padé PSD*, for estimating the PSD for a general nonlinear network by applying Padé approximation theory which is known to be immensely useful in computing accurate rational function approximations for analytic non-linear functions. In particular, we shall employ the method of two-point Padé approximation for finding a rational function approximant for the function *G*(*s*) (see (7)) which then provides the PSD (see (8)). In such an approximation, the rational approximant is constructed by matching its power series expansions at two arbitrarily chosen points, up to a certain number of terms, to the corresponding power series expansion of the function being approximated (i.e. *G*(*s*) in our case) [41]. The number of terms up to which each power series is matched is given by a pair **p** = (*p*_1_, *p*_2_) of nonnegative integers called the *order* of the Padé approximation. This nonnegative integer pair should have an even sum, and letting

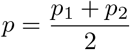

be the arithmetic mean of these two integers, the order **p** Padé approximant has the form

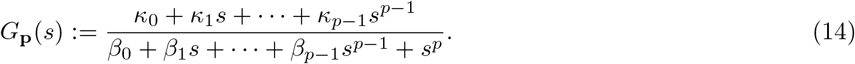

Notice that the degree of the numerator polynomial is (*p* − 1) while the degree of the denominator polynomial is *p*. We shall now outline how the 2*p* = (*p*_1_ + *p*_2_) coefficients κ_0_, … , κ_*p−*1_, β_0_, … , β_*p−*1_ can be reliably estimated and how the resulting Padé approximant can be validated. For more details we refer the readers to Section S2.4 in the Supplement.

Let us first comment on why a rational function of the form might be an accurate approximation for the function *G*(*s*). Recall representation (10) of the autocovariance function which is equivalent to representation (9) for the function *G*(*s*). Previous studies have established that usually the autocovariance function is well-approximated by only the first few terms in this infinite series. This fact can be justified by appealing to the compactness of the resolvent operator which ensures that it is close to a finite-rank operator (see Section S2.1 in the Supplement). If we only keep the first *p* terms in the infinite sum (9), then we obtain a rational function of the form (14).

Since function *G*(*s*) is complex-analytic it suffices to estimate it on the real-line. In our method, the two points at which we match the power series expansions are given by a small positive real number *s*_0_ and ∞. This way we ensure that the constructed Padé approximant *G*_**p**_(*s*) can reliably describe the approximated function *G*(*s*) at both small and large values of *s*. Suppose *G*(*s*) has the following power-series expansion around *s* = *s*_0_

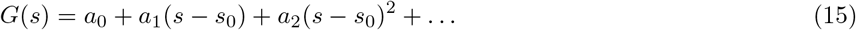

and the following power-series expansion for large *s*

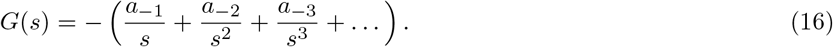

The order **p** = (*p*_1_, *p*_2_) Padé approximant *G*_**p**_(*s*) is such that its power series expansion at *s* = *s*_0_ agrees with the first p1 terms in (15) and its power series expansion at *s* = ∞ agrees with the first *p*_2_ terms in (16). As shown in [41], *G*_**p**_(*s*) can be constructed by defining two (*p* + 1) × (*p* + 1) matrices in terms of the power series coefficients *a*_0_, *a*_±1_, *a*_±2_. Let *Q*(*z*) be the (*p* + 1) × (*p* + 1) matrix given by

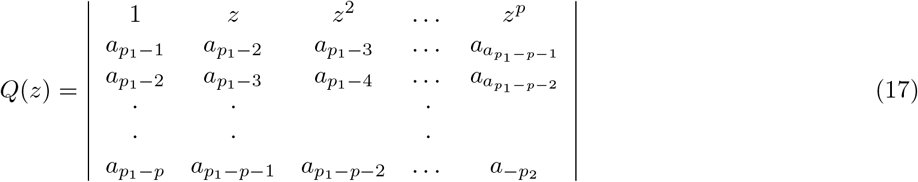

and let *P*(*z*) be the (*p* + 1) × (*p* + 1) matrix obtained from *Q*(*z*) by replacing its first row with the vector *v*(*z*) = (*v*_0_(*z*), … , *v_m_*(*z*)), where each *v_j_*(*z*), for *j* = 0, … , *p*, is defined as

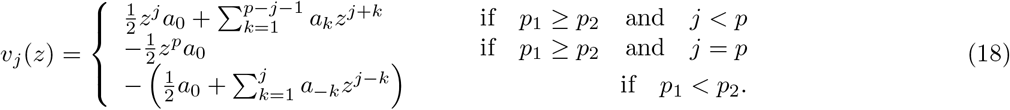

Then the order **p** = (*p*_1_, *p*_2_) Padé approximant *G*_**p**_(*s*) can be computed as

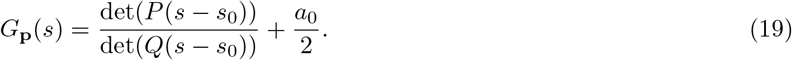

When either *p*_1_ = 0 or *p*_2_ = 0, *G*_**p**_(*s*) reduces to the classical one-point Padé approximant [42, 29].

Once the power series coefficients 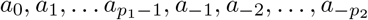 have been estimated, we can compute *G*_**p**_(*s*) using formula (19) and substituting this Padé approximant instead of *G*(*s*) in (8) yields an estimate of the PSD. The main challenge is to develop a method for reliable estimation of these power series coefficients from a handful of trajectory simulations. In the upcoming sections we describe such a method and also discuss how the resulting Padé approximant can be validated.

### 4.1 Estimation of the coefficients of power series (16)

Let 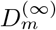 be the *m*-th Padé derivative at ∞ defined by

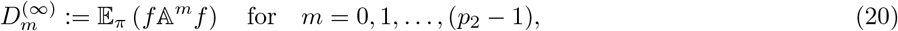

where *f* is the output function (6) and 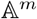 denotes the m-th iterate of the generator 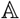 with 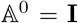 (the identity operator). Then it can be shown (see Section S2.4.2 in the Supplement) that the first *p*_2_ coefficients of the power series (16) are given by

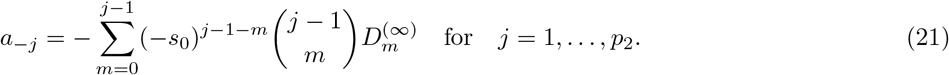

The steady-state expectation (20) can be simultaneously estimated for each *m* = 0, 1, … , (*p*_2_ − 1) from *Q* CTMC trajectories simulated over some large time-interval [0, *T_f_*]. Denoting these trajectories by (*X*^(*q*)^(*t*))_*t≥*0_ for 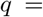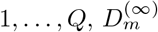 can be estimated by the Monte Carlo (MC) estimator

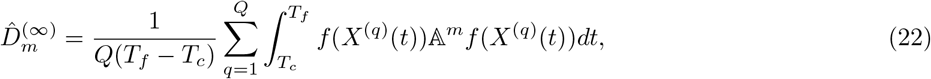

where *T_c_* ≪ *T_f_* is the cut-off time at which stationarity is assumed to be reached and the initial part of each trajectory in the time-interval [0, *T_c_*] is discarded. Observe that if *T_f_* is large enough then even a single trajectory (i.e. *Q* = 1) is sufficient for this estimation due to Birkhoff’s Ergodic Theorem [43]. However using multiple trajectories enhances the MC estimator’s statistical accuracy which can be measured by estimating its sample variance.

Generally we find that the estimator (22) has a very large variance unless the simulation time-period [0, *T_f_*] is extremely large. To mitigate this issue we design suitable covariates that can be added to the integrands in (22) in order to aid convergence with respect to *T_f_* (see Section S2.4.3 in the Supplement). The resulting integrand is given by

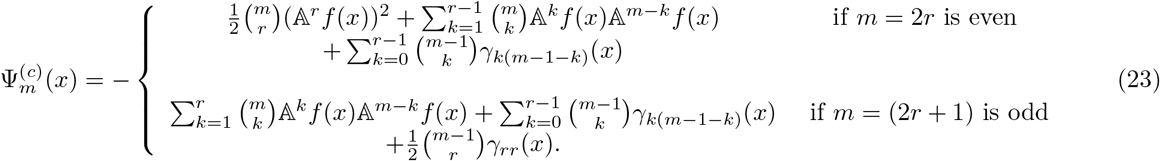

where the function *γ*_*jl*_(*x*) is defined as

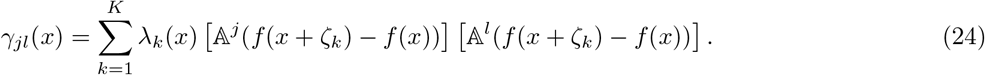

It can be shown that 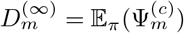 and hence we can estimate it from *Q* CTMC trajectories as

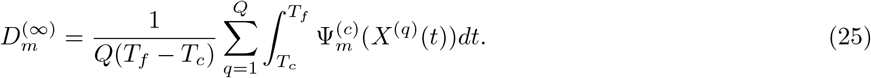

In practice we find that this covariate-based MC estimator (25) typically has much lower variance than the simpler MC estimator (22).

### 4.2 Estimation of the coefficients of power series (15)

Our next goal is to estimate the first *p*_1_ coefficients in the power series (15). It can be shown (see Section S2.4.2 in the Supplement) that these coefficients are given by

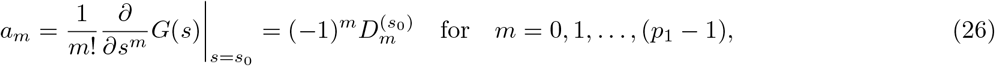

where 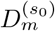 is the *m*-th Padé derivative at *s*_0_ defined by

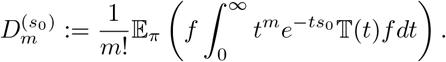

Here *f* is the output function (6) and 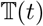 denotes the transition semigroup operator (4). Appealing to the ergodicity of the CTMC we can express 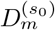 as

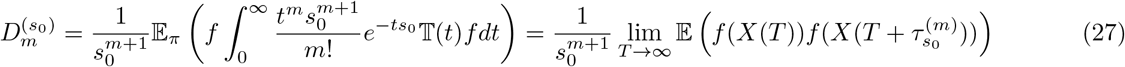

where 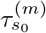 is an independent random variable with `Erlang` distribution with shape parameter (*m* + 1) and rate parameter *s*_0_. In other words, the probability density function of 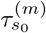 is given by

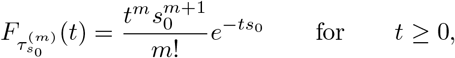

and we can view 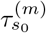 as the sum of (*m* + 1) independent and identically distributed exponential random variables with rate parameter *s*_0_.

We can estimate the steady-state expectation (27) simultaneously for each *m* = 0, 1, … , (*p*_1_ − 1). For this we augment the CTMC state with *p*_1_ additional state components, denoted by 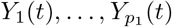, and an extra reaction, called 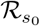 that fires at the constant rate of *s*_0_. If this reaction fires at time *t*, then we reset these additional state components as

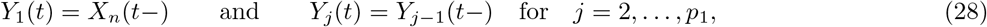

where *X_n_*(*t*−) is the copy-number of the output species **X**_*n*_, *just before* the reaction firing time. Similarly for *j* ≥ 2, *Y_j_*(*t*) assumes the value of the previous state component before the jump time, which is *Y*_*j−*1_(*t*). The steady-state expectation for each *Y_j_*(*t*) is the same as that of the output i.e. 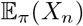, and for any *T*

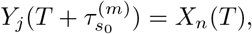

where 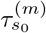 is the Erlang-distributed random variable mentioned above. Substituting this in (27), one can see that

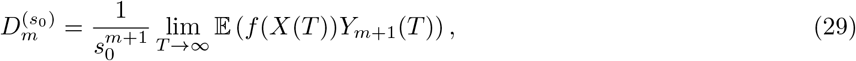

and hence it can be estimated from *Q* augmented CTMC trajectories, denoted by (*X*^(*q*)^(*t*), *Y*^(*q*)^(*t*))_*t≥*0_ for *q* = 1, … , *Q*

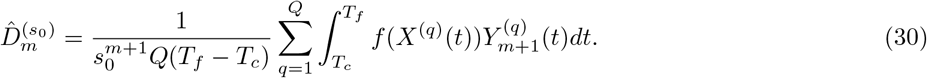

### 4.3 Validation of the Padé approximant

Once the required power series coefficients at *s*_0_ and ∞ have been estimated, as described in Sections 4.1 and 4.2, then we can compute the Padé approximant *G*_**p**_(*s*) with formula (19) and then use this approximant to compute the PSD. For this PSD estimation procedure to work well it is crucial that the Padé approximant *G*_**p**_(*s*) is an accurate surrogate for the function *G*(*s*), which depends on many factors, such as the order of the approximation **p** = (*p*_1_, *p*_2_), the length of the simulation time-period *T_f_*, and most importantly the statistical precision of the Padé derivative estimates.

In order to test if a computed Padé approximant is accurate we can validate it using direct statistical estimates (i.e. without rational approximation) of the function *G*(*s*) at multiple values of *s*, prescribed by some vector **s** = (*s*_1_, … , *s_R_*) of positive real numbers. Similar to Section 4.2, these direct estimates can be estimated by augmenting the CTMC state with R additional state components, denoted by *Z*_1_(*t*), … , *Z_R_*(*t*), to keep track of the copy number *history* of the output species **X**_*n*_ at random exponential times in the past. Assume that there are *R* additional reactions 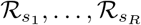 that fire independently at constant rates *s*_1_, … , *s_R_* respectively. If reaction 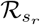 fires at time *t*, then we set

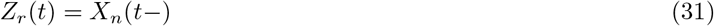

where *X_n_*(*t*−) is the copy-number of the output species **X**_*n*_, *just before* the reaction firing time. As in Section 4.2 we can conclude that for each *r* = 1, … , *R* the value *G*(*s_r_*) can be estimated with *Q* augmented CTMC trajectories, denoted by (*X*^(*q*)^(*t*), *Z*^(*q*)^(t))_*t≥*0_ for *q* = 1, … , *Q*

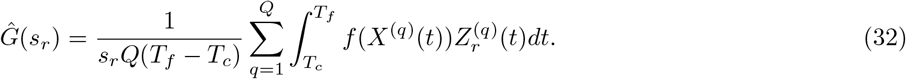

If the estimated Padé approximant *G*_**p**_(*s*) is accurate, each 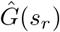 would be close to the value *G***p**(*s_r_*), even though both these estimates would have some inaccuracies due to finite sampling and the finiteness of the simulation time-period. Upon comparing the graphs 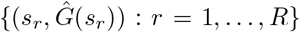 and {(*s_r_*, *G*_**p**_(*s_r_*)) : *r* = 1, … , *R*}, the Padé approximant can be validated. If the discrepancy is too high then it suggests that we either need to increase the approximation order *p*, or the simulation time *T_f_*, or both.

### 4.4 Computational implementation of *Padé PSD*

We now discuss the computational implementation of our PSD estimation method that we refer to as *Padé PSD*. The detailed algorithms for this method are provided in Section S3 of the Supplement and its full `Python` implementation is available on GitHub: https://github.com/ankitgupta83/PadePSD_python.git.

Suppose that the order **p** = (*p*_1_, *p*_2_) of the Padé approximant, the value *s*_0_ and the test values **s** = (*s*_1_, … , *s_R_*) are fixed. The main computational tasks that Padé PSD performs are:

1. **Estimate the required Padé derivatives:** Quantities 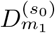 and 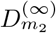 are estimated, for *m*_1_ = 0, 1, … , (*p*_1_ − 1) and *m*_2_ = 0, 1, … , (*p*_2_ − 1), as discussed in Sections 4.1 and 4.2 respectively.
2. **Obtain direct estimates for validation**: Quantities (*G*(*s*_1_), … , *G*(*s_R_*)) are directly estimated as discussed in Section 4.3.

Upon completing these tasks, the power series’ coefficients (20) and (26) are obtained, and the order **p** Padé approximant *G*_**p**_(*s*) is computed with (19). This Padé approximant is then validated with the direct estimates (*G*(*s*_1_), … , *G*(*s_R_*)), and if the validation is successful, the PSD 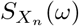 is obtained by applying formula (8) with *G*(*z*) = *G*_**p**_(*z*).

All the required quantities are simultaneously estimated with *Q* trajectories of the augmented CTMC 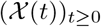 with

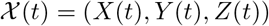

where

- *X*(*t*) = (*X*_1_(*t*), … , *X_d_*(*t*)) is the vector of species copy-numbers.
- 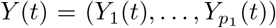 is the vector of additional state-components used for estimating 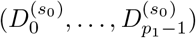 (see Section 4.2).
- *Z*(*t*) = (*Z*_1_(*t*), … , *Z_R_*(*t*)) is the vector of additional state-components used for estimating (*G*(*s*_1_), … , *G*(*s_R_*)) (see Section 4.3).

The augmented process has

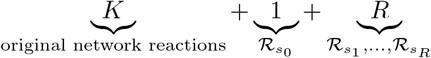

reactions. Note that for each *j* = 0, … , *R*, reaction 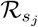 has the constant propensity of λ_*K*+1+*j*_(*x*) ≔ *s_j_*. Our Padé PSD method simulates such a reaction network over the time-interval [0, *T_f_*], by extending the classical Gillespie’s Stochastic Simulation Algorithm [37], and then estimates the Padé derivatives and the direct estimates (*G*(*s*_1_), … , *G*(*s_R_*)). Under this extension, when the firing reaction is *k* = 1, … , *K*, then the state (*x, y, z*) moves to (*x* + ζ_*k*_, *y, z*) as in the original CTMC. However when the firing reaction is *k* = (*K* + 1) then the state (*x, y, z*) moves to (*x, y′, z*) where

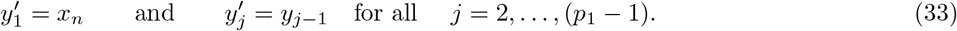

Similarly if the firing reaction is *k* = (*K* + *r*) for some *r* = 1, … , *R* then the state (*x, y, z*) moves to (*x, y, z′*) where

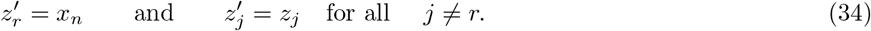

Estimation of the Padé derivatives at ∞ (i.e. 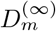 for *m* = 0, … , (*p*_2_ −1)) requires several evaluations of functions of the form 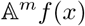. This can be done recursively but it is computationally very intensive. In order to minimise these evaluations we exploit the fact that ergodic Markov chains visit the same set of states again and again. Therefore if we can intelligently store the values 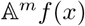 generated by this function, and quickly retrieve them as needed, then it provides a way to leverage the vast memory resources in modern computers in order to gain computational efficiency. Fortunately, `Python` provides an ideal data structure, called a `dictionary`, for this purpose and we use it in our computational implementation to boost the efficiency of Padé PSD.

## 5 Examples

In this section we present several biological examples to illustrate applications of Padé PSD method and also the PSD decomposition result for linear networks (Theorem 3.1). We start by considering some simple linear networks where analytical expressions for the exact PSDs are known and we show that Padé PSD is able to provide very accurate approximations to the PSD (see Section 5.1). Next we discuss how our PSD decomposition result allows us to identify a key criterion that enables differentiation between adapting circuit topologies [33] (see Section 5.2). We then provide two case studies to illustrate the usefulness of our PSD estimation method for synthetic biology applications. In Section 5.3 we examine the problem of optimising the oscillation strength of the *repressilator* [34] and in Section 5.4 we consider the problem of reducing single-cell oscillations that typically arise due to the recently proposed *antithetic integral feedback* (AIF) controller [35] that has the important property of ensuring robust perfect adaptation for arbitrary intracellular networks with stochastic dynamics. In Section 5.5 we examine how the PSD decomposition result can help us in studying the phenomenon of single-cell entrainment in the stochastic setting. Lastly in Section 5.6 we present an example to show how Padé PSD facilitates parameter inference with experimental single-cell trajectories that measure the copy-numbers of the output species up to an unknown constant of proportionality.

Detailed descriptions of the networks considered in the paper and their PSD analysis can be found in Section S4 of the Supplement. All propensity functions are assumed to follow mass-action kinetics (2) unless stated otherwise.

### 5.1 Validation of Padé PSD with linear networks

We now provide analytical expressions for the PSD of certain simple networks, like the birth-death, the classical gene expression network [44] and the recently proposed RNA splicing network [45]. We then show that Padé PSD is able to approximate the PSD quite accurately.

#### Gene Transcription

Consider a simple model of constitutive gene transcription and mRNA degradation, given by a single-species birth-death network with rate of production *k* and the rate of degradation *γ*

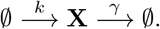

The stationary distribution for this network is Poisson with parameter *k*/*γ*. Hence the stationary mean and variance and equal to *k*/*γ* and applying formula (12) we can compute the PSD as

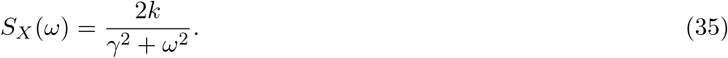

This shows that up to a multiplicative constant, the PSD follows the fat-tailed *Cauchy Distribution* with infinite mean and variance.

#### Gene Expression Network

We now analyse the gene expression model shown in Figure 2(A) that consists of two species - the mRNA (**X**_1_) and the protein (**X**_2_). There are four reactions corresponding to mRNA transcription, protein translation and the first-order degradation of both the species. Observe that the mRNA dynamics is birth-death and hence we can compute its PSD using (35) with (*k*, *γ*) ↦ (*k_r_*, *γ*_*r*_). Since mRNA stimulates the creation of protein via a reaction of the form (13) we can apply our PSD Decomposition result (Theorem 3.1) to express the protein PSD as a sum of two components corresponding to translation and transcription respectively:

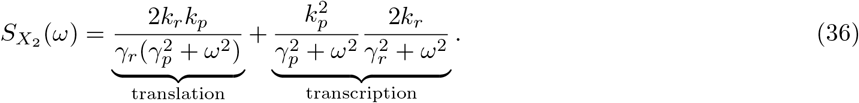

**Figure 2:**
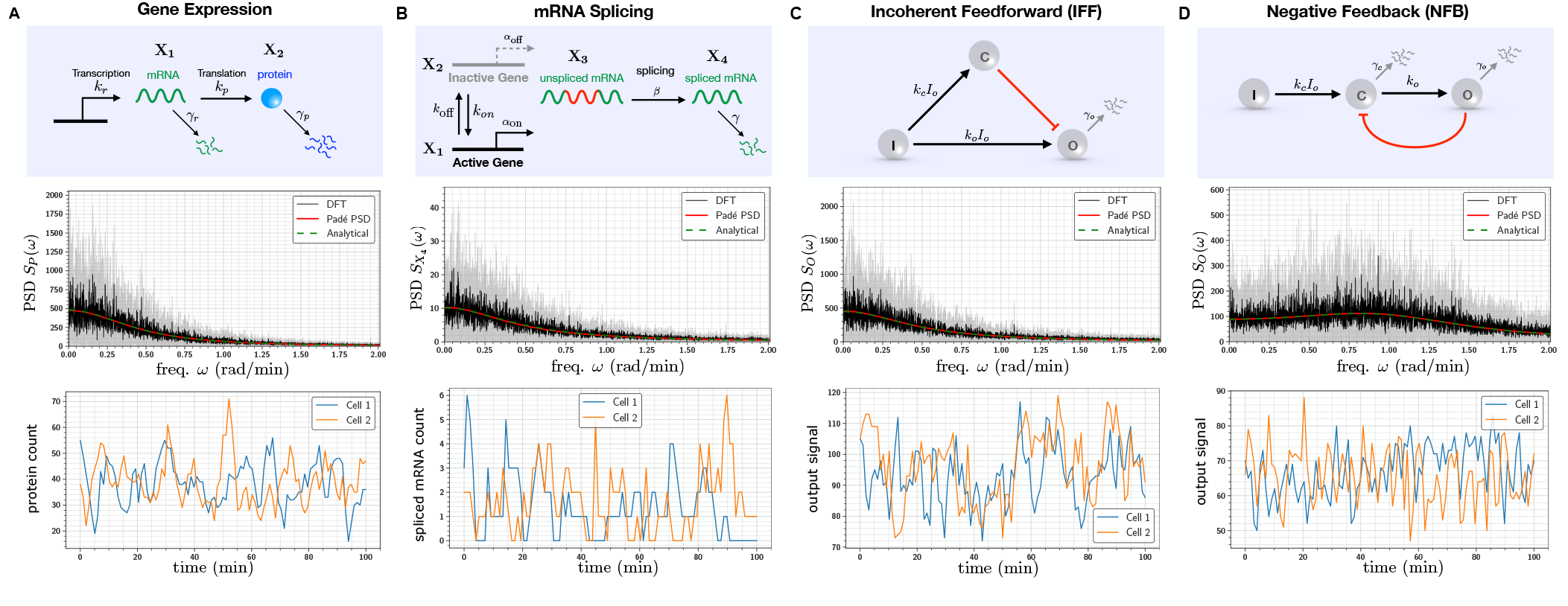
Frequency domain analysis of linear propensity networks: (A) This is the standard gene expression model where mRNA (**X**_1_) is transcribed constitutively and it translates into protein (**X**_2_). (B) In this RNA splicing network a gene can randomly switch between inactive (**X**_2_) (low transcription) and active (**X**_1_) (high transcription) states. When transcription occurs, unspliced mRNA (**X**_3_) is created which is then converted into spliced mRNA (**X**_4_) by the splicing machinery. (C) In the Incoherent Feedforward (IFF) network an input **I**(constant level I0) directly produces the output **O** and it produces the controller species **C**, which represses the production of output **O**. (D) In the Negative Feedback (NFB) network the input **I**(constant level I0) produces the controller species **C**, that produces the output species **O** which in turn inhibits the production of **C** from **I**. For all the networks single-cell output trajectories in the stationary phase are plotted. We provide a comparison of the single-cell PSDs estimated with three approaches - 1) analytically (see Table 1), 2) the Padé PSD method (see Table 1) using ten simulated trajectories and 3) the averaged periodogram or the DFT method mentioned in Box 1 using discrete samples from ten simulated trajectories. For the DFT estimator, the black curve represents the mean of the ten PSDs and the shaded grey region represents the symmetric one standard deviation interval around the mean. For the NFB network one can see that detecting the presence of oscillations in the fluctuations is much easier in the frequency-domain than in the time-domain.

**Table 1:** Expressions for PSDs estimated analytically and with the Padé PSD method. The order of the Padé approximant is **p** = (2, 4) for the RNA splicing network and for all other networks it is **p** = (0, 4).

| Network | Analytical PSD | Padé PSD |
| --- | --- | --- |
| Gene Expression | $\frac{40\omega^2+120}{\omega^4+1.25\omega^2+0.25}$ | $\frac{39.9629\omega^2+119.4918}{\omega^4+1.2368\omega^2+0.2521}$ |
| RNA Splicing | $\frac{1.8\omega^4+36\omega^2+162.234}{\omega^6+20.25\omega^4+69\omega^2+16}$ | $\frac{1.7898\omega^4+146.7412\omega^2+106.7541}{\omega^6+82.2862\omega^4+45.6011\omega^2+11.16}$ |
| IFF | $\frac{94\omega^2+112}{\omega^4+1.25\omega^2+0.25}$ | $\frac{93.988\omega^2+114.53}{\omega^4+1.2752\omega^2+0.2527}$ |
| NFB | $\frac{66.6667\omega^2+200}{\omega^4-0.75\omega^2+2.25}$ | $\frac{66.6892\omega^2+199.7149}{\omega^4-0.754\omega^2+2.2426}$ |

The translation term is computed by setting the mRNA level to its stationary mean 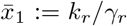 and then viewing the protein dynamics as a birth-death process with production rate 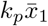 and degradation rate *γ*_*p*_. The transcription term is simply the PSD of mRNA modulated by the frequency dependent factor given by Theorem 3.1.

#### RNA Splicing network

The recently proposed RNA Splicing network (see Figure 2(B)) was used to model the concept of RNA velocity that can help in understanding cellular differentiation from single-cell RNA-sequencing data [45]. Here a single gene-transcript can randomly switch between inactive (**X**_1_) and active (**X**_2_) states with different rates of transcription of unspliced mRNA (**X**_3_). The splicing process converts these unspliced mRNAs into spliced mRNAs (**X**_4_) that can then undergo first-order degradation. Applying formula (12) we can write the PSD of the dynamics of active gene count as

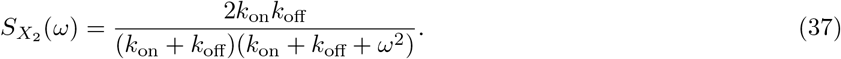

Note that when the active gene count is *X*_2_ ∈ {0, 1} the transcription rate is α_off_ + (α_on_ − α_off_)*X*_2_. We can view transcription as a superposition of two reactions - a constitutive reaction with rate α_off_ and reaction of the form (13) where the stimulant is the active gene **X**_2_. Applying Theorem 3.1 we can decompose the PSD of the spliced mRNA count as

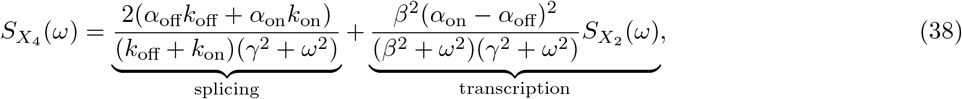

where 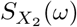 is given by (37).

Observe that for both gene expression and RNA splicing networks we can find an analytical expression for the PSD by directly applying formula (12) for the full network. However using our PSD decomposition result we not only simplify the computation but also identify the contribution of the network mechanisms to the PSD.

For a specific parameterisation of these two networks we compare the PSDs obtained analytically with those obtained by our Padé PSD method described in Section 4 and the standard periodogram estimator for PSD that is based on discrete-sampling and DFT (see Box 1). The results are presented in Figure 2(A-B) and they show good agreement, despite the noisy nature of the DFT estimate. The analytical expressions for the PSD along with the PSD estimates produced by Padé PSD are given in Table 1. One can see that the PSD estimated by our method is quite “close” to the analytical PSD for the gene expression network. The same holds for the RNA splicing network (see the PSD plots in Figure 2(B)) even though it is not apparent from the expressions in Table 1.

### 5.2 Differentiation between adapting regulatory topologies

We consider simple three-node IFF and NFB topologies depicted in Figure 2(C,D) with stochastic kinetics. We provide analytical expressions for the PSDs under the assumption of linearised propensity functions for the repression mechanisms. These expressions inform us about qualitative *structural* differences between the PSDs obtained from IFF and NFB topologies, regardless of the choice of reaction rate parameters. This shows that in the stochastic setting, the PSD of single-cell trajectories serves as a key “response signature” that can differentiate between adapting circuit topologies. We demonstrate this finding with our Padé PSD model for a specific parametrisation of these networks and we argue why this result holds for arbitrarily-sized IFF and NFB networks.

We begin by analysing the IFF topology, where the controller species **C** catalytically produces the output species **O** at rate *F_f_*(*x_c_*) which is a monotonically decreasing function of the controller species copy-number *x_c_* and it represents the repression of **O** by **C**. We linearise the function *F_f_*(*x_c_*) as

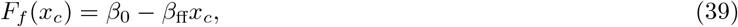

where β_0_ and β_ff_ are positive constants denoting the *basal* production rate and the strength of the incoherent feedforward mechanism respectively. With this linearisation, all propensity functions become affine and hence we can apply the results from Section 3 for linear networks. Specifically the steady-state means 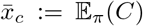 and 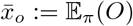 are given by

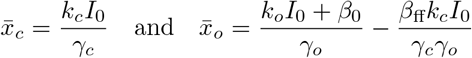

and it is immediate that if β_ff_ ≈ *k_o_*γ_*c*_/*k_c_*, then the mean output value 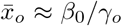 becomes insensitive to the input abundance level *I*_0_. This shows the adaptation property of the IFF network.

As the dynamics of **C** is simply birth-death with production rate *k_c_I*_0_ and degradation rate *γ*_*c*_, its PSD is given by

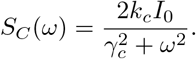

Under the assumption of linearity of the feedforward function *F_f_* the stimulation of **O** by **C** can be viewed as zeroth order degradation. Applying Theorem 3.1 we can evaluate the output PSD as

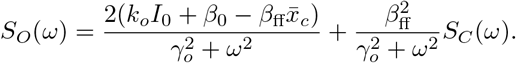

Since this is a sum of two nonnegative monotonically decreasing functions of ω, we can conclude that S*O*(ω) is also monotonically decreasing. Hence output trajectories cannot show oscillations *regardless of the IFF network parameters*. This same argument can be extended to IFF networks with arbitrary number of nodes (see the Supplement, Section S4.1.3).

In the NFB topology, the production of the controller species **C** is repressed by the output species **O**, and we model the production rate by a monotonically decreasing function *F_b_*(*x_o_*) of the output species copy-number *x_o_*. As before, we linearise this function as

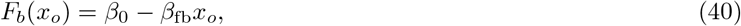

where β_0_ is the basal production rate and β_fb_ is the feedback strength. Under this linearisation, the steady-state means 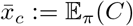 and 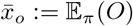 are given by

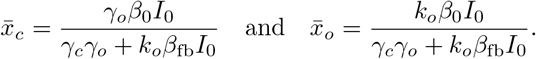

Observe that if the input abundance level *I*_0_ is high, then mean output value 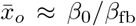 only depends on the feedback function *F_b_* and it is insensitive to *I*_0_, thereby demonstrating the adaptation property. Applying formula (12) we arrive at the following expression for the PSD for the output trajectory

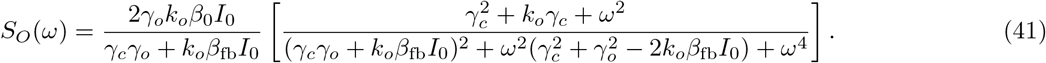

Proposition S4.1 in the Supplement proves that the mapping ω ↦ *S_O_*(ω) has a positive local maximum (which is also the global maximum) if and only if

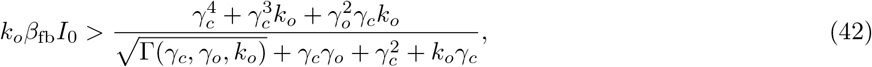

where 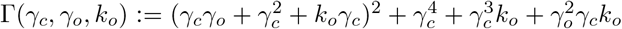. This condition shows that *regardless of the choice of NFB network parameters*, the output trajectories will exhibit oscillation if the input abundance level *I*_0_ is high enough. Using the standard root-locus argument [46] we can draw the same conclusion for arbitrarily-sized NFB networks (see the Supplement, Section S4.1.3). This shows that existence of oscillations and non-monotonicity of the PSD is a differentiator between the NFB and the IFF networks as the latter never exhibits oscillations. Note that high *I*_0_ is precisely the condition for NFB to show adaptation and hence imposing this requirement is not very restrictive. The role of negative feedback in causing stable stochastic oscillations was explored theoretically in [27] with CLE, and it has also been demonstrated experimentally.

For a specific parameterisation of the three node IFF and NFB networks we compare the PSD produced by our method with the analytical PSD^1^ and the DFT-based estimator. The results are shown in Figure 2(C-D) and one can see that Padé PSD is quite accurate in estimating the PSD, which is also evident from the PSD expressions provided in Table 1.

### 5.3 Improving oscillation strength for the *repressilator*

The *repressilator* [34] is the first synthetic genetic oscillator and it consists of three genes repressing each other in a cyclic fashion (see Figure 3(A)). These three genes are *tetR* from the Tn10 transposon, *cI* from bacteriophage λ and *lacI* from the lactose operon. These three genes create three repressor proteins which are TetR, cI and LacI respectively, and the cyclic repression mechanism can be represented as

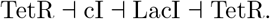

**Figure 3:**
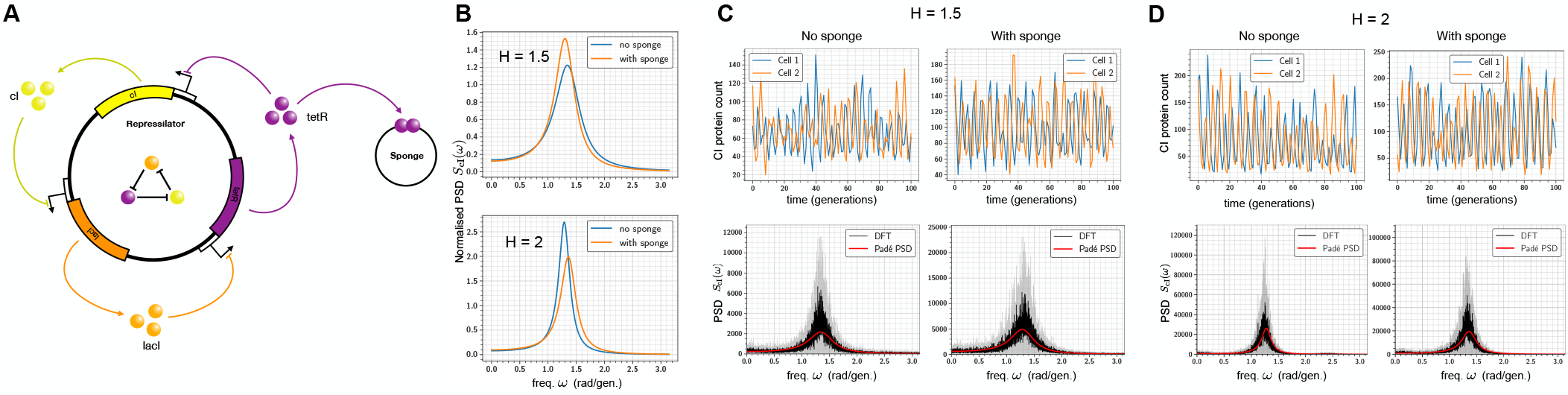
Improving the *repressilator*’s oscillatory strength: (A) Depiction of the *repressilator* network with three gene expression systems whose output proteins cyclically repress each other. When present, the *sponge* plasmid can bind TetR proteins, thereby raising the derepression threshold of the *cI* gene. (B) Shows the effect of the sponge on the normalised PSD obtained by dividing the PSD by the total area under its curve. It can be seen that the sponge sharpens the PSD peak for promoter cooperativity *H* = 1.5 but the opposite occurs for *H* = 2. (C) Plots the single-cell trajectories with and without the sponge for promoter cooperativity *H* = 1.5, and they show that the oscillations are more regular in the latter case. Comparison of the PSD estimated with our Padé PSD method with the PSDs estimated with DFT is provided. (D) Repeats the computational analysis in part (C) for promoter cooperativity *H* = 2.

Due to intrinsic noise in the dynamics, the *repressilator* loses oscillations at the bulk or the population-average level after a few generations. At the single-cell level this intrinsic noise broadens the output PSD peak, making the oscillations less regular in both amplitude and phase. In other words, intrinsic noise compromises the ability of the circuit to *keep track of time*. This issue was addressed in a recent paper [47] which elaborately studied the various sources of noise in the original circuit and eliminated them to construct a modified *repressilator* circuit that showed regular oscillations over several generations. It was found that most of the noise was generated when TetR protein levels were low and the derepression of the TetR controlled promoter occurred at a low threshold. To raise this threshold a *sponge* plasmid was introduced and this had the remarkable effect of regularising the oscillations and sharpening the single-cell PSD peak.

It is also known that increasing the cooperativity of the repression mechanism improves regularity of the oscillations [34]. A fundamental question then arises is that - does the PSD-sharpening effect of the sponge plasmid persist when the repression cooperativity is increased? If this is true then one can regularise oscillations even more by designing cooperative promoters in addition to employing the sponge device. We study this question using an adaptation of the stochastic model given in [47]. The stochastic model is detailed in Section S4.2.1 of the Supplement. The repression mechanism is encoded with a nonlinear Hill function whose coefficient *H* represents the degree of cooperativity among the promoter binding sites. The sponge plasmid, if present, can competitively bind the free TetR molecules, reducing the number of these molecules available for repressing the *cI* gene.

We demonstrate that our method is able to accurately estimate the single-cell PSD and exhibit the sharpening of the PSD in the presence of the sponge plasmid when the cooperativity is set to *H* = 1.5. Surprisingly when the cooperativity is increased to *H* = 2, the sponge has the opposite effect of broadening the PSD. This shows that in certain parameter regimes, the oscillation-regularising effects of the sponge plasmid and the repressor binding cooperativity *are not additive*, possibly due to the fact that increased cooperativity makes the repression mechanism more ultrasensitive [48].

With our method we estimate the PSD for the dynamics of the copy-numbers of the cI protein, whose expression is directly repressed by TetR. For the promoter cooperativity (i.e. the Hill coefficient) of *H* = 1.5, the PSD (area under the PSD curve is normalised to 1) indeed exhibits a sharper peak, in the presence of the sponge plasmid, at the peak frequency of around ω_max_ ≈ 1.35 rad./gen. (see Figure 3(B)). This sharpness in PSD suggests more regularity in oscillations which is also evident from the single-cell trajectories plotted in Figure 3(C). We compare our PSD estimation method with the DFT method in both the cases (with and without sponge) and the results are shown in Figure 3(C). The same analysis is repeated for the promoter cooperativity of *H* = 2 and the results are shown in Figure 3(B, D). From Figure 3(B) it is immediate that for *H* = 2, instead of sharpening the PSD, the addition of the sponge plasmid, actually broadens the PSD slightly.

### 5.4 Reducing single-cell oscillations due to the antithetic integral feedback controller

In recent years genetic engineering has allowed researchers to implement bio-molecular control systems within living cells (see [35, 49, 50, 51, 52, 53, 54, 55, 56, 57, 58]). This area of research, popularly known as *Cybergenetics* [49], offers promise in enabling control of living cells for applications in biotechnology [59, 60] and therapeutics [61]. A particularly important challenge in Cybergenetics is to engineer an intracellular controller that facilitates cellular homeostasis by achieving *robust perfect adaptation* (RPA) for an output state-variable in an arbitrary intracellular stochastic reaction network. This challenge was theoretically addressed in [35] which introduced the *antithetic integral feedback* (AIF) controller and demonstrated its ability to achieve RPA for the population-mean of output species. This controller has been synthetically implemented *in vivo* in bacterial cells, and it has been shown that any bio-molecular controller that achieves RPA for arbitrary reaction networks with noisy dynamics, must embed this controller [58].

Computational analysis has revealed that AIF controller can cause high-amplitude oscillations in the single-cell dynamics in certain parameter regimes [35, 62] which could potentially be undesirable and/or unfavorable. Hence it is important to find ways to augment the AIF controller, so that single-cell oscillations are attenuated but the RPA property is preserved. It is known that adding an extra negative feedback from the output species to the actuated species maintains the RPA property, while decreasing both the output variance and the settling-time for the mean dynamics [63]. Using the PSD estimation method developed in this paper we now demonstrate how adding such a negative feedback also helps in diminishing single-cell oscillations.

The AIF controller is depicted in Figure 4(A) and it is acting on the gene expression model considered in Section 5.1. The AIF controller robustly steers the mean copy-number level of the protein **X**_2_ to the desired set-point *μ/θ*, where *μ* is the production rate of **Z_1_** and *θ* is the reaction rate constant for the output sensing reaction. The AIF affects the output by actuating the production of mRNA **X**_1_ and the feedback loop is closed by the annihilation reaction between **Z_1_** and **Z_2_**. This annihilation reaction can be viewed as mutual inactivation or sequestration and it can be realised using bio-molecular pairs such as sigma/anti-sigma factors [64, 65, 52], scaffold/anti-scaffold proteins [66] or toxin/antitoxin proteins [67].

**Figure 4:**
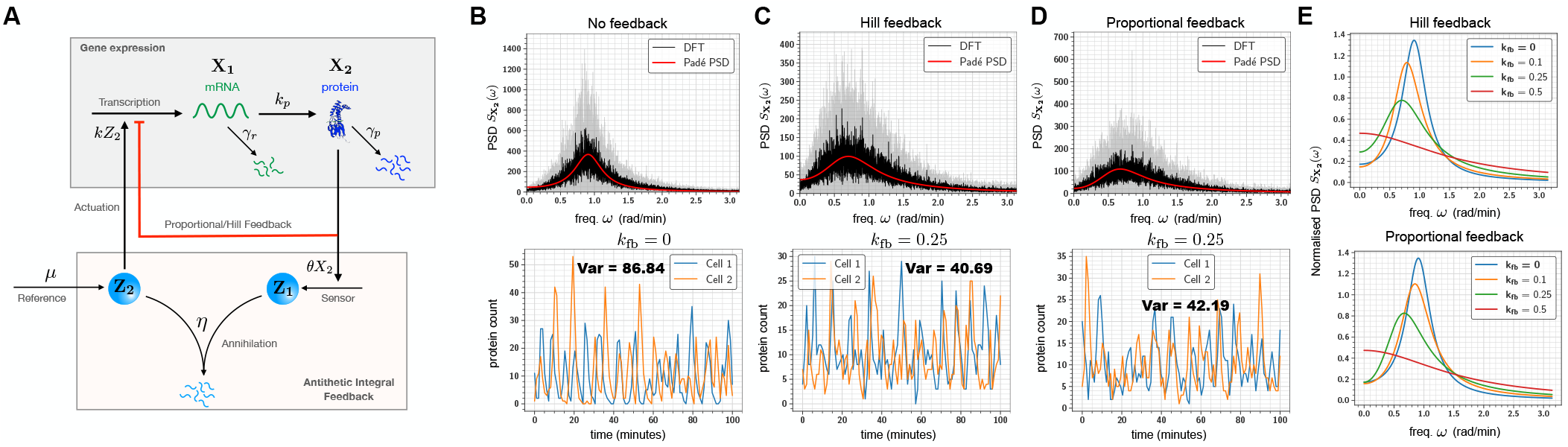
(A) Depiction of the bio-molecular *antithetic integral feedback* (AIF) controller regulating the gene expression network. Here mRNA (**X**_1_) is the actuated species and the protein (**X**_2_) is the output species. This protein output is sensed by the controller species **Z_2_** which annihilates the other controller species **Z_1_** that is constitutively produced at rate *μ*. The species **Z_1_** actuates the gene expression network by catalysing the production of mRNA **X**_1_. The red arrow indicates an extra negative feedback from the output species (protein) to the actuated species (mRNA). In (B) the single-cell oscillatory trajectories for the protein counts (without the extra feedback) are plotted and the corresponding PSD is estimated with Padé PSD and the DFT method. (C) Same plots as in panel (B) for Hill feedback with *k*_fb_ = 0.25 min^*−*1^. (D) Same plots as in panel (B) for proportional feedback with *k*_fb_ = 0.25 min^*−*1^. For other values of *k*_fb_, comparison plots between Padé PSD and DFT are provided in Figure S5 in the Supplement. The plots for the single-cell trajectories in panels (B-D) also indicate the total signal power which is equal to the stationary output variance (see Box 1). Notice the *≥* 50% reduction in this variance in the presence of feedback. (E) Comparison of the normalised PSDs estimated with the Padé PSD method for the Hill and proportional feedback for three choices of feedback parameter *k*_fb_.

It is known from [35] that the combined closed-loop dynamics is ergodic and mean steady-state protein copy-number is *μ/θ*

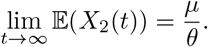

As discussed in [63], this ergodicity is preserved under certain conditions when an extra negative feedback reaction from protein **X**_2_ to mRNA **X**_1_ is added. Letting *z*_1_ and *x*_2_ denote the copy-numbers of **Z_1_** and **X**_2_ respectively, we add the extra feedback by changing the rate of the actuation reaction from *kz*_1_ to (*kz*_1_ + *F_b_*(*x*_2_)) where *F_b_* is a monotonically decreasing feedback function which takes nonnegative values. As in [63] we consider two types of feedback. Letting 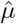 to be the reference point, the first is Hill feedback of the form

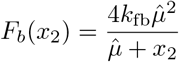

which is based on the actual output copy-number *x*_2_, while the second is the *proportional* feedback that is essentially the linearisation of the Hill feedback at the reference point 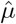

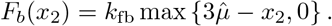

One can easily see that at the reference point, the values of this feedback function 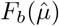 and its derivative 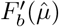 (equal to −*k*_fb_) are the same for both types of feedback. We can view *k*_fb_ as the feedback gain parameter. The Hill feedback is biologically more realisable, while the proportional feedback captures the classical controller where the feedback strength depends linearly on the deviation of the output *x*_2_ from the reference point 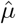, in the output range 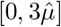. In our analysis we set the reference point 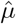 as the set-point *μ/θ*.

For a particular network parametrization we use our method to estimate the PSD for the single-cell protein dynamics in the AIF regulated gene expression network, and the results are displayed in Figure 4. When the extra negative feedback is absent (i.e. *k*_fb_ = 0) the single-cell trajectory has high-amplitude oscillations which is also evident from the estimated PSD (see Figure 4(B)). In Figure 4(C) we apply our Padé PSD method to examine how the PSD changes when extra feedback of Hill type is added with varying strengths given by parameter *k*_fb_. Observe that as the feedback strength increases, the PSD peak declines and the the oscillations become almost non-existent for *k*_fb_ = 0.5 min^*−*1^ (i.e. the PSD becomes monotonic). The same holds true for the proportional feedback (see Figure 4(D)). These results suggest that both feedback mechanisms are more or less equally effective in reducing oscillations. This is further corroborated by the single-cell trajectories plotted in Figure 4(C-D) which also shows that addition of feedback decreases the stationary output variance, that is equal to the signal power (see Box 1). The details on the computations for the AIF regulated gene expression network can be found in Section S4.2.2 of the Supplement.

### 5.5 Stochastic entrainment by a noisy upstream oscillator

The phenomenon of entrainment occurs when an oscillator, upon stimulation by a periodic input, loses its natural frequency and adopts the frequency of the input. This phenomenon has several applications in physical, engineering and biological systems [68]. The most well-known biological example of this phenomenon is the entrainment of the circadian clock oscillator by day-night cycles. The circadian clock is an organism’s time-keeping device and its entrainment is necessary to robustly maintain its periodic rhythm [69]. The circadian clock is one example among several intracellular oscillators that have been found and their functional roles have been identified [70]. Often these oscillators provide entrainment cues to other networks within cells [71] and hence it is important to study entrainment at the single-cell level, where the dynamics is intrinsically noisy due to low copy-number effects.

We now illustrate how our PSD decomposition result (Theorem 3.1) can be used to study single-cell entrainment in the stochastic setting where the dynamics is described by CTMCs. We consider the example of the *repressilator* stimulating a gene expression system, as shown in Figure 5(A). This gene expression network is the same as in Section 5.1 but we include transcriptional feedback from the protein molecules and so the mRNA transcription rate is given by a monotonic decreasing function *F_b_*(*x*_2_) of the protein copy-number *x*_2_. We shall linearize *F_b_*(*x*_2_) as

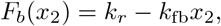

where *k_r_* is the basal transcription rate and *k*_fb_ is the feedback strength. When this gene expression network is connected to the *repressilator* (described later in Section 5.3) the transcription rate changes from *F_b_*(*x*_2_) to

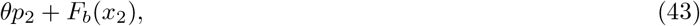

where *p*_2_ is the molecular count of protein cI in the *repressilator* and parameter *θ* captures the “strength” of the interconnection. In other words, cI acts as an activating transcription factor in our example. The parameters of the *repressilator* are chosen as in Section 5.3 in the “no sponge” and Hill coefficient *H* = 1.5 case, but the time-units are changed to minutes. We can view the gene expression network as simply the negative feedback (NFB) network from Section 5.2 with the controller species **C** as mRNA **X**_1_ and the output species **O** as protein **X**_2_. Using the same parameters as the NFB network, we study how the PSD of the protein output varies as a function of *θ*. In order for the gene expression network to be entrained to the *repressilator* the global maxima of this protein PSD should be near the *repressilator’s* natural (or peak) frequency of about 1.35 rad./min (see Figure 3(C)).

**Figure 5:**
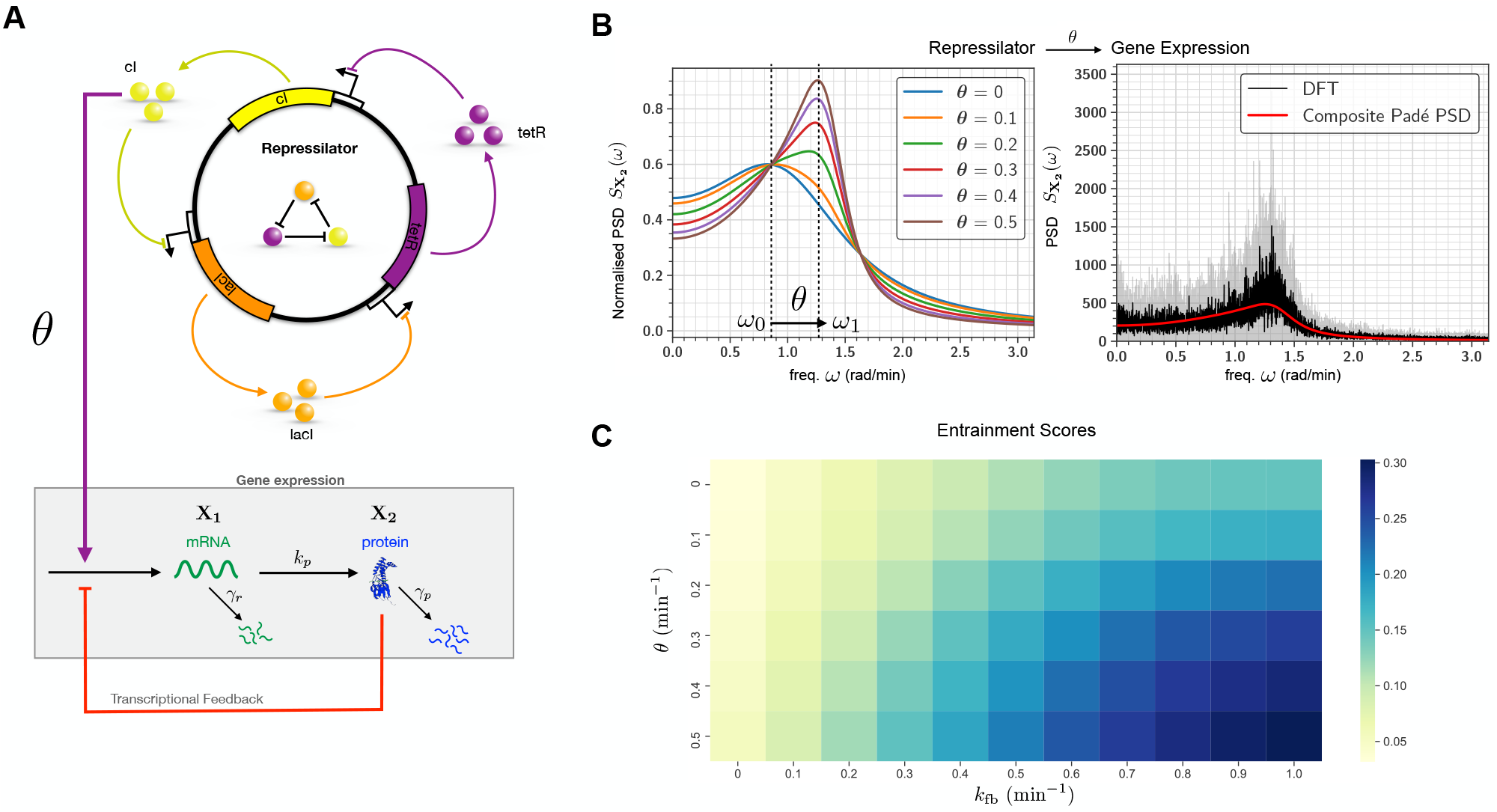
Stochastic entrainment of gene expression by the *repressilator*: (A) Schematic diagram of the *repressilator* driving a gene expression network. The cI protein from the *repressilator* acts as an activating transcription factor for mRNA (**X**_1_) which translates into output protein (**X**_2_). The red arrow from **X**_2_ to **X**_1_ indicates negative transcriptional feedback from the protein molecules. When the *repressilator* is connected to the gene expression network, the PSD can be estimated with the *composite* Padé PSD method which is based on Theorem 3.1. In (B) these PSD estimates (after normalisation) are plotted for six values of *θ* and compared for *θ* = 0.4 min^*−*1^ to the PSD obtained with the DFT method. One can observe the *stochastic entrainment* phenomenon as *θ* increases. (D) The heat-map for the entrainment score (see (45)) as a function of *θ* and the feedback strength parameter *k*_fb_. Observe that the entrainment score is monotonically increasing in both variables *k*_fb_ and *θ*, but it is more sensitive to *k*_fb_.

To compute the PSD of the combined network we shall apply Theorem 3.1. For this we first consider the gene expression network in isolation with *p*_2_ in the transcription rate (43) replaced by the constant steady-state mean of *p*_2_ (denoted by 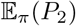). Hence using (41) we can estimate the protein dynamics PSD 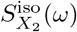 as

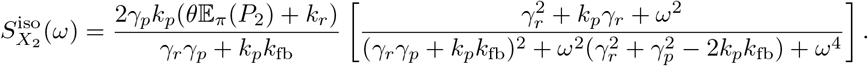

Irrespective of the value of *θ*, the PSD 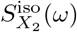 has a global maxima at ω_max_ ≈ 0.85 rad./min. which is the natural frequency of the gene expression circuit in isolation.

When the *repressilator* is connected to the gene expression network, we can apply Theorem 3.1 to compute the PSD of the protein output as

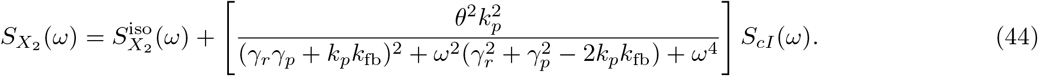

We call this method *composite* Padé PSD as it estimates the PSD for the full network by combining two approaches - Padé PSD for the nonlinear subnetwork (*repressilator*) with the analytical expression for the linear subnetwork (gene-expression). In Figure 5(B) we plot the normalised PSD (area under the PSD curve is normalised to 1) for six values of *θ* and we also validate this composite method with the DFT method for *θ* = 0.4 min^*−*1^. One can clearly see that as *θ* gets higher, the gene expression network gives up its natural frequency upon stimulation and adopts a frequency which is close to the *repressilator* frequency. This exemplifies the phenomenon of single-cell entrainment in the stochastic setting.

In order to investigate this entrainment phenomenon further we define an *entrainment score* as

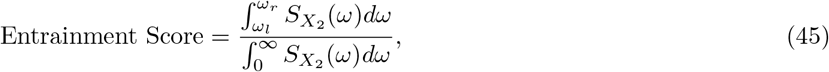

where [ω_*l*_, ω_*r*_] = [0.9ω_0_, 1.1ω_0_] represents an interval of relative length 10% on either side of the *repressilator’s* natural frequency ω_0_. In Figure 5(C) we plot a heat-map for the entrainment score as a function of the feedback strength parameter *k*_fb_ and the connection strength parameter *θ*. One can see that the entrainment score increases monotonically with *θ* which is to be expected as the first term on the r.h.s. of (44) scales linearly with *θ* while the second term scales quadratically. Similarly by computing the ratio of the two terms we can conclude that entrainment score is also a monotonically increasing function of *k*_fb_. However as the heat-map clearly indicates, the entrainment score is more sensitive to *k*_fb_ than *θ*, thereby suggesting that transcriptional feedback could be a critical mechanism for facilitating entrainment of gene expression networks.

### 5.6 PSD as a tool for parameter inference

Consider a self-regulatory gene expression system (see Figure 6(A)) modelled as a simple birth-death network where the production rate is given by the repressing Hill function

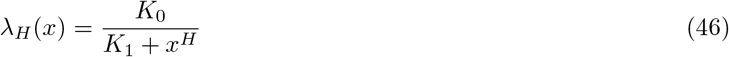

of the output copy-number x and the degradation rate is *γ*. Fixing all other parameters, our goal is to use the experimental PSD to infer the degree of cooperativity *H*. This experimental PSD is generated via simulations with *H* = 1 and we average the PSDs over 100 single-cell trajectories in order to reduce the variance in the DFT-based PSD estimate. We assume that the experimental single-cell trajectories are proportional to the output copy-number but the constant of proportionality is *unknown* as is often the case in time-lapse microscopy experiments. We also assume that there is no measurement noise - if the measurement noise appears as an independent process then its PSD simply appears as an additive term in the output PSD, which can be easily removed to recover the output PSD without the measurement noise.

**Figure 6:**
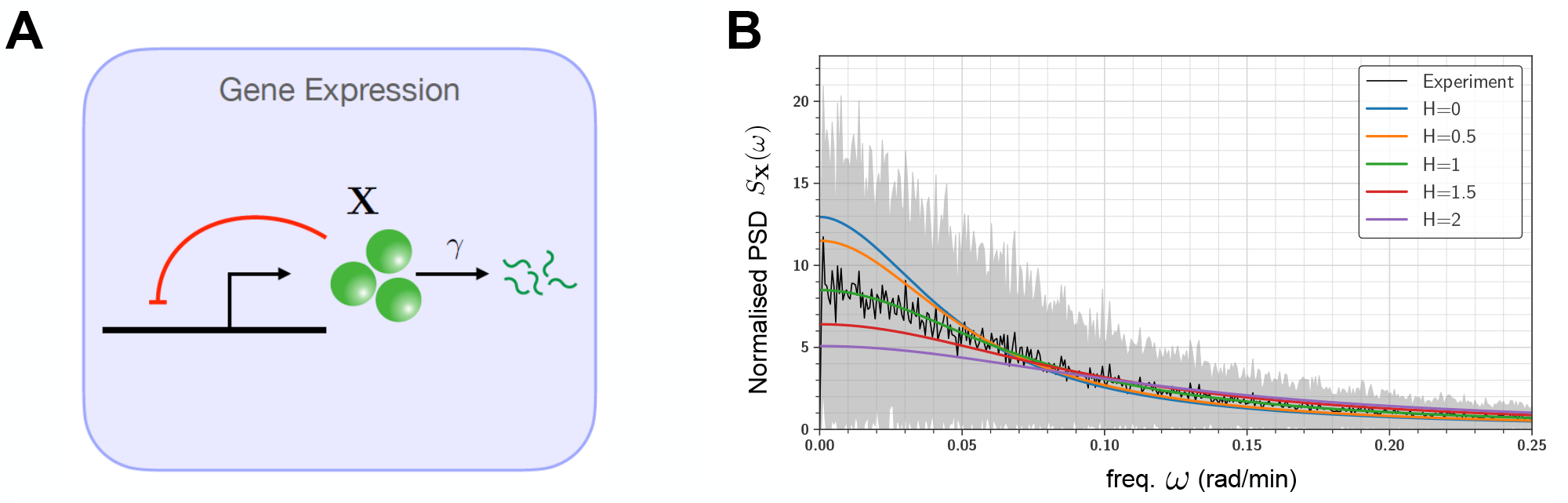
PSD-based inference of a self-regulatory gene expression model: (A) Depicts the self-regulatory gene expression system where the output represses the gene (shown in red) via a nonlinear Hill function (46) with cooperativity coefficient *H*. (B) Plots the normalised PSDs obtained by Padé PSD for various values of *H*, and compares it with the normalised PSD obtained from experimental single-cell trajectories. The experimental PSD was computed by averaging the PSDs from 100 single-cell traces, and the black curve represents the mean while the shaded grey region represents the symmetric one standard deviation interval around the mean.

Observe that the unknown constant of proportionality drops out when we compute the normalised PSD (i.e. area under the PSD curve is normalised to 1). Hence we can infer the unknown parameter *H* by estimating the normalised PSD with our Padé PSD method and comparing it with the experimentally obtained normalised PSD. This comparison is performed for various values of *H* in Figure 6(B) and it is evident that the experimental traces come from the network with *H* = 1. Note that the clean estimates for the normalised PSD produced by our Padé PSD method, greatly facilitate the inference of *H*. If the same estimates were obtained with DFT then the estimator noise would obfuscate the dependence of the PSD on *H* and make the inference task difficult.

## 6 Discussion

Recent advances in microscopic imaging and fluorescent reporter technologies have enabled high-resolution monitoring of processes within living cells [1]. As the accessibility of this time-course data rapidly increases, there is an urgent need to design novel theoretical and computational approaches that make use of the full scope of such data, in order to understand intracellular processes and design effective synthetic circuits. An important feature of time-course measurements, which is lacking in the data generated by the more common experimental technique of Flow-Cytometry, is that they capture temporal correlations at the single-cell level which are rich in information about the underlying dynamical model. Frequency domain analysis provides a viable approach to extract this information, if we have an efficient framework to connect network models to the frequency spectrum or the power spectral density (PSD) of the single-cell trajectories measured with time-lapse microscopy [20, 18]. The dynamics within cells is invariably stochastic, owing to the presence of many low abundance biomolecular species, and it is commonly described as a continuous-time Markov chain (CTMC). In this context, the aim of this paper is to develop a computational method for reliably estimating the PSD for single-cell trajectories from CTMC models. Existing approaches for PSD estimation for stochastic network models, are either applicable to a particular class of networks [26, 17], or they are based on dynamical approximations that are known to be inaccurate over large time-intervals and in situations where low abundance species are present [20, 19]. The method we develop in this paper, called Padé PSD, especially pertains to the low abundance regime. It applies generically to any stable network and it yields an accurate PSD expression using a small number of CTMC trajectory simulations. Moreover for networks with affine propensity functions we provide a novel PSD decomposition result that expresses the output PSD in terms of its constituent parts.

The tools we develop in this paper are of significance to both systems and synthetic biology. We demonstrate that in the presence of intrinsic noise, PSD estimation can successfully differentiate between adapting Incoherent Feedforward (IFF) and Negative Feedback (NFB) topologies [33], and it can facilitate performance optimisation of synthetic oscillators [34] as well as synthetic *in vivo* controllers [35]. Moreover it can also aid the study of stochastic entrainment at the single-cell level. This is of particular relevance for applications such as designing pulsatile dynamics of transcription factors, which is known to enable graded multi-gene regulation [72]. Typically experimental single-cell trajectories measure an output species up to a constant of proportionality which is often poorly characterised, causing problems in parameter inference from experimental data. We present a simple example to illustrate that the use of PSDs can bypass this issue and our Padé PSD method can play a useful role in parameter inference.

The main contribution of this paper is to show how the theory of Padé approximations can be effectively applied to the PSD estimation problem for reaction networks with stochastic CTMC dynamics. In Padé PSD a low dimensional approximation of the PSD is computed based on estimates of Padé derivatives that are expressible as certain stationary expectations for which efficient Monte Carlo estimators were developed. These ideas can be combined with several existing techniques to significantly improve the efficiency of our PSD estimation method. Specifically the problem of reliably estimating expectations under the CTMC model has received a lot of attention in recent years [73], and various methods designed for this problem, like τ-leaping [74] and/or multilevel schemes [75], can be easily integrated with Padé PSD, in order to speed up the estimation process and also to reduce the variance of the Monte Carlo estimators. Moreover model reductions [76, 77] and simulation tools [78, 79] for multiscale networks can be readily applied to simplify the estimation of Padé derivatives. Such extensions would greatly expand the scope of applicability of our method and pave the way for frequency-based analysis and design of stochastic biomolecular reaction networks.

## Acknowledgments

This project has received funding from the European Research Council (ERC) under the European Union’s Horizon 2020 research and innovation programme grant agreement no. 743269 (CyberGenetics project).

## Footnotes

1 Since negative propensities cannot be allowed, we perform simulations with the positive part of the linear feedforward (see (39)) and feedback (see (40)) functions. Hence the analytical PSD expressions for the IFF and the NFB networks are not exact.

