## Supplement for "Frequency Spectra and the Color of Cellular Noise"

### Supplementary Text

**S1 Preliminaries** **3**
S2 Expressions for the Padé approximants and the PSDs estimated with the Padé PSD method for the antithetic integral feedback controller. The estimated Padé approximants were validated and the results are shown in Figure S4. . 42
S3 Expressions for the Padé approximants and the PSDs estimated with the Padé PSD method for the self-regulated gene-expression network. The estimated Padé approximants were validated and the results are shown in Figure S6. . 42

### Supplementary Text

#### S1 Preliminaries

It is now well-established that intracellular dynamics is often very noisy due to the stochastic nature of reactions that occur within the small spatial unit of a cell. Many biomolecular species are present in low copy-numbers (i.e. molecular counts) within cells [7, 17] and reactions involving them are a significant source of dynamic heterogeneity and cell-to-cell variation [24]. To capture these heterogeneities and accurately model the dynamics, stochastic models are necessary and we begin this section by describing how stochastic models of reaction networks are constructed. We then review some important concepts from Markov process theory that will help us in developing our method for spectral analysis of single-cell trajectories generated by such stochastic models of intracellular dynamics.

**Notation:** Throughout this text we denote the sets of all reals as $\mathbb{R}$, all complex numbers as $\mathbb{C}$, all nonnegative reals as $\mathbb{R}_+$, all integers as $\mathbb{Z}$ and all nonnegative integers as $\mathbb{Z}_+$. For any set $\mathcal{E} \subset \mathbb{Z}_+^d$, a function $f : \mathcal{E} \to \mathbb{R}$ is called *polynomially growing*, if there exist constants $C, r \geq 0$ such that

$$|f(x)| \leq C(1 + \|x\|^r) \qquad \text{for each} \qquad x \in \mathcal{E},$$

where $\| \cdot \|$ denotes the standard norm on $\mathbb{R}^d$. Note that if $r = 0$ then this function $f$ is simply a bounded function. We denote by $\mathcal{B}(\mathcal{E})$ and $\mathcal{B}_b(\mathcal{E})$ the set of real-valued functions on $\mathcal{E}$ which are polynomially growing and bounded respectively. We refer to the expectation of a function $f : \mathcal{E} \to \mathbb{R}$ under some probability distribution $\mu$ on $\mathcal{E}$ as

$$\mathbb{E}_\mu(f) := \sum_{x \in \mathcal{E}} f(x)\mu(x).$$

When $f(x) = x_m$ then we may write $\mathbb{E}_\pi(f)$ as $\mathbb{E}_\pi(X_m)$.

##### S1.1 Definition of stochastic reaction networks

Suppose we have a reaction network with $d$ species, called $\mathbf{X}_1, \dots, \mathbf{X}_d$, and $K$ reactions of the form

$$\sum_{i=1}^{d} \nu_{ik} \mathbf{X}_i \longrightarrow \sum_{i=1}^{d} \nu'_{ik} \mathbf{X}_i, \qquad k = 1, \dots, K \tag{S1}$$

with $\nu_{ik}$ and $\nu'_{ik}$ being nonnegative integers denoting the number of molecules of species $\mathbf{X}_i$ that are consumed and produced by the $k$-th reaction. In the classical stochastic reaction network model, the dynamics is described as a *continuous-time Markov chain* (CTMC) [1] whose states represent the copy numbers of the $d$ network species. If at some time, the state is $x = (x_1, \dots, x_d) \in \mathbb{Z}_+^d$ and reaction $k$ fires, then the state is displaced by the *stoichiometric* vector $\zeta_k = (\nu'_{1k} - \nu_{1k}, \dots, \nu'_{dk} - \nu_{dk}) \in \mathbb{Z}^d$. Associated with each reaction $k$ is the *propensity* *function* $\lambda_k : \mathbb{Z}_+^d \to \mathbb{R}_+$ which says that at state $x$ this reaction fires at rate $\lambda_k(x)$. Often *mass action kinetics* is assumed, where each $\lambda_k$ is given by

$$\lambda_k(x_1, \ldots, x_d) = \theta_k \prod_{i=1}^{d} \frac{x_i(x_i - 1) \ldots (x_i - \nu_{ik} + 1)}{\nu_{ik}!}, \tag{S2}$$

with the positive parameter $\theta_k$ being the associated rate constant. Our method does not rely on mass-action assumption but we do assume that all propensity functions belong to $\mathcal{B}(\mathcal{E})$, i.e. they are polynomially growing.

We model the reaction dynamics as a CTMC which at any state $x$ does one of two things. Either $x$ is an *absorbing state* (when $\lambda_0(x) := \sum_{k=1}^{K} \lambda_k(x) = 0$) and the CTMC stays at $x$ for all times, or $x$ is not an absorbing state and the CTMC jumps from state $x$ after a random waiting time which is exponentially distributed with rate $\lambda_0(x)$, and this jump is in direction $\zeta_k$ with probability $\lambda_k(x)/\lambda_0(x)$. Formally this CTMC can be specified by its generator which is an operator which specifies the rate of change of the probability distribution of the process (see Chapter 4 in [9]). This generator $\mathbb{A}$ is given by

$$\mathbb{A}f(x) = \sum_{k=1}^{K} \lambda_k(x) \left( f(x + \zeta_k) - f(x) \right), \tag{S3}$$

for any $f \in \mathcal{B}_b(\mathbb{Z}_+^d)$.

We assume that this CTMC evolves on some nonempty state-space $\mathcal{E} \subset \mathbb{Z}_+^d$. This means that for any state $x \in \mathcal{E}$ if there is a positive probability of reaction $k$ firing (i.e. $\lambda_k(x) > 0$) then the resulting state $(x + \zeta_k)$ must also be in $\mathcal{E}$. We call $\mathcal{E}$ as the state space for the CTMC. Let $(X(t))_{t \geq 0}$ be the CTMC with generator $\mathbb{A}$ and an initial state $X(0) \in \mathcal{E}$. For any state $x \in \mathcal{E}$, let

$$p(t, x) = \mathbb{P}\left(X(t) = x\right) \tag{S4}$$

be the probability that the CTMC is in state $x$ at time $t$. These probabilities evolve in time according to the Chemical Master Equation (CME) given by

$$\frac{dp(t,x)}{dt} = \sum_{k=1}^{K} p(t, x - \zeta_k)\lambda_k(x - \zeta_k) - p(t, x) \sum_{k=1}^{K} \lambda_k(x), \tag{S5}$$

for each $x \in \mathcal{E}$. Note that this system has as many ODEs as the number of elements in the state-space $\mathcal{E}$, which is generally infinite or very large. Due to this reason, typically, the CME cannot be directly solved but its solutions can be estimated from Monte Carlo simulations of the sample paths of the CTMC $(X(t))_{t \geq 0}$. The most well-known method for realising these sample-paths is Gillespie's *stochastic simulation algorithm* (SSA) [11].

We end this subsection with a useful *product-rule* for generator $\mathbb{A}$ defined by (S3). For any two functions $g, h : \mathcal{E} \to \mathbb{R}$ the action of generator $\mathbb{A}$ on the product function $g(x)h(x)$ is given by

$$\mathbb{A}\left(g(x)h(x)\right) = g(x)\mathbb{A}h(x) + h(x)\mathbb{A}g(x) + \sum_{k=1}^{K} \lambda_k(x)\Delta_k g(x)\Delta_k h(x), \tag{S6}$$

where $\Delta_k$ is the difference operator

$$\Delta_k g(x) = g(x + \zeta_k) - g(x).$$

This follows by simply applying the generator to the product function $w$ to obtain

$$\mathbb{A}w(x) = \sum_{k=1}^{K} \lambda_k(x) \left( g(x+\zeta_k)h(x+\zeta_k) - g(x)h(x) \right),$$

and then expanding the quantity inside the parenthesis using the identity $(a_1 b_1 - a_2 b_2) = a_2(b_1 - b_2) + b_2(a_1 - a_2) + (a_1 - a_2)(b_1 - b_2)$.

#### S1.2 Steady-state behaviour

In this paper we are interested in the properties of the system after it has settled down, or in other words, the CTMC $(X(t))_{t \geq 0}$ has reached a *steady-state* which is characterised by a stationary distribution $\pi$ that is essentially a fixed-point for the CME (S5). The CTMC is said to be *ergodic* if this fixed-point is unique and globally attracting in the sense that for any initial probability distribution $p(0)$ over the state-space $\mathcal{E}$, the CME solution $p(t)$ satisfies

$$\lim_{t \to \infty} \|p(t) - \pi\|_{\ell_1} = 0, \tag{S7}$$

where $\|p(t) - \pi\|_{\ell_1} = \sum_{x \in \mathcal{E}} |p(t,x) - \pi(x)|$ denotes the $\ell_1$-distance between probability measures $p(t)$ and $\pi$. If the convergence in (S7) is exponentially fast then the CTMC is called *exponentially ergodic*. More precisely, for an exponentially ergodic CTMC there must exist constants $C, \rho > 0$, with $\rho$ independent of the initial probability distribution $p(0)$, such that

$$\|p(t) - \pi\|_{\ell_1} \leq C e^{-\rho t}. \tag{S8}$$

Checking ergodicity for a CTMC arising from reaction networks is a difficult problem because for most networks of interest the state-space $\mathcal{E}$ is infinite. Fortunately there exists a two-step computational procedure for verifying ergodicity for reaction networks that commonly arise in systems and synthetic biology. First, using the techniques in [13] we restrict ourself to a state-space $\mathcal{E}$ which is *irreducible*, in the sense that there is a positive probability of reaching any state in $\mathcal{E}$ from any other state in $\mathcal{E}$ via a sequence of reactions. Second, following the approach in [19], we construct a norm-like[1] Foster-Lyapunov function $V : \mathcal{E} \to [1, \infty)$ such that for some $C_1, C_2 > 0$, we have

$$\mathbb{A}V(x) \leq C_1 - C_2 V(x) \qquad \text{for all} \qquad x \in \mathcal{E}, \tag{S9}$$

where $\mathbb{A}$ is the generator of the CTMC. A systematic procedure based on linear programming is provided in [12] to construct a linear Foster-Lyapunov function of the form

$$V(x) = 1 + \langle v, x \rangle \tag{S10}$$

[1] A positive function is called norm-like if all its sub-level sets are compact.

that satisfies (S9). Here $\langle \cdot, \cdot \rangle$ is the standard inner product and $v$ is vector with positive components. Existence of such a Foster-Lyapunov function satisfying (S9) is enough to guarantee exponential ergodicity (see Proposition 4 in [12]), but if we need to ensure that all moments of the stationary distribution $\pi$ are finite (see Theorem 5 in [12]) then we need to additionally assume that for some $C_3, C_4 > 0$

$$\mathbb{A}V^2(x) \leq 2V(x)\mathbb{A}V(x) + C_3 + C_4 V(x) \qquad \text{for all} \qquad x \in \mathcal{E}. \tag{S11}$$

We now state the main assumptions we make to develop our method for PSD estimation.

**Assumption S1.1** *(A) The state-space $\mathcal{E}$ is irreducible for the CMTC with generator $\mathbb{A}$ modelling the stochastic reaction network.*

*(B) The CTMC is exponentially ergodic which is certified by the existence of a linear Foster-Lyapunov function $V$ satisfying* (S9) *and* (S11)*.*

It can be concluded from Echeverria's Theorem (see Theorem 9.17 in [9]) that under Assumption S1.1, the stationary expectation is zero for any function in the range of the generator $\mathbb{A}$

$$\mathbb{E}_\pi(\mathbb{A}f) = \sum_{x \in \mathcal{E}} \mathbb{A}f(x)\pi(x) = 0. \tag{S12}$$

Under our assumption of exponential ergodicity, relation (S12) will hold for any polynomially growing function $f$ (i.e. $f \in \mathcal{B}(\mathcal{E})$). For such a function $f$ it also holds that the long-term average of this function along a trajectory is almost surely equal to the population or ensemble average under the stationary distribution

$$\lim_{T \to \infty} \frac{1}{T} \int_0^T f(X(t))dt \stackrel{\text{a.s.}}{=} \mathbb{E}_\pi(f). \tag{S13}$$

This classical result is called Birkhoff's Ergodic Theorem (see [20]) and it implies that all the information about the stationary distribution in contained in a single infinitely long trajectory. One can use this result to estimate the stationary expectation $\mathbb{E}_\pi(f)$ using a single trajectory simulated over a time-interval $[0, T]$ as

$$\mathbb{E}_\pi(f) \approx \frac{1}{T} \int_0^T f(X(t))dt. \tag{S14}$$

The r.h.s. is a random variable which would be close to $\mathbb{E}_\pi(f)$ for large enough $T$. One question that naturally arises is that - how large must $T$ be is order to obtain an accurate estimate of $\mathbb{E}_\pi(f)$? This question is answered by the *Central Limit Theorem* for Markov chains (see [2]) that says that the convergence of this estimator with time $T$ is captured by the *Time Average Variance Constant* (TAVC) defined by

$$\text{TAVC}(f) = 2 \int_0^\infty \mathbb{E}_\pi \left[ (f(X(0)) - \mathbb{E}_\pi(f))(f(X(t)) - \mathbb{E}_\pi(f)) \right] dt. \tag{S15}$$

#### S1.3 Spectral density and the Wiener-Khinchine Theorem

Consider the situation where the dynamics of one of the species $\mathbf{X}_n$ is observed under a microscope. We are interested in estimating the strengths of oscillatory components of various frequencies in the output signal $(X_n(t))_{t\geq 0}$. As $\mathbb{E}_\pi(X_n)$ is the stationary mean, we first construct the signal with zero stationary mean as

$$\widetilde{X}_n(t) = X_n(t) - \mathbb{E}_\pi(X_n).$$

A direct implication of (S13) is that the stationary variance of the signal

$$\mathrm{Var}_\pi(X_n) = \mathbb{E}_\pi(X_n^2) - (\mathbb{E}_\pi(X_n))^2$$

is equal to the time-averaged signal power computed as

$$P(X_n) = \lim_{T\to\infty} \frac{1}{T}\int_0^T \left(\widetilde{X}_n(t)\right)^2 dt. \tag{S16}$$

We now define the truncated one-sided Fourier Transform of the signal $(\widetilde{X}_n(t))_{t\geq 0}$ as

$$\mathcal{F}_T(\omega) = \frac{1}{\sqrt{T}}\int_0^T \widetilde{X}_n(t)e^{-i\omega t}dt,$$

where $\omega$ is the frequency and $i = \sqrt{-1}$. The *power spectral density (PSD)* for this output signal is defined by

$$S_{X_n}(\omega) = \lim_{T\to\infty} \mathbb{E}\left(|\mathcal{F}_T(\omega)|^2\right).$$

Since our process is ergodic, we can drop the expectation and estimate $S_{X_n}(\omega)$ from a single trajectory as

$$S_{X_n}(\omega) = \lim_{T\to\infty} |\mathcal{F}_T(\omega)|^2. \tag{S17}$$

Intuitively $S_{X_n}(\omega)$ captures the strength of the oscillatory component with frequency $\omega$ in the mean-zero signal $(\widetilde{X}_n(t))_{t\geq 0}$. This PSD is related to the *autocovariance*[2] function

$$\mathrm{ACV}_{X_n}(\tau) := \mathbb{E}\left[\widetilde{X}_n(t)\widetilde{X}_n(t+\tau)\right] \tag{S18}$$

which can be computed from a single infinitely-long trajectory as

$$\mathrm{ACV}_{X_n}(\tau) := \lim_{T\to\infty}\frac{1}{T}\int_0^T \widetilde{X}_n(t)\widetilde{X}_n(t+\tau)dt. \tag{S19}$$

The well-known *Wiener-Khintchine Theorem* [16] shows that the PSD can be written as the two-sided Fourier Transform of the autocovariance function

$$S_{X_n}(\omega) = \int_{-\infty}^{\infty} \mathrm{ACV}_{X_n}(\tau)e^{-i\omega\tau}d\tau. \tag{S20}$$

[2]The autocovariance function is denoted by R in the main text, but we shall denote it by ACV in this Supplement.

Due to ergodicity, the autocovariance function, as computed by (S19) does not depend on the specific realisation of the stochastic output trajectory $(\widetilde{X}_n(t))_{t\geq 0}$. Hence a simple consequence of (S20) is that the PSD is identical for each stochastic realisation and it only depends on the reaction network model that generates the output.

Taking the inverse Fourier Transform and using the fact that PSD is an even function (i.e. $S_{X_n}(\omega) = S_{X_n}(-\omega)$) we get

$$\text{ACV}_{X_n}(\tau) = \frac{1}{\pi}\int_0^\infty S_{X_n}(\omega)e^{i\omega\tau}d\omega.$$

In particular for $\tau = 0$ we obtain Parseval's Theorem which expresses the stationary output variance $\text{Var}_\pi(X_n)$ or the total power $P(X_n)$ as

$$\text{Var}_\pi(X_n) = P(X_n) = \frac{1}{\pi}\int_0^\infty S_{X_n}(\omega)d\omega. \tag{S21}$$

Therefore the total signal power is simply the integral of the strengths of the oscillatory components for each frequency $\omega$, divided by $\pi$. In this paper we shall at times work with the *normalized PSD* $\overline{S}_{X_n}(\omega)$ obtained by

$$\overline{S}_{X_n}(\omega) = \frac{S_{X_n}(\omega)}{P(X_n)\pi}, \tag{S22}$$

so that the area under the PSD curve is one. It is immediate from (S20) that this normalised PSD is the Fourier Transform of the *autocorrelation* function

$$\text{ACF}_{X_n}(\tau) := \frac{\text{ACV}_{X_n}(\tau)}{\text{Var}_\pi(X_n)} \tag{S23}$$

divided by $\pi$. In other words

$$\overline{S}_{X_n}(\omega) = \frac{1}{\pi}\int_{-\infty}^\infty \text{ACF}_{X_n}(\tau)e^{-i\omega\tau}d\tau. \tag{S24}$$

#### S1.4 Numerical estimation of the PSD from empirical data

In this section we briefly discuss how the PSD for an experimentally observed output trajectory can be numerically estimated. Essentially there are two approaches. The first is the *periodogram* approach based on formula (S17) and it relies on computing the *Discrete Fourier Transform (DFT)* of the output signal trajectory. The second approach is based on the *Wiener-Khintchine* representation (S20) and it relies on computing the DFT of the autocovariance function. Note that computation of DFT is highly efficient thanks to the *Fast Fourier Transform* (FFT) method developed in [4].

Typically an output trajectory is obtained as a discrete time-series $x_1, \dots, x_N$ which represents the abundance of the output species $\mathbf{X_n}$ at uniformly-spaced time-points $t_1, \dots, t_N$ in some time-interval $[T_0, T_f]$. In other words for time-step $\delta = (T_f - T_0)/(N-1)$ we have

$$t_i = T_0 + (i-1)\delta \quad \text{and} \quad x_i = X_n(t_i) \quad \text{for} \quad i = 0, 1 \dots, N.$$

As we are only interested in the stationary behaviour, typically one assumes that stationarity is reached at some *cut-off* time $T_c \in [T_0, T_f]$, and then the part of the time-series which is associated with the initial time-interval $[T_0, T_c]$ is discarded. Letting

$$N_c = 1 + \left\lceil \frac{T_c - T_0}{\delta} \right\rceil$$

we shall estimate the PSD using the truncated time-series $x_{N_c}, \dots, x_N$. Here $\lceil x \rceil$ denotes the smallest integer greater than $x$ and we shall assume that the size $L := (N - N_c + 1)$ of this truncated series is even.

From the truncated time-series, we compute the empirical mean as

$$\widehat{\mu} = \frac{1}{L} \sum_{j=N_c}^{N_c+L-1} x_j$$

and then obtain the mean-zero time-series $\widetilde{x}_{N_c}, \dots, \widetilde{x}_N$ by letting

$$\widetilde{x}_j = x_j - \widehat{\mu} \quad \text{for each} \quad j = N_c, N_c + 1, \dots, (N_c + L - 1).$$

In the periodogram approach we compute the DFT of this time series $\widetilde{x}_{N_c}, \dots, \widetilde{x}_{N_c+L-1}$ to estimate the power spectral density $\widehat{S}_{X_n}(\omega)$ as

$$\widehat{s}_{X_n}(\omega_k) = \frac{\delta}{L} \left| \sum_{j=N_c}^{N_c+L-1} \widetilde{x}_j e^{-\omega_k (j-N_c)\delta} \right|^2 \quad \text{for} \quad \omega_k = \frac{2\pi k}{\delta L} \quad \text{and} \quad k = 0, \dots, \frac{L}{2}.$$

In the Wiener-Khintchine approach we first compute the autocovariance $\mathrm{ACV}_{X_n}(\tau)$ for any time $\tau$ of the form $\tau = k\delta$ as

$$\widehat{\mathrm{ACV}}_{X_n}(\tau) = \widehat{a}_k := \frac{1}{L} \sum_{i=N_c}^{N-k} \widetilde{x}_i \widetilde{x}_{i+k}.$$

Note that the denominator is $L$ instead of $(L - k)$. We fix a $K \ll (N - N_c)$ and compute the DFT of the ACV time series $\widehat{a}_0, \dots, \widehat{a}_K$ to estimate the power spectral density $\widehat{S}_{X_n}(\omega)$ as

$$\widehat{s}_{X_n}(\omega_k) = 2\delta \sum_{j=1}^{L} \widehat{a}_j e^{-\omega_k (j-1)\delta} \quad \text{for} \quad \omega_k = \frac{2\pi k}{\delta L} \quad \text{and} \quad k = 0, \dots, \frac{L}{2}.$$

The main problem with the Wiener-Khintchine approach is that it is computationally very cumbersome because it relies on computation of the autocovariance function that requires several iterations over the signal time-series. Hence the periodogram approach is the more popular method for estimating a signal's PSD. However the main drawback of this approach is that it is *inconsistent*, i.e. even for discrete time-series the variance of the PSD estimator does not converge to zero as the length of the time-series goes to infinity (see Chapter 7 in [8]). Due to this inconsistency the PSD estimates based on the periodogram of a single output time-series can be extremely noisy and unreliable. If one has several independent output time-series then one can mitigate this problem by averaging the PSDs estimated for all the time-series (see Chapter 7 in [8]). This averaging makes the PSD estimate less noisy and much more reliable.

### S2 Model based estimation of PSD

Suppose we have a reaction network model that yields an output trajectory $(X_n(t))_{t\geq 0}$. If we can efficiently estimate the PSD for such a trajectory then it provides a powerful new approach for connecting the model with the single-cell frequency spectrum, which can pave the way for many applications in systems and synthetic biology. The *first-principles* approach for estimating the PSD from a stochastic model would be to first simulate a trajectory, sample it at finitely many time-points $t_1, \dots, t_N$ and then use the *periodogram* described in Section S1.3 to obtain an estimate of the PSD. This approach often produces a noisy and unreliable estimate because irrespective of how large the number of time-points ($N$) is, certain frequencies will always be poorly represented in the signal leading to large statistical inaccuracies in the PSD estimate at these frequencies. However as discussed in Section S1.3, we can improve our estimate by simulating several independent trajectories and averaging their PSDs. This averaging procedure would increase the computational costs as trajectory simulations are time-consuming. More importantly, the averaged PSD may still not be accurate because it is based on discrete sampling of continuous signals. It is known that this sampling can cause the problem of *aliasing* which distorts the estimated PSD by introducing frequency components corresponding to the sampling operation (see Chapter 1 in [8]). As shown by the Nyquist's Sampling Theorem [21] we can remove this aliasing effect by choosing the time-step parameter $\delta$ to be smaller than half of the reciprocal of the maximum frequency represented in the signal. However for stochastic dynamics it is difficult to use this criterion as the range of frequencies in the signal is often very wide and picking a very small $\delta$ can lead to an exorbitant computational burden.

These issues motivated us to devise a parametric approach to estimate the PSD that rather than relying on only the output signal, uses full information contained in the stochastic model of the dynamics. Our method does not require any discrete-sampling and it efficiently produces a reliable representation of the PSD based on a single trajectory simulation of the stochastic model. To develop this method we utilise a connection between the PSD and the resolvent operator for a continuous-time Markov chain. Apart from helping us devise our parametric PSD estimation method for general reaction networks, this connection also yields a PSD decomposition result for *linear* reaction networks where all the propensity functions are affine (see Section S2.3). The rest of this section is devoted to the theoretical and computational description of our method.

#### S2.1 The resolvent operator

The resolvent operator corresponding to the Markov chain generator $\mathbb{A}$ plays a central role in the development of our method for PSD estimation. We now briefly describe this operator and discuss some of its properties. Let $(X(t))_{t\geq 0}$ be a CTMC with state-space $\mathcal{E}$ and generator $\mathbb{A}$. On the space of bounded real-valued functions on $\mathcal{E}$, the transition semigroup $\mathbb{T}(t)$ generated by $\mathbb{A}$ is the linear operator which for any output function $f$ evaluates the expected value of $f(X(t))$ given that the initial state is $x$, i.e.

$$\mathbb{T}(t)f(x) = \mathbb{E}\left(f(X(t))|X(0) = x\right).$$

Informally we can view the transition semi-group as the exponential operator

$$\mathbb{T}(t)f(x) = e^{t\mathbb{A}}f(x).$$

For any complex number $z \in \mathbb{C}$ with a positive real part (i.e. Real$(z) > 0$) the resolvent operator (see [15]) is defined as the infinite integral

$$\mathbb{R}(z)f(x) = \int_0^\infty e^{-zt}\mathbb{T}(t)f(x)dt. \tag{S25}$$

On the Banach space $\mathcal{B}_b(\mathcal{E})$ the resolvent operator is bounded because the transition semi-group $\mathbb{T}(t)$ is a contraction operator. We can view the resolvent operator as

$$\mathbb{R}(z)f(x) = (z\mathbf{I} - \mathbb{A})^{-1} f(x) = \sum_{k=0}^{\infty} z^{-(k+1)}\mathbb{A}^k f(x), \tag{S26}$$

where $\mathbf{I}$ is the identity operator.

Let $\sigma_0, \sigma_1, \sigma_2, \dots$ be the set of eigenvalues of $\mathbb{A}$, repeated according to their multiplicity and arranged in descending order of their real parts

$$\text{Real}(\sigma_k) \geq \text{Real}(\sigma_j) \quad \text{for} \quad k < j. \tag{S27}$$

At the CTMC is ergodic, we must have $\sigma_0 = 0$ and Real$(\sigma_j) < 0$ for all $j \geq 1$. Let $\phi_j$ be the eigenfunction corresponding to eigenvalue $\sigma_j$. Note that $\phi_0$ is simply the constant function $\mathbf{1}$ which takes value one at each $x$.

**Lemma S2.1** *The set of eigenvalues of the resolvent operator is exactly the set*

$$\sigma(\mathbb{R}(z)) = \{(z - \sigma_j)^{-1} : j = 0, 1, \dots\}.$$

*Moreover the eigenfunction corresponding to eigenvalue* $(z - \sigma_j)^{-1}$ *is* $\phi_j$*.*

**Proof.** Let us start with an eigenvalue-eigenfunction pair $(\sigma_j, \phi_j)$ for $\mathbb{A}$. Then since

$$\mathbb{A}\phi_j(x) = \sigma_j\phi_j(x) \tag{S28}$$

we have

$$\mathbb{T}(t)\phi_j(x) = e^{\sigma_j t}\phi_j(x)$$

and hence from (S25) we obtain

$$\mathbb{R}(z)\phi_j(x) = \int_0^\infty e^{-zt}e^{\sigma_j t}\phi_j(x)dt = \int_0^\infty e^{-(z-\sigma_j)t}\phi_j(x)dt = \frac{\phi_j(x)}{z - \sigma_j}.$$

Therefore $\phi_j$ is also an eigenfunction for $\mathbb{R}(z)\phi_j(x)$ with eigenvalue $(z - \sigma_j)^{-1}$. To prove the converse, note that the resolvent operator satisfies the following identity

$$\mathbb{A}\mathbb{R}(z) = \mathbb{R}(z)\mathbb{A} = z\mathbb{R}(z) - \mathbf{I}. \tag{S29}$$

Suppose $(\sigma, \phi)$ is an eigenvalue-eigenfunction pair for $\mathbb{R}(z)$. Then applying this identity we obtain

$$\sigma \mathbb{A}\phi = \mathbb{A}\mathbb{R}(z)\phi = z\mathbb{R}(z)\phi - \phi = (z\sigma - 1)\phi.$$

Hence $\phi$ is an eigenfunction for generator $\mathbb{A}$ corresponding to eigenvalue $(z - \sigma^{-1})$. Therefore $\phi = \phi_j$ and $\sigma = (z - \sigma_j)^{-1}$ for some $j$. This completes the proof of this lemma. □

Consider the Hilbert space $\mathcal{L}_2(\mathcal{E})$ of those real-valued functions on $\mathcal{E}$ that have zero expectation and finite variance under the stationary distribution $\pi$, i.e.

$$\mathcal{L}_2(\mathcal{E}) = \{f : \mathcal{E} \to \mathbb{R} : \mathbb{E}_\pi(f) = 0 \text{ and } \mathbb{E}_\pi(f^2) < \infty\}. \tag{S30}$$

On this Hilbert space the inner product is given by $\langle f, g\rangle_{\mathcal{L}_2(\mathcal{E})} = \mathbb{E}_\pi(fg)$.

For each $j \geq 1$ taking expectation on both sides of (S28) w.r.t. $\pi$, using (S12) and $\sigma_j \neq 0$ we obtain $\mathbb{E}_\pi(\phi_j) = 0$. Hence for each $j \geq 1$, if $\phi_j$ has finite stationary variance (i.e. $\mathbb{E}_\pi(\phi_j^2) < \infty$) then it belongs to $\mathcal{L}_2(\mathcal{E})$. We shall develop our method under the assumption that $\mathcal{L}_2(\mathcal{E})$ is the linear span of $\{\phi_1, \phi_2, \dots\}$

$$\mathcal{L}_2(\mathcal{E}) = \mathrm{Span}(\{\phi_1, \phi_2, \dots\}). \tag{S31}$$

Under this assumption, the resolvent operator $\mathbb{R}(z)$ maps the Hilbert space $\mathcal{L}_2(\mathcal{E})$ to itself and it can also be seen as the limit of finite rank operators (see [14]). Therefore by Theorem 4.4 in Chapter 2 of [3], the resolvent $\mathbb{R}(z)$ is a compact operator. Hence its set of eigenvalues can only have zero as an accumulation point (see [25]), thereby showing that

$$(z - \sigma_j)^{-1} \to 0 \text{ as } j \to \infty. \tag{S32}$$

Now consider an output function $f \in \mathcal{L}_2(\mathcal{E})$. We can express it as a linear combination of eigenfunctions $\{\phi_1, \phi_2, \dots\}$ as

$$f(x) = \sum_{j=1}^{\infty} c_j \phi_j(x) \tag{S33}$$

and the action of the resolvent operator $\mathbb{R}(z)$ on this function is given by

$$\mathbb{R}(z)f(x) = \sum_{j=1}^{\infty} \frac{c_j}{z - \sigma_j}\phi_j(x). \tag{S34}$$

As $\mathrm{Real}(\sigma_1) < 0$ and (S27) holds, the resolvent operator remains well-defined for $z$ on the imaginary axis.

#### S2.2 Connection between the resolvent operator and PSD

We now establish a relation between the power spectral density $S_{X_n}(\omega)$ and the resolvent operator. Let $f$ be the function

$$f(x) = x_n - \mathbb{E}_\pi(X_n). \tag{S35}$$

Note that $f \in \mathcal{L}_2(\mathcal{E})$ and recall from (S20) that $S_{X_n}(\omega)$ is the Fourier transform of the autocovariance function $\text{ACV}_{X_n}(\tau)$ defined by (S18). Without loss of generality we let $(X(t))_{t\geq 0}$ be the CTMC with generator $\mathbb{A}$ and initial state $X(0)$ distributed according to the stationary distribution $\pi$. Then using $\text{ACV}_{X_n}(\tau) = \text{ACV}_{X_n}(-\tau)$ (i.e. autocovariance is an even function) we can rewrite (S20) as

$$\begin{aligned}
S_{X_n}(\omega) &= \int_0^\infty \text{ACV}_{X_n}(\tau)\left(e^{-i\omega\tau} + e^{i\omega\tau}\right) d\tau \\
&= \int_0^\infty \mathbb{E}(f(X(0))f(X(\tau)))\left(e^{-i\omega\tau} + e^{i\omega\tau}\right) d\tau \\
&= \mathbb{E}\left[f(X(0))\int_0^\infty \mathbb{E}\left(f(X(\tau))|X(0)\right)\left(e^{-i\omega\tau} + e^{i\omega\tau}\right) d\tau\right] \\
&= \mathbb{E}\left[f(X(0))\int_0^\infty e^{-i\omega\tau}\mathbb{T}_\tau f(X(0))d\tau\right] + \mathbb{E}\left[f(X(0))\int_0^\infty e^{i\omega\tau}\mathbb{T}_\tau f(X(0))d\tau\right] \\
&= \mathbb{E}\left[f(X(0))\mathbb{R}(i\omega)f(X(0))\right] + \mathbb{E}\left[f(X(0))\mathbb{R}(-i\omega)f(X(0))\right] \\
&= 2\text{Real}\left(\mathbb{E}\left[f(X(0))\mathbb{R}(i\omega)f(X(0))\right]\right),
\end{aligned}$$

where the last relation holds because $\mathbb{R}(i\omega)$ and $\mathbb{R}(-i\omega)$ are complex conjugates. This calculation shows that if we define $G(z)$ as

$$G(z) = \mathbb{E}_\pi\left(f\mathbb{R}(z)f\right), \tag{S36}$$

then the PSD $S_{X_n}(\omega)$ is given by

$$S_{X_n}(\omega) = 2\text{Real}(G(i\omega)). \tag{S37}$$

In our method we first estimate $G(z)$ and then obtain the PSD using (S37). In Section S2.3 we show that $G(z)$ can be exactly computed for linear reaction networks and we also consider the situation when such a network is stimulated by an upstream signal. In Section S2.4 we present our method to efficiently estimate $G(z)$ for a general nonlinear network with Monte Carlo simulations

#### S2.3 Exact PSD derivation for linear networks

In this section we prove formula (12) which provides an analytical expression for the exact PSD for linear networks. We prove this by exploiting the connection between the PSD and the resolvent operator described in Section S2.2. This enables us to obtain interesting insights about certain simple motifs commonly found in biological networks, and it also provides us with a large collection of test examples where we can check the accuracy of the approximation provided by our PSD estimation method that we shall develop in Section S2.4.

Linear networks are characterised by all propensity functions being affine functions of the state variables. Under the mass-action hypothesis (S2), linear networks are necessarily unimolecular in the sense that they only consist of reactions with at most one reactant (i.e. $\sum_i \nu_{ik} \leq 1$ for each reaction of the form (S1)). In other words, all the reactions are of the form $\emptyset \longrightarrow \star$ and $\mathbf{X_i} \longrightarrow \star$, where $\star$ represents any linear combination of species. Suppose there are $d$ species (viz. $\mathbf{X_1}, \dots, \mathbf{X_d}$) and $K$ reactions. By definition, for linear networks, the vector of propensity functions $\lambda(x) = (\lambda_1(x), \dots, \lambda_K(x))$ is described by an affine map

$$\lambda(x) = \Lambda x + \widetilde{b} \tag{S38}$$

where $\Lambda$ is some $K \times d$ matrix and $\widetilde{b}$ is a $K \times 1$ vector. If we consider a real-valued function $g(x) = \alpha^T x$ for some vector $\alpha = (\alpha_1, \dots, \alpha_d)$, then the image of this function under generator $\mathbb{A}$ can be expressed as

$$\mathbb{A} g(x) = \alpha^T S \Lambda x + \alpha^T S \widetilde{b},$$

where $S$ is the $d \times K$ stoichiometry matrix with columns $\zeta_1, \dots, \zeta_K$ (see Section S1.1). Henceforth let $A$ be the $d \times d$ matrix and $b$ be the $d \times 1$ vector defined by

$$A = S\Lambda \qquad \text{and} \qquad b = S\widetilde{b}.$$

Under our assumption of exponential ergodicity, the matrix $A$ is Hurwitz-stable, i.e. all its eigenvalues have strictly negative real parts. One can easily show that the expectation of the species copy-number under the stationary distribution $\pi$ is given by

$$\overline{x} = \mathbb{E}_\pi(X) = -A^{-1} b \tag{S39}$$

and the stationary variance-covariance matrix $\Sigma$ can be computed by solving the following Lyapunov equation

$$A\Sigma + \Sigma A^T + S \text{diag}(\Lambda \overline{x} + \widetilde{b}) S^T = 0. \tag{S40}$$

**Proof.**[Proof of formula (12) in the main paper] Note that the action of generator $\mathbb{A}$ on a linear function $g(x) = \alpha^T x$ can be expressed as

$$\mathbb{A} g(x) = \alpha^T A x + \alpha^T b.$$

Since the output function $f(x) = x_n - \mathbb{E}_\pi(X_n)$ is affine, we can assume that its image under the resolvent $\mathbb{R}(z)$ is also affine i.e.

$$\mathbb{R}(z) f(x) = g_z(x) := \alpha_z^T x + \beta_z.$$

This implies that

$$\begin{aligned} f(x) &= (z\mathbf{I} - \mathbb{A}) g_z(x) \\ &= (z\mathbf{I} - \mathbb{A}) \left( \alpha_z^T x + \beta_z \right) \\ &= z\alpha_z^T x + z\beta_z - \alpha_z^T A x - \alpha_z^T b \\ &= \alpha_z^T (z\mathbf{I} - A) x + z\beta_z - \alpha_z^T b. \end{aligned}$$

As this must hold for each $x$, we must have

$$\alpha_z^T (z\mathbf{I} - A) = e_n^T \qquad \text{and} \qquad z\beta_z = \alpha_z^T b - \mathbb{E}_\pi(X_n).$$

Since matrix $A$ is Hurwitz stable, if $z$ is a complex number with a non-negative real part then the matrix $(z\mathbf{I} - A)$ is invertible and hence $\alpha_z$ can be computed as

$$\alpha_z = \left[(z\mathbf{I} - A)^{-1}\right]^T e_n. \tag{S41}$$

Now from (S36) we obtain

$$G(z) = \mathbb{E}_\pi \left(f\mathbb{R}(z)f\right) = \mathbb{E}_\pi \left[(X_n - \mathbb{E}_\pi(X_n)) \left(\alpha_z^T X + \beta_z\right)\right] = e_n^T \Sigma \alpha_z.$$

Therefore using (S41) and (S37) we get that the power spectral density is given by

$$S_{X_n}(\omega) = 2\text{Real}(e_n^T \Sigma \left[(i\omega\mathbf{I} - A)^{-1}\right]^T e_n) = 2\text{Real}(e_n^T (i\omega\mathbf{I} - A)^{-1}\Sigma e_n).$$

Expressing the matrix $2\text{Real}((i\omega\mathbf{I} - A)^{-1})$ as

$$\begin{aligned}
2\text{Real}((i\omega\mathbf{I} - A)^{-1}) &= (i\omega\mathbf{I} - A)^{-1} + (-i\omega\mathbf{I} - A)^{-1} \\
&= (i\omega\mathbf{I} - A)^{-1}(-i\omega\mathbf{I} - A)^{-1}\left[(-i\omega\mathbf{I} - A) + (i\omega\mathbf{I} - A)\right] \\
&= -2(i\omega\mathbf{I} - A)^{-1}(-i\omega\mathbf{I} - A)^{-1}A \\
&= -2\left((-i\omega\mathbf{I} - A)(i\omega\mathbf{I} - A)\right)^{-1}A \\
&= -2\left(\omega^2\mathbf{I} + A^2\right)^{-1}A
\end{aligned} \tag{S42}$$

completes the proof of this theorem. □

**Remark S2.2** *It is evident from this proof that if we want to compute the PSD of some linear combination of species* $\mathbf{Y} = c_1\mathbf{X_1} + \cdots + c_d\mathbf{X_d}$ *then we can simply replace* $e_n$ *by the coefficient vector* $c = (c_1, \dots, c_d)$ *to obtain*

$$S_Y(\omega) = -2c^T(\omega^2\mathbf{I} + A^2)^{-1}A\Sigma c.$$

We now consider the situation where a linear network is not operating in isolation but it is being stimulated by $m$ independent time-varying signals $(Y_1(t))_{t\geq 0}, \dots, (Y_m(t))_{t\geq 0}$. Our goal is to understand how the PSD $S_{X_n}(\omega)$ of our output species $\mathbf{X_n}$ depends on the PSDs $S_{Y_j}(\omega)$ for $j = 1, \dots, m$. We assume that stimulation occurs through $m$ zeroth-order reactions of the form

$$\emptyset \xrightarrow{\theta_k Y_k(t)} \sum_{j=1}^{d} c_{jk}\mathbf{X_j} \tag{S43}$$

for $k = 1, \dots, m$. Here $\theta_k$ is a positive constant and $c_k = (c_{1k}, \dots, c_{dk}) \in \mathbb{Z}_+^d$ is the stoichiometric vector representing the number of molecules of each species $\mathbf{X_1}, \dots, \mathbf{X_d}$ created by reaction $k$. Recall that these network species participate in $K$ reactions of their own with propensity functions $\lambda_1(x), \dots, \lambda_K(x)$ and stoichiometric vectors $\zeta_1, \dots, \zeta_K$. Due to stimulation, the infinitesimal generator for the process $(X(t))_{t\geq 0}$ depends on the current signal state $Y(t) = (Y_1(t), \dots, Y_m(t)) = y = (y_1, \dots, y_m)$ and it can be expressed as

$$\mathbb{A}_y g(x) = \sum_{k=1}^{K} \lambda_k(x)\left(g(x + \zeta_k) - g(x)\right) + \sum_{j=1}^{m} \theta_j y_j \left(g(x + c_j) - g(x)\right) \tag{S44}$$

Let matrix $A$ and vector $b$ be as before, let $B$ be the $d \times m$ matrix with columns $c_k$ for $k = 1, \dots, m$ and let

$$\Theta = \mathrm{diag}(\theta_1, \dots, \theta_m).$$

We shall assume that process $(Y(t))_{t \geq 0}$, which includes all the stimulating signals, is an exponentially ergodic Markov process. Let $\overline{y} = (\overline{y}_1, \dots, \overline{y}_m)$ be its stationary expectation. Then one can show (see proof of Theorem S2.3) that the stationary expectation of the species copy-number is given by

$$\overline{x} = \mathbb{E}_\pi(X) = -A^{-1}(b + B\Theta\overline{y}). \tag{S45}$$

Let $\overline{\Sigma}$ be the $d \times d$ positive semidefinite matrix computed by solving the following Lyapunov equation

$$A\overline{\Sigma} + \overline{\Sigma}A^T + S\mathrm{diag}(\Lambda\overline{x} + \widetilde{b})S^T + B\Theta B^T = 0, \tag{S46}$$

where $\Lambda$ and $\widetilde{b}$ are as in (S38). Observe that $\overline{\Sigma}$ can be viewed as the stationary variance-covariance matrix for the process $(X(t))_{t \geq 0}$ *when each stimulating signal is deterministic and fixed to its stationary mean at all times*, i.e. $Y(t) = \overline{y}$ for all $t \geq 0$. We now present the proof of the PSD decomposition result (Theorem 3.1) in the main paper which is stated as Theorem S2.3 below. This result provides an analytic relationship between the PSD $S_{X_n}(\omega)$ of our output species $\mathbf{X_n}$ and the of PSDs $S_{Y_j}(\omega)$ for $j = 1, \dots, m$.

**Theorem S2.3** *Consider a linear reaction network comprising species* $\mathbf{X_1}, \dots, \mathbf{X_d}$*, stimulated by independent time-varying signals* $(Y_1(t))_{t \geq 0}, \dots, (Y_m(t))_{t \geq 0}$*, through zeroth-order reactions of the form* (S43). *We assume that each* $Y_j$ *is an exponentially ergodic Markov process with PSD* $S_{Y_j}(\omega)$*. Defining* $d \times d$ *matrices* $A$ *and* $\overline{\Sigma}$ *as above, the PSD of the output species* $\mathbf{X_n}$ *is given by*

$$S_{X_n}(\omega) = -2e_n^T(\omega^2\mathbf{I} + A^2)^{-1}A\overline{\Sigma}e_n + \sum_{j=1}^{m} \theta_j^2 |e_n^T(A + i\omega\mathbf{I})^{-1}c_j|^2 S_{Y_j}(\omega).$$

**Proof.** Let $\mathbb{B}$ be the generator for the Markov process $(Y(t))_{t \geq 0}$. We denote the generator for the *combined* dynamics $(X(t), Y(t))_{t \geq 0}$ as $\mathbb{C}$ and it can be defined by its action on product functions of the form $g(x)h(y)$ as

$$\mathbb{C}(g(x)h(y)) = h(y)\mathbb{A}_y g(x) + g(x)\mathbb{B}h(y),$$

where $\mathbb{A}_y$ is given by (S44). Under our assumptions, the combined process is ergodic and we denote its stationary distribution by $\pi$. Henceforth let $\mathbb{R}_\mathbb{C}(z)$ (resp. $\mathbb{T}_\mathbb{C}(t)$) and $\mathbb{R}_\mathbb{B}(z)$ (resp. $\mathbb{T}_\mathbb{B}(t)$) denote the resolvent operators (resp. transition semigroups) corresponding to $\mathbb{C}$ and $\mathbb{B}$ respectively. Note that the definition of each operator can be extended from real-valued functions to vector or matrix-valued functions by applying it to each component.

For the vector-valued function $\phi(x) = x$ the image under generator $\mathbb{C}$ can be expressed as

$$\mathbb{C}\phi(x) = \mathbb{A}_y\phi(x) = Ax + b + B\Theta y.$$

Setting $\mathbb{E}_\pi(\mathbb{C}\phi(x)) = 0$ (see (S12)) we see that the stationary mean $\overline{x}$ is related to the stationary signal mean $\overline{y}$ according to (S45).

Now let $g(x) = x - \overline{x}$ and $h(y) = y - \overline{y}$. Due to Dynkin's Theorem (see [9]) we have

$$\begin{aligned}\frac{d}{dt}\mathbb{T}_{\mathbb{C}}(t)g(x) &= \mathbb{T}_{\mathbb{C}}(t)\mathbb{C}g(x)\\ &= \mathbb{T}_{\mathbb{C}}(t)(Ax + b + B\Theta y)\\ &= \mathbb{T}_{\mathbb{C}}(t)Ag(x) + \mathbb{T}_{\mathbb{C}}(t)B\Theta h(y),\end{aligned}$$

where the last relation holds due to (S45). Applying integration by parts we obtain

$$\begin{aligned}\mathbb{R}_{\mathbb{C}}(z)g(x) &= \int_0^\infty e^{-zt}\mathbb{T}_{\mathbb{C}}g(x)dt\\ &= z^{-1}g(x) + z^{-1}\int_0^\infty e^{-zt}\mathbb{T}_{\mathbb{C}}\mathbb{C}g(x)dt\\ &= z^{-1}g(x) + z^{-1}\int_0^\infty e^{-zt}\mathbb{T}_{\mathbb{C}}Ag(x)dt + z^{-1}\int_0^\infty e^{-zt}\mathbb{T}_{\mathbb{C}}B\Theta h(y)dt\\ &= z^{-1}g(x) + z^{-1}A\int_0^\infty e^{-zt}\mathbb{T}_{\mathbb{C}}g(x)dt + z^{-1}B\Theta\int_0^\infty e^{-zt}\mathbb{T}_{\mathbb{C}}h(y)dt\\ &= z^{-1}g(x) + z^{-1}A\mathbb{R}_{\mathbb{C}}(z)g(x) + z^{-1}B\Theta\mathbb{R}_{\mathbb{C}}(z)h(y),\end{aligned}$$

where we have exploited the linearity of the transition semigroup operator in the second last step. Multiplying by $z$ and solving for $\mathbb{R}_{\mathbb{C}}(z)g(x)$ we get

$$\mathbb{R}_{\mathbb{C}}(z)g(x) = (z\mathbf{I} - A)^{-1}\left(g(x) + B\Theta\mathbb{R}_{\mathbb{C}}(z)h(y)\right).$$

Multiplying both sides by the row vector $g^T(x) = (x - \overline{x})^T$, and taking expectation w.r.t. $\pi$ we obtain

$$\begin{aligned}G_X(z) :&= \mathbb{E}_\pi\left([\mathbb{R}_{\mathbb{C}}(z)g(x)]g^T(x)\right) = (z\mathbf{I} - A)^{-1}\left(\mathbb{E}_\pi(g(x)g^T(x)) + B\Theta\mathbb{E}_\pi([\mathbb{R}_{\mathbb{C}}(z)h(y)]g^T(x))\right)\\ &= (z\mathbf{I} - A)^{-1}\left(\Sigma_X + B\Theta\mathbb{E}_\pi([\mathbb{R}_{\mathbb{C}}(z)h(y)]g^T(x))\right),\end{aligned} \tag{S47}$$

where $\Sigma_X$ is the stationary variance-covariance matrix for the molecular counts of species $\mathbf{X_1}, \dots, \mathbf{X_d}$.

We now let $H_k(y) = \mathbb{C}^k h(y) = \mathbb{B}^k h(y)$ for any $k \in \mathbb{Z}_+$. Then

$$\begin{aligned}\mathbb{C}(H_k(y)g^T(x)) &= H_k(y)\mathbb{A}_y g^T(x) + [\mathbb{B}H_k(y)]g^T(x)\\ &= H_k(y)(Ax + b + B\Theta y)^T + H_{k+1}(y)g^T(x)\\ &= H_k(y)(Ag(x) + B\Theta h(y))^T + H_{k+1}(y)g^T(x)\\ &= H_k(y)g^T(x)A^T + H_k(y)h^T(y)(B\Theta)^T + H_{k+1}(y)g^T(x).\end{aligned}$$

Using (S12) we obtain

$$\mathbb{E}_\pi(H_k(y)h^T(y))(B\Theta)^T = -\mathbb{E}_\pi(H_k(y)g^T(x))A^T - \mathbb{E}_\pi(H_{k+1}(y)g^T(x)). \tag{S48}$$

Recall the resolvent expansion formula (S26). Dividing both sides of (S48) by $z^{k+1}$ and summing over $k \in \mathbb{Z}_+$ we get

$$\mathbb{E}_\pi([\mathbb{R}_\mathbb{C}(z)h(y)]h^T(y))(B\Theta)^T = -\mathbb{E}_\pi([\mathbb{R}_\mathbb{C}(z)h(y)]g^T(x))A^T - \mathbb{E}_\pi([\mathbb{R}_\mathbb{C}(z)\mathbb{C}h(y)]g^T(x)). \tag{S49}$$

Due to the resolvent identity (S29), $\mathbb{R}_\mathbb{C}(z)\mathbb{C} = z\mathbb{R}_\mathbb{C} - \mathbf{I}$. Plugging this in (S49) we obtain

$$\mathbb{E}_\pi([\mathbb{R}_\mathbb{C}(z)h(y)]h^T(y))(B\Theta)^T = -\mathbb{E}_\pi([\mathbb{R}_\mathbb{C}(z)h(y)]g^T(x))(z\mathbf{I} + A^T) + \mathbb{E}_\pi(h(y)g^T(x)),$$

which multiplication by $(z\mathbf{I} + A^T)^{-1}$ yields

$$\mathbb{E}_\pi([\mathbb{R}_\mathbb{C}(z)h(y)]g^T(x)) = \left(\mathrm{Cov}_\pi(X,Y) - \mathbb{E}_\pi([\mathbb{R}_\mathbb{C}(z)h(y)]h^T(y))(B\Theta)^T\right)(z\mathbf{I} + A^T)^{-1}, \tag{S50}$$

where we have used the fact that $\mathbb{E}_\pi(h(y)g^T(x)) = \mathrm{Cov}_\pi(X,Y)$ is the $m \times d$ stationary covariance matrix between the molecular counts of species $\mathbf{X_1}, \dots, \mathbf{X_d}$ and the signal $Y$. Our next task is to relate this covariance matrix with the variance-covariance matrix $\Sigma_X$ encountered in equation (S47). For this purpose we apply the generator $\mathbb{C}$ on the matrix $g(x)g^T(x)$ to obtain

$$\begin{aligned}\mathbb{C}(g(x)g^T(x)) &= (Ag(x) + B\Theta h(y))g^T(x) + g(x)(Ag(x) + B\Theta h(y))^T \\ &\quad + S\mathrm{diag}(\Lambda\overline{x} + \widetilde{b})S^T + B\Theta B^T \\ &= Ag(x)g^T(x) + B\Theta h(y)g^T(x) + g(x)g^T(x)A^T + g(x)h^T(y)(B\Theta)^T \\ &\quad + S\mathrm{diag}(\Lambda\overline{x} + \widetilde{b})S^T + B\Theta B^T,\end{aligned}$$

where $\Lambda$ and $\widetilde{b}$ are as in (S38). Using (S12), $\Sigma_X = \mathbb{E}_\pi(g(x)g^T(x))$ and $\mathrm{Cov}_\pi(X,Y) = \mathbb{E}_\pi(h(y)g^T(x))$ we get

$$A\Sigma_X + B\Theta\mathrm{Cov}_\pi(X,Y) + \Sigma_X A^T + \mathrm{Cov}_\pi^T(X,Y)(B\Theta)^T + S\mathrm{diag}(\Lambda\overline{x} + \widetilde{b})S^T + B\Theta B^T = 0.$$

We decompose $\Sigma_X$ as

$$\Sigma_X = \overline{\Sigma} + \widetilde{\Sigma}, \tag{S51}$$

where $\overline{\Sigma}$ is the solution of the Lyapunov equation (S46). Subtracting this Lyapunov equation from the one above, we obtain the following equation for $\widetilde{\Sigma}$

$$A\widetilde{\Sigma} + \widetilde{\Sigma}A^T + B\Theta\mathrm{Cov}_\pi(X,Y) + \mathrm{Cov}_\pi^T(X,Y)(B\Theta)^T = 0. \tag{S52}$$

Substituting $\Sigma_X$ from (S51) and $\mathbb{E}_\pi([\mathbb{R}_\mathbb{C}(z)h(y)]g^T(x))$ from (S50) in (S47) we get

$$G_X(z) = (z\mathbf{I} - A)^{-1}\overline{\Sigma} - (z\mathbf{I} - A)^{-1}B\Theta\mathbb{E}_\pi([\mathbb{R}_\mathbb{C}(z)h(y)]h^T(y))(B\Theta)^T(z\mathbf{I} + A^T)^{-1} + \Psi(z) \tag{S53}$$

where

$$\Psi(z) := (z\mathbf{I} - A)^{-1}\widetilde{\Sigma} + (z\mathbf{I} - A)^{-1}B\Theta\mathrm{Cov}_\pi(X,Y)(z\mathbf{I} + A^T)^{-1}.$$

Upon multiplying (S52) on the left by $(z\mathbf{I}-A)^{-1}$, using $(z\mathbf{I}-A)^{-1}A = -\mathbf{I} + z(z\mathbf{I}-A)^{-1}$ and rearranging we obtain

$$(z\mathbf{I}-A)^{-1}\widetilde{\Sigma}(z\mathbf{I}+A^T) + (z\mathbf{I}-A)^{-1}B\Theta\mathrm{Cov}_\pi(X,Y) = \widetilde{\Sigma} - (z\mathbf{I}-A)^{-1}\mathrm{Cov}_\pi^T(X,Y)(B\Theta)^T.$$

Now multiplying on the right by $(z\mathbf{I}+A^T)^{-1}$ yields

$$\Psi(z) = \widetilde{\Sigma}(z\mathbf{I}+A^T)^{-1} - (z\mathbf{I}-A)^{-1}\mathrm{Cov}_\pi^T(X,Y)(B\Theta)^T(z\mathbf{I}+A^T)^{-1}.$$

Therefore

$$\Psi^T(-z) = -(z\mathbf{I}-A)^{-1}\widetilde{\Sigma} - (z\mathbf{I}-A)^{-1}B\Theta\mathrm{Cov}_\pi(X,Y)(z\mathbf{I}+A^T)^{-1} = -\Psi(z).$$

Hence for $z = i\omega$ we have

$$\Psi(i\omega) = -\Psi^T(-i\omega) = -\Psi^\dagger(i\omega),$$

where super-script $\dagger$ denotes the conjugate transpose. This shows that $\Psi(i\omega)$ is a skew-Hermitian matrix which implies that real part of matrix $(\Psi(i\omega)+\Psi^T(i\omega))$ is component-wise zero. Setting $z = i\omega$ in (S53) and adding $G_X(i\omega)$ to its transpose we arrive at

$$\begin{aligned} G_X(i\omega) + G_X^T(i\omega) &= (i\omega\mathbf{I}-A)^{-1}\overline{\Sigma} + \overline{\Sigma}(i\omega\mathbf{I}-A^T)^{-1} \\ &\quad - (i\omega\mathbf{I}-A)^{-1}B\Theta\mathbb{E}_\pi([\mathbb{R}_\mathbb{C}(i\omega)h(y)]h^T(y))(B\Theta)^T(i\omega\mathbf{I}+A^T)^{-1} \\ &\quad - (i\omega\mathbf{I}+A)^{-1}B\Theta\mathbb{E}_\pi(h(y)[\mathbb{R}_\mathbb{C}(i\omega)h(y)]^T)(B\Theta)^T(i\omega\mathbf{I}-A^T)^{-1} \\ &\quad + \Psi(i\omega) + \Psi^T(i\omega). \end{aligned}$$

Multiplying by $e_n^T$ on the left and $e_n$ on the right and taking real part we obtain

$$\begin{aligned} 2\mathrm{Real}(e_n^T G_X(i\omega)e_n) &= 2\mathrm{Real}(e_n^T(i\omega\mathbf{I}-A)^{-1}\overline{\Sigma}e_n) \\ &\quad - 2\mathrm{Real}\left(e_n^T(i\omega\mathbf{I}-A)^{-1}B\Theta\mathbb{E}_\pi([\mathbb{R}_\mathbb{C}(i\omega)h(y)]h^T(y))(B\Theta)^T(i\omega\mathbf{I}+A^T)^{-1}e_n\right). \end{aligned} \tag{S54}$$

The first term on the l.h.s. can be simplified as in (S42). We now simplify the second term. Since the components of signal $(Y(t))_{t\geq 0}$ are independent, $\mathbb{E}_\pi([\mathbb{R}_\mathbb{C}(i\omega)h(y)]h^T(y))$ would be a $m \times m$ diagonal matrix, which we write as

$$\mathbb{E}_\pi([\mathbb{R}_\mathbb{C}(i\omega)h(y)]h^T(y)) = \mathrm{diag}(\psi_1(i\omega), \ldots, \psi_m(i\omega)),$$

where due to (S37) we have

$$2\mathrm{Real}(\psi_j(i\omega)) = S_{Y_j}(\omega) \quad \text{for} \quad j = 1, \ldots, m.$$

Letting $\alpha = (i\omega\mathbf{I}+A^T)^{-1}e_n$, we can write the second term on the l.h.s. of (S54) as

$$\begin{aligned} &- 2\mathrm{Real}\left(e_n^T(i\omega\mathbf{I}-A)^{-1}B\Theta\mathbb{E}_\pi([\mathbb{R}_\mathbb{C}(i\omega)h(y)]h^T(y))(B\Theta)^T(i\omega\mathbf{I}+A^T)^{-1}e_n\right) \\ &= 2\mathrm{Real}\left(\alpha^\dagger B\Theta\mathrm{diag}(\psi_1(i\omega), \ldots, \psi_m(i\omega))(B\Theta)^T\alpha\right) \\ &= 2\mathrm{Real}\left(\alpha^\dagger\left[\sum_{j=1}^m \theta_j^2 c_j c_j^T \psi_j(i\omega)\right]\alpha\right) \end{aligned}$$

$$
\begin{aligned}
&= 2\text{Real}\left(\sum_{j=1}^{m} \theta_j^2 |c_j^T \alpha|^2 \psi_j(i\omega)\right) \\
&= \sum_{j=1}^{m} \theta_j^2 |c_j^T \alpha|^2 2\text{Real}\left(\psi_j(i\omega)\right) \\
&= \sum_{j=1}^{m} \theta_j^2 |\alpha^T c_j|^2 S_{Y_j}(\omega).
\end{aligned}
$$

This completes the proof of this theorem. □

#### S2.4 Padé approximation of PSD for general nonlinear networks

In this section we describe our semi-analytic approach for obtaining a reliable estimate of the PSD for general stochastic reaction networks from a handful of stochastic trajectory simulations. We refer to this approach as Padé PSD as it is based on the theory of Padé approximations. We start by summarising the main idea behind the type of Padé approximants that we shall be constructing.

##### S2.4.1 Two-point Padé approximation

Suppose $\mathcal{G}(z)$ is some complex-analytic function, which has the following power-series expansion around $z = 0$

$$\mathcal{G}(z) = a_0 + a_1 z + a_2 z^2 + \dots \tag{S55}$$

and the following power-series expansion for large $|z|$

$$\mathcal{G}(z) = -\left(\frac{a_{-1}}{z} + \frac{a_{-2}}{z^2} + \frac{a_{-3}}{z^3} + \dots\right). \tag{S56}$$

Let $\mathbf{p} = (p_1, p_2)$ be a couple of nonnegative integers, such that their sum is even and let

$$p = \frac{p_1 + p_2}{2}. \tag{S57}$$

The goal of the two-point Padé approximation [18, 23] of order $\mathbf{p}$ is to construct a rational function of the form

$$\mathcal{G}_{\mathbf{p}}(z) := \frac{\kappa_0 + \kappa_1 z + \cdots + \kappa_{p-1} z^{p-1}}{\beta_0 + \beta_1 z + \cdots + \beta_{p-1} z^{p-1} + z^p}, \tag{S58}$$

such that its power series at $z = 0$ agrees with the first $p_1$ terms in (S55) and its power series in $1/z$ agrees with the first $p_2$ terms in (S56). In other words, $\mathcal{G}_{\mathbf{p}}(z)$ serves as an accurate approximation to $\mathcal{G}(z)$ both near $z = 0$ and near the region $|z| = \infty$.

The order $\mathbf{p}$ Padé approximant $\mathcal{G}_{\mathbf{p}}(z)$ can be constructed by computing a ratio of the determinant of certain matrices which are defined by the power series coefficients (see Section 4 in the main paper). Note that the power series coefficients $a_j$ in (S55) are given by

$$a_j = \frac{1}{j!} \frac{\partial}{\partial z^j} \mathcal{G}(z) \bigg|_{z=0} \quad \text{for} \quad j = 0, 1, \dots, (p_1 - 1). \tag{S59}$$

To obtain the coefficients in (S56), we first transform the function $\mathcal{G}(z)$ as

$$\mathcal{H}(z) = \frac{1}{z}\mathcal{G}\left(\frac{1}{z}\right). \tag{S60}$$

The power series of this function around $z = 0$ is given by

$$\mathcal{H}(z) = -(a_{-1} + a_{-2}z + a_{-3}z^2 + \dots)$$

and hence each $a_{-j}$ can be identified as

$$a_{-j} = -\frac{1}{(j-1)!}\frac{\partial}{\partial z^{j-1}}\mathcal{H}(z)\bigg|_{z=0} \quad \text{for} \quad j = 1, 2, \dots, p_2. \tag{S61}$$

##### S2.4.2 Estimating $G(z)$ via two-point Padé approximation

We now explain why a rational function of the form (S58) may be a good approximation of the function $G(z)$, defined by (S36), that characterises the output PSD (see Section S2.2). Since the resolvent operator is complex-analytic, the function $G(z)$ is also complex-analytic and so it suffices to estimate this function on the positive real line. We fix $z$ as some positive real number $s$. We shall assume in this section that function $f$, defined by (S35) can be expressed as (S33) and hence from (S34) we obtain

$$G(s) = \sum_{j=1}^{\infty} \frac{\alpha_j}{s - \sigma_j}, \tag{S62}$$

where $\alpha_j = c_j \mathbb{E}_\pi(f\phi_j) = c_j\langle f, \phi_j\rangle_{\mathcal{L}_2(\mathcal{E})}$. On the Hilbert space $\mathcal{L}_2(\mathcal{E})$ we can view $\alpha_j$ as the component of the function $f$ in the direction specified by the eigenfunction $\phi_j$. For a positive integer $p$, if we only keep the first $p$ terms in the series (S62) then we obtain a rational function of the form (S58) with numerator of degree $(p-1)$ and denominator of degree $p$. For $p$ large enough we can expect this approximation to be good due to (S32). Our examples how that often even a lower order approximation works very well for the purpose of PSD estimation.

Fix an order $\mathbf{p} = (p_1, p_2)$ given by a couple of nonnegative integers summing to an even number and let $p$ is given by (S57). We now discuss how we can construct the order $\mathbf{p}$ Padé approximant $G_{\mathbf{p}}(s)$ of the function $G(s)$, using the two-point Padé approximation method. First we select a finite positive number $s_0$ and define the translated function $\mathcal{G}(s)$ as

$$\mathcal{G}(s) = G(s + s_0). \tag{S63}$$

If we construct the order $\mathbf{p}$ approximant $\mathcal{G}_{\mathbf{p}}(s)$ for this function, and then the $G_{\mathbf{p}}(s)$ is simply given by

$$G_{\mathbf{p}}(s) = \mathcal{G}_{\mathbf{p}}(s - s_0). \tag{S64}$$

As discussed in Section S2.4.1, in order to construct $\mathcal{G}_{\mathbf{p}}(s)$ we will need to estimate the power series coefficients in (S55) and (S56). Observe that for the first power series, the coefficients (S59) can be expressed in terms of the derivatives of the function $G(s)$ as $s = s_0$

$$a_j = \frac{1}{j!}\frac{\partial}{\partial s^j}\mathcal{G}(s)\bigg|_{s=0} = \frac{1}{j!}\frac{\partial}{\partial s^j}G(s)\bigg|_{s=s_0} \quad \text{for} \quad j = 0, 1, \dots, (p_1 - 1). \tag{S65}$$

Note that

$$\begin{aligned}\frac{1}{j!}\frac{\partial}{\partial s^j}G(s) &= \frac{1}{j!}\frac{\partial}{\partial s^j}\mathbb{E}_\pi\left(f\mathbb{R}\left(s\right)f\right)\\ &= \frac{1}{j!}\frac{\partial}{\partial s^j}\mathbb{E}_\pi\left(f\int_0^\infty e^{-ts}\mathbb{T}(t)f dt\right)\\ &= \frac{(-1)^j}{j!}\mathbb{E}_\pi\left(f\int_0^\infty t^j e^{-ts}\mathbb{T}(t)f dt\right)\end{aligned}$$

Setting $s = s_0$ and substituting this relation in (S65) we obtain

$$a_j = \frac{1}{j!}\frac{\partial}{\partial s^j}G(s)\bigg|_{s=s_0} = \frac{(-1)^j}{j!}\mathbb{E}_\pi\left(f\int_0^\infty t^j e^{-ts_0}\mathbb{T}(t)f dt\right) \quad \text{for} \quad j = 0, 1, \ldots, (p_1 - 1). \tag{S66}$$

To identify coefficients (S61) of the second power series, we transform the function $\mathcal{G}(s)$ as

$$\begin{aligned}\mathcal{H}(s) = \frac{1}{s}\mathcal{G}\left(\frac{1}{s}\right) &= \frac{1}{s}G\left(s_0 + \frac{1}{s}\right)\\ &= \frac{1}{s}\mathbb{E}_\pi\left(f\mathbb{R}\left(s_0 + \frac{1}{s}\right)f\right)\\ &= \frac{1}{s}\mathbb{E}_\pi\left(f\int_0^\infty e^{-t\left(s_0+\frac{1}{s}\right)}\mathbb{T}(t)f dt\right)\\ &= \mathbb{E}_\pi\left(f\int_0^\infty e^{-t(s_0 s+1)}\mathbb{T}\left(ts\right)f dt\right),\end{aligned} \tag{S67}$$

where the last relation follows from the change of integration variable $t \mapsto ts$. Due to Dynkin's Theorem (see [9]), for any nonnegative integer $j$, we have

$$\frac{\partial^j}{\partial s^j}\mathbb{T}(ts)f = t^j\mathbb{A}^j\mathbb{T}(ts)f.$$

Applying the generalised product rule we obtain

$$\frac{d^j}{ds^j}\mathcal{H}(s) = \sum_{m=0}^{j}(-s_0)^{j-m}\binom{j}{m}\mathbb{E}_\pi\left(f\int_0^\infty e^{-t(s_0 s+1)}t^j\mathbb{A}^m\mathbb{T}\left(ts\right)f dt\right),$$

where

$$\binom{j}{m} = \frac{j!}{m!(j-m)!}$$

denotes the binomial coefficient. In order to identify the coefficients (S61) we now set $s = 0$. As $\mathbb{T}(0)$ is just the identity operator $\mathbf{I}$ and $\int_0^\infty e^{-t}t^j dt = j!$, we arrive at the following expression

$$a_{-j} = -\frac{1}{(j-1)!}\frac{\partial}{\partial s^{j-1}}\mathcal{H}(s)\bigg|_{s=0}$$

$$= -\sum_{m=0}^{j-1}(-s_0)^{j-m-1}\binom{j-1}{m}\mathbb{E}_\pi\left(f\mathbb{A}^m f\right) \quad \text{for} \quad j = 1,2,\dots,p_2. \tag{S68}$$

Henceforth we define the $j$-th Padé derivative at $s_0$ as

$$D_j^{(s_0)} := \mathbb{E}_\pi\left(f\int_0^\infty \frac{t^j}{j!}e^{-ts_0}\mathbb{T}(t)f dt\right) \tag{S69}$$

and the $j$-th Padé derivative at $\infty$ as

$$D_j^{(\infty)} := \mathbb{E}_\pi\left(f\mathbb{A}^j f\right). \tag{S70}$$

Then the power series coefficients are given by

$$a_j := \begin{cases} -\sum_{m=0}^{|j|-1}(-s_0)^{|j|-m-1}\binom{|j|-1}{m}D_m^{(\infty)} & \text{if } j = -1,-2,\dots,-p_2 \\ (-1)^j D_j^{(s_0)} & \text{if } j = 0,1,\dots,(p_1-1). \end{cases} \tag{S71}$$

With these coefficients the two-point Padé approximant $\mathcal{G}_{\mathbf{p}}(s)$ (see (S58)) of order $\mathbf{p}$ can be obtained for function $\mathcal{G}(s)$ which can then be translated into such an approximant $G_{\mathbf{p}}$ for function $G(s)$ via (S64). Note that if $p_1 = 0$ then we do not need the Padé derivatives at $s_0$ and in this case

$$a_{-j} = -D_{j-1}^{(\infty)} \qquad \text{for} \qquad j = 1,2,\dots,p_2.$$

We next turn our attention to how the steady-state expectations defining the Padé derivatives can be efficiently and accurately estimated.

##### S2.4.3 Design of covariates for the estimation of Padé derivatives at $\infty$

Recall that the Padé derivative $D_m^{(\infty)}$ can be estimated by the Monte Carlo estimator (22) stated in the main paper. We now discuss how we can design covariates to improve the estimation efficiency. For a fixed number of trajectories $Q$, this efficiency crucially depends on the time $T_f$ it takes for the integrals estimating $D_m^{(\infty)}$ (see (22)) to converge to their steady-state values. Ideally we would like this convergence to happen for low values of $T_f$ and for this we design *covariates* that speed up the convergence in (22). Use of covariates is a common variance reduction technique for Monte Carlo estimators (see [2]) and we now describe how we employ these in our setting. Letting $\Psi_m = f\mathbb{A}^m f$ we have $D_m^{(\infty)} = \mathbb{E}_\pi(\Psi_m)$ and as we mentioned in Section S1.2, the convergence of the estimator (22) with time $T_f$ is captured by the *Time Average Variance Constant* (TAVC) defined by (S15). A good covariate for the estimator (22) would be a function $\Phi_m : \mathcal{E} \to \mathbb{R}$ such that

$$\mathbb{E}_\pi(\Phi_m) = 0 \tag{S72}$$

and

$$\text{TAVC}(\Psi_m + \Phi_m) < \text{TAVC}(\Psi_m). \tag{S73}$$

The first condition (S72) ensures that adding $\Phi_m$ to $\Psi_m$ does not change the stationary expectation $D_m^{(\infty)}$ to be estimated (i.e. $D_m^{(\infty)} = \mathbb{E}_\pi(\Psi_m + \Phi_m)$) and the second condition (S73) ensures that the estimator (22) with $\Psi_m$ replaced by $(\Psi_m+\Phi_m)$ has better convergence properties.

Observe that if $\Phi_m$ is in the range of generator $\mathbb{A}$ (i.e. $\mathbb{A}\phi_m = \Phi_m$ for some function $\phi_m$) then (S72) is automatically satisfied due to (S12). We shall use this fact to come up with suitable covariates $\Phi_m$ for $m = 0, \ldots, (p_2 - 1)$. Applying the product-rule (S6) on the product of output function $f$ with itself we get

$$\mathbb{A}f^2(x) = 2f(x)\mathbb{A}f(x) + \sum_{k=1}^{K} \lambda_k(x)(\Delta_k f(x))^2. \tag{S74}$$

Since $\Psi_1 = f\mathbb{A}f$ we can write this relation as

$$\Psi_1 - \frac{1}{2}\mathbb{A}f^2(x) = -\frac{1}{2}\sum_{k=1}^{K} \lambda_k(x)(\Delta_k f(x))^2.$$

This suggests that we can choose the covariate for $m = 1$ as $\Phi_1 = -\frac{1}{2}\mathbb{A}f^2(x)$ and obtain

$$\Psi_1^{(c)}(x) := \Psi_1(x) + \Phi_1(x) = -\frac{1}{2}\gamma_{00}(x), \tag{S75}$$

where $\gamma_{jl}(x)$ is defined by (24). We find the covariate for $m = 2$ by applying the generator to equation (S74) and simplifying with the help of product-rule (S6). This yields

$$\mathbb{A}^2 f^2(x) = 2f(x)\mathbb{A}^2 f(x) + 2\left(\mathbb{A}f(x)\right)^2 + \mathbb{A}\gamma_{00}(x) + 2\gamma_{01}(x) \tag{S76}$$

and we choose the covariate for $m = 2$ as

$$\Phi_2(x) = -\frac{1}{2}\left[\mathbb{A}^2 f^2(x) - \mathbb{A}\gamma_{00}(x)\right] = -\left[\frac{1}{2}\mathbb{A}^2 f^2(x) + \mathbb{A}\Psi_1^{(c)}(x)\right].$$

Adding this covariate to $\Psi_2(x)$ we get

$$\Psi_2^{(c)}(x) := \Psi_2(x) + \Phi_2(x) = -\left[\left(\mathbb{A}f(x)\right)^2 + \gamma_{01}(x)\right]. \tag{S77}$$

Continuing this way and applying the generator $\mathbb{A}$ iteratively to function $f^2$ we can find the covariate $\Phi_m$ for each $m$, and adding this covariate to $\Psi_m(x)$ we obtain expressions (23) given in the main paper. The exact expression for the covariate $\Phi_m$ is not needed, as once we have $\Psi_m^{(c)}$ we can estimate $D_m^{(\infty)} = \mathbb{E}_\pi(\Psi_m^{(c)})$ via the Monte Carlo estimator (25). We find that in practice using covariates immensely helps in decreasing the time $T_f$ required for estimator integral to converge to its steady-state value.

### S3 Algorithms for Padé PSD

In this section we provide detailed algorithms for the implementation of Padé PSD. Recall from Section 4.4 in the main paper that Padé PSD requires simulation of a suitably

**Algorithm 1** Computes the next time increment $\Delta t$ and the next reaction number $k$ at a given state $x$ which is the species copy-number vector.

**function** SSA-STEP($x$)
    Set $r_1 = \text{rand}()$ , $r_2 = \text{rand}()$ and $k = 0$
    Set $\lambda_0(x) := \sum_{k=1}^{K} \lambda_k(x) + s_0 + \sum_{r=1}^{R} s_r = \sum_{k=1}^{K+1+R} \lambda_k(x)$
    Calculate $\Delta t = -\log(r_1)/\lambda_0(x)$
    Set $S = 0$
    **while** $S < r_2$ **do**
        Update $k \leftarrow k + 1$
        Update $S \leftarrow S + \lambda_k(x)/\lambda_0(x)$
    **end while**
**return** $(\Delta t, k)$.
**end function**

constructed augmented CTMC by extending the classical Gillespie's Stochastic Simulation Algorithm [11]. For this the next reaction event is generated by Algorithm 1 which relies on the function *rand()* that produces an independent sample from the uniform distribution on $[0, 1]$.

To estimate the Padé derivative $D_m^{(\infty)}$ for $m \geq 1$ we shall use the estimator (25) that employs covariates to reduce the convergence time. Note that this estimator requires evaluation of the function $\psi_m^{(c)}(x)$ (see (23)) at each state $x$ visited by the CTMC. This function is defined in terms of the output function

$$f(x) = x_n - \mathbb{E}_\pi(X_m)$$

and since $\psi_m^{(c)}(x)$ only depends on differences[3] of function $f$, we can drop the constant term $\mathbb{E}_\pi(X_m)$ and just work with the output function $f(x) = x_n$. The estimators (27) and (32), can be easily adapted to work with this simpler output function. For example, (27) changes to

$$\widehat{D}_m^{(s_0)} = \frac{1}{s_0^{m+1} Q(T_f - T_c)} \sum_{q=1}^{Q} \int_{T_c}^{T_f} f(X_q(t)) Y_{m+1}(t) dt - \frac{1}{s_0^{m+1}} \frac{1}{Q} \sum_{q=1}^{Q} \left[ \frac{1}{(T_f - T_c)} \int_{T_c}^{T_f} f(X_q(t)) dt \right]^2 .$$

In order to compute $\psi_m^{(c)}(x)$ we need a method to compute $\mathbb{A}^m f(x)$, which is generator $\mathbb{A}$ applied $m$ times to the output function $f$ and then evaluated at state $x$. A simple recursive method for this task is presented as Algorithm 2. Using this method we can compute the function $\gamma_{jl}(x)$ (see (24)) as described in Algorithm 3. The function $\psi_m^{(c)}(x)$ can then be evaluated at each state $x$ visited by the CTMC using Algorithm 4.

Upon simulating the augmented CTMC with extended SSA using Algorithm 1, we obtain a continuous-time trajectory $(\mathcal{X}_q(t))_{t \geq 0}$ over the time-interval $[0, T_f]$. We ignore the portion

[3]Note that generator $\mathbb{A}$ given by (S3) only depends on differences of function $f$.

of the trajectory in the cutoff period $[0, T_c]$ and discrete-sample the rest with time-step $h = (T_f - T_c)/N_f$, where $N_f$ is some large positive integer denoting the number of discrete time-steps. Letting

$$t_j = T_c + (j-1)h \quad \text{for} \quad j = 1, \dots, N_f,$$

we obtain the discretised version of $(\mathcal{X}_q(t))_{t \geq 0}$ as the time-series $\widehat{\mathcal{X}}_q = (\mathcal{X}_q(t_1), \dots, \mathcal{X}_q(t_{N_f}))$. Generating such a time-series for each $q = 1, \dots, Q$, the time-integrals in the Monte Carlo estimators (25), (27) and (32) can be replaced by the finite-sum

$$\frac{1}{(T_f - T_c)} \int_{T_c}^{T_f} \mathcal{J}(\mathcal{X}_q(t))dt \mapsto \frac{1}{N_f} \sum_{j=1}^{N_f} \mathcal{J}(\mathcal{X}_q(t_j)),$$

where $\mathcal{J}$ is an arbitrary function of the state. Note that this discrete-sampling *does not add any bias* to the estimated integrals in the ergodic limit $T_f, N_f \to \infty$ with fixed time-step $h$. Algorithm 5 takes these discrete-sampled augmented CTMC trajectories as inputs and computes the Monte Carlo estimators for all the required quantities. The standard deviations of these estimates can also be computed to adjudge their accuracy (not shown in Algorithm 5).

**Algorithm 2** Recursively computes the value $\mathbb{A}^m f(x)$ for any state $x$ and any nonnegative integer $m$.

**function** RecursiveGenerator($x$,$m$)
  **if** $m = 0$ **then return** $f(x) = x_n$
  **else**
    Compute $\lambda_0(x) := \sum_{k=1}^{K} \lambda_k(x)$
    Set `value` $= -\lambda_0(x) \times$ RecursiveGenerator$(x, m-1)$
    **for** $k = 1, \dots, K$ **do**
      Update `value` $\leftarrow$ `value` $+ \lambda_k(x) \times$ RecursiveGenerator$(x + \zeta_k, m-1)$
    **end for**
  **end if**
**return** `value`
**end function**

One issue that arises in the estimation of the Padé derivatives $D_m^{(\infty)}$ is that each call to function RecursiveGenerator($x$,$m$) (to compute $\mathbb{A}^m f(x)$) requires $(K+1)$ calls to the same function with argument $m$ reduced by one (see Algorithm 2). Here $K$ is the number of reactions and the calls are generated at state $x$ and all its adjacent states $(x + \zeta_k)$ for $k = 1, \dots, K$. Consequently the number of operations needed to evaluate RecursiveGenerator($x$,$m$) is proportional to $(K+1)^m$, which grows very quickly with $m$ if $K$ is large. To circumvent this problem we exploit the fact that ergodic Markov chains visit the same set of states again and again. Hence if we have computed the value RecursiveGenerator($x$,$m$) when a CTMC was at state $x$, then we can simply reuse this value without having to compute it again, for any of the simulated CTMCs. Therefore if we can intelligently store the values $\mathbb{A}^m f(x)$ generated by this function, and quickly retrieve them as needed, then it provides a way to leverage the vast memory resources in modern computers for gaining

**Algorithm 3** Computes the value $\gamma_{jl}(x)$ (see (24)) for any state $x$ and any nonnegative integers $j, l$.

```
function COMPUTE-GAMMA(x,j,l)
    Set f_1 = RECURSIVEGENERATOR(x, j) and f_2 = RECURSIVEGENERATOR(x, l)
    Set value = 0
    for k = 1, ..., K do
        Set f_3 = RECURSIVEGENERATOR(x + ζ_k, j)
        Set f_4 = RECURSIVEGENERATOR(x + ζ_k, l)
        Update value ← value + λ_k(x) × (f_3 − f_1) × (f_4 − f_2)
    end for
return value
end function
```

computational efficiency. We do this with the `dictionary` data structure in Python that is essentially a fancier version of the *Hash Table* [5]. In our implementation, such a `dictionary` was used for storing values $\mathbb{A}^m f(x)$ according to their 'key' given by the state vector $x$. The function RECURSIVEGENERATOR(x,m) was updated to first look for the value $\mathbb{A}^m f(x)$ in the `dictionary`, and if it is not found then this value is recursively computed and stored in the `dictionary` with key $x$. This storage and retrieval method greatly enhanced the computational speed of the Padé PSD method.

### S4 Numerical Examples

In this section we provide details on the examples considered in the main paper to illustrate the developed results. Throughout the paper, Padé PSD is applied with $Q = 10$ CTMC trajectories simulated till time $T_f = 10'100$ and we set the cut-off time as $T_c = 100$ in the time units relevant to the example. For the averaged periodogram approach for PSD estimation, the ten trajectories were discrete-sampled with a time-step of 1 and then their periodogram was computed. The order $\mathbf{p} = (p_1, p_2)$ of Padé approximant varies for each example but $s_0$ is fixed at 1.0. The estimated approximant is validated with four test values $\mathbf{s} = (0.5, 0.75, 1.25, 1.5)$, as discussed in Section 4.3, and the result is plotted to adjudge the accuracy of the approximant. We provide expressions for the Padé approximants and the estimated PSD for all the examples.

#### S4.1 Linear Networks

We validated our Padé PSD method with linear networks where analytical expressions for the PSD are available. Below we provide details on these examples.

**Algorithm 4** Computes the value $\psi_m^{(c)}(x)$ (see (23)) for any state $x$ and any nonnegative integer $m$.

```
function Compute-Psi(x,m)
    Set value = 0
    if m if even then
        Set q = m/2
        for k = 1, ..., (q − 1) do
            Set f_1 = RecursiveGenerator(x, k)
            Set f_2 = RecursiveGenerator(x, m − k)
            Update value ← value + binom(m, k) × f_1 × f_2
            Update value ← value + binom(m−1, k) × Compute-Gamma(x, k, m − 1 − k)
        end for
        Set f_3 = RecursiveGenerator(x, q)
        Update value ← value + Compute-Gamma(x, 0, m − 1) + (1/2) binom(m, q) × f_3^2
    else
        Set q = (m − 1)/2
        for k = 1, ..., q do
            Set f_1 = RecursiveGenerator(x, k)
            Set f_2 = RecursiveGenerator(x, m − k)
            Update value ← value + binom(m, k) × f_1 × f_2
            Update value ← value + binom(m−1, k−1) × Compute-Gamma(x, k − 1, m − k)
        end for
        Update value ← value + (1/2) binom(m−1, q) × Compute-Gamma(x, q, q)
    end if
return value
end function
```

##### S4.1.1 Gene Expression Network

The gene expression network (see Figure 2(A)) involves two species, namely the mRNA ($\mathbf{X_1}$) and the protein ($\mathbf{X_2}$), and the following four reactions:

$$\emptyset \xrightarrow{k_r} \mathbf{X_1} \quad \text{(mRNA transcription)}$$
$$\mathbf{X_1} \xrightarrow{k_p} \mathbf{X_1} + \mathbf{X_2} \quad \text{(protein translation)}$$
$$\mathbf{X_1} \xrightarrow{\gamma_r} \emptyset \quad \text{(mRNA degradation)}$$
$$\mathbf{X_2} \xrightarrow{\gamma_p} \emptyset \quad \text{(protein degradation).}$$

Here rate constants are indicated above the reaction arrows and mass-action kinetics is assumed.

In the notation in Section S2.3, the $2 \times 2$ matrix $A$ and the $2 \times 1$ vector $b$ are given by

$$A = \begin{bmatrix} -\gamma_r & 0 \\ k_p & -\gamma_p \end{bmatrix} \quad \text{and} \quad b = \begin{bmatrix} k_r \\ 0 \end{bmatrix}$$

and the stationary mean (S39) can be computed as

$$\overline{x}_1 = \mathbb{E}_\pi(X_1) = \frac{k_r}{\gamma_r} \qquad \text{and} \qquad \overline{x}_2 = \mathbb{E}_\pi(X_2) = \frac{k_r k_p}{\gamma_r \gamma_p}.$$

On solving the Lyapunov equation (S40) we obtain the second column of the stationary variance-covariance $\Sigma$ as

$$\begin{aligned} \Sigma_{21} = \mathrm{Cov}_\pi(X_1, X_2) &= \frac{k_p k_r}{\gamma_r(\gamma_r + \gamma_p)} \\ \text{and} \quad \Sigma_{22} = \mathrm{Var}_\pi(X_2) &= \frac{k_p k_r(\gamma_r + \gamma_p + k_p)}{\gamma_r \gamma_p(\gamma_r + \gamma_p)}. \end{aligned}$$

Using formula (12) in the main paper and simplifying, we arrive at the following expression for the PSD for the output trajectory

$$S_{X_2}(\omega) = \frac{2k_r k_p}{\gamma_r}\left[\frac{\gamma_r^2 + k_p\gamma_r + \omega^2}{\gamma_r^2\gamma_p^2 + \omega^2(\gamma_r^2 + \gamma_p^2) + \omega^4}\right]. \tag{S78}$$

This is equivalent to the expression (36) derived in the main paper using Theorem 3.1. For the numerical results shown in Figure 2(A) the parameters for the gene expression network are

$$k_r = 10 \text{ min}^{-1}, \quad k_p = 2 \text{ min}^{-1}, \quad \gamma_r = 1 \text{ min}^{-1} \quad \text{and} \quad \gamma_p = 0.5 \text{ min}^{-1}$$

and for these parameters the exact PSD (S78) evaluates to

$$S_{X_2}(\omega) = \frac{40\omega^2 + 120}{\omega^4 + 1.25\omega^2 + 0.25}. \tag{S79}$$

With order $\mathbf{p} = (0, 4)$ the Padé approximant produced by our method is

$$G_{\mathbf{p}}(s) = \frac{92.8379s + 118.9966}{s^2 + 1.497s + 0.5021}$$

and this yields the following expression for the PSD (see (S37))

$$\widehat{S}_{X_2}(\omega) = \frac{39.9629\omega^2 + 119.4918}{\omega^4 + 1.2368\omega^2 + 0.2521}.$$

Notice that this expression is very close to the exact PSD expression (S79). We validated the estimated Padé approximant $G_{\mathbf{p}}(s)$ and the result is shown in Figure S1(A).

##### S4.1.2 RNA Splicing Network

The RNA Splicing network (see Figure 2(B)) consists of four species, which are, inactive gene-transcript ($\mathbf{X_1}$), active gene-transcript ($\mathbf{X_2}$), unspliced mRNA ($\mathbf{X_3}$) and spliced mRNAs ($\mathbf{X_4}$). These species undergo the following five reactions:

$$\mathbf{X_1} \xrightarrow{k_{\mathrm{on}}} \mathbf{X_2} \quad \text{(gene activation)}$$

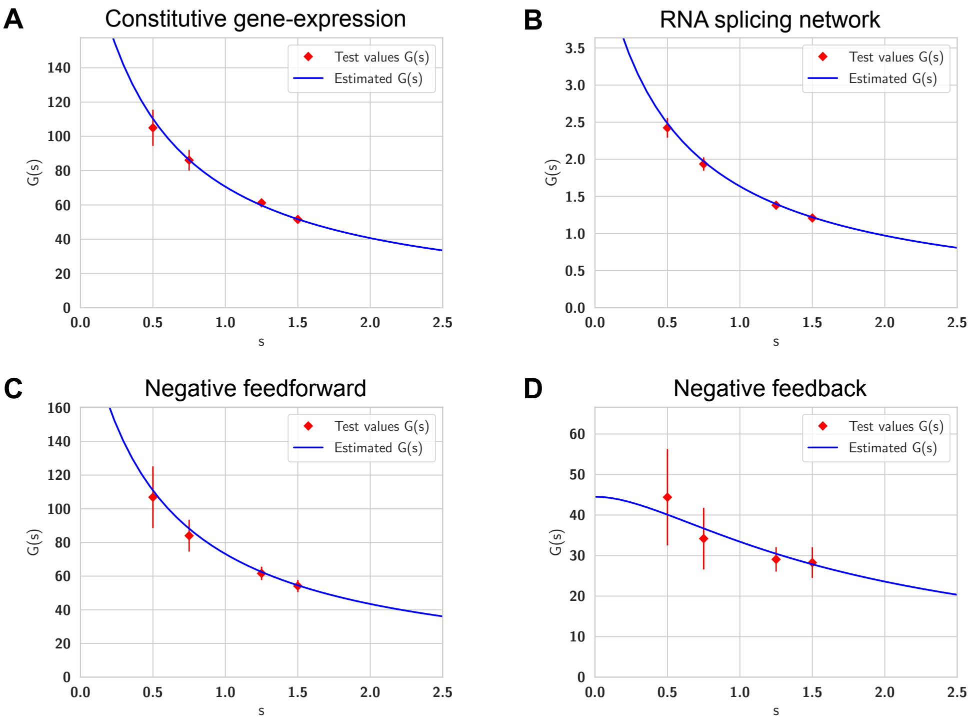

Supplementary Figure S1: Validation of the estimated Padé approximant $G_{\mathbf{p}}(s)$ for linear networks. Direct estimates for $G(s)$ are plotted in red for four test values $\mathbf{s} = (0.5, 0.75, 1.25, 1.5)$. The point denotes the estimator mean and the error bar denotes the symmetric one standard deviation interval. The blue curve denotes the estimated Padé approximant.

$$\mathbf{X_2} \xrightarrow{k_{\text{off}}} \mathbf{X_1} \quad \text{(gene inactivation)}$$
$$\mathbf{X_1} \xrightarrow{\alpha_{\text{off}}} \mathbf{X_1} + \mathbf{X_3} \quad \text{(inactive gene transcription)}$$
$$\mathbf{X_2} \xrightarrow{\alpha_{\text{on}}} \mathbf{X_2} + \mathbf{X_3} \quad \text{(active gene transcription)}$$
$$\mathbf{X_3} \xrightarrow{\beta} \mathbf{X_4} \quad \text{(splicing)}$$
$$\mathbf{X_4} \xrightarrow{\gamma} \emptyset \quad \text{(degradation)}.$$

Here rate constants are indicated above the reaction arrows and mass-action kinetics is assumed. Note that the number of inactive gene-transcripts is $X_1 = 1 - X_2$ where $X_2 \in \{0, 1\}$ is the number of active gene transcripts. Hence we do not need to model the copy-number dynamics of $\mathbf{X_1}$ explicitly. For the three other species, the $3 \times 3$ matrix $A$ and the $3 \times 1$ vector $b$ are given by

$$A = \begin{bmatrix} -(k_{\text{on}} + k_{\text{off}}) & 0 & 0 \\ (\alpha_{\text{on}} - \alpha_{\text{off}}) & -\beta & 0 \\ 0 & \beta & -\gamma \end{bmatrix} \quad \text{and} \quad b = \begin{bmatrix} k_{\text{on}} \\ \alpha_{\text{off}} \\ 0 \end{bmatrix}$$

and the stationary mean (S39) can be computed as

$$\overline{x}_2 = \mathbb{E}_\pi(X_2) = \frac{k_{\text{on}}}{k_{\text{on}} + k_{\text{off}}}, \quad \overline{x}_3 = \mathbb{E}_\pi(X_3) = \frac{\alpha_{\text{on}} k_{\text{on}} + \alpha_{\text{off}} k_{\text{off}}}{\beta(k_{\text{on}} + k_{\text{off}})} \quad \text{and} \quad \overline{x}_4 = \mathbb{E}_\pi(X_4) = \frac{\beta}{\gamma}\overline{x}_3.$$

On solving the Lyapunov equation (S40) we obtain that the last column of the stationary variance-covariance $\Sigma$ is

$$\begin{aligned} \Sigma_{24} &= \text{Cov}_\pi(X_2, X_4) = \frac{\beta k_{\text{on}} k_{\text{off}}(\alpha_{\text{on}} - \alpha_{\text{off}})}{(k_{\text{on}} + k_{\text{off}})^2(\beta + k_{\text{on}} + k_{\text{off}})(\gamma + k_{\text{on}} + k_{\text{off}})}, \\ \Sigma_{34} &= \text{Cov}_\pi(X_3, X_4) = \frac{(\alpha_{\text{on}} - \alpha_{\text{off}})(\beta + \gamma + k_{\text{on}} + k_{\text{off}})}{\beta(\beta + \gamma)}\Sigma_{24} \\ \text{and} \quad \Sigma_{44} &= \text{Var}_\pi(X_4) = \overline{x}_4 + \frac{\beta}{\gamma}\Sigma_{34}. \end{aligned}$$

Using (12) in the main paper and simplifying, we arrive at expression (38) in the main paper which was derived using Theorem 3.1. For the numerical results shown in Figure 2(B) the parameters for the RNA Splicing network are

$$\begin{aligned} &k_{\text{on}} = 1 \text{ min}^{-1}, \quad k_{\text{off}} = 3 \text{ min}^{-1}, \quad \alpha_{\text{on}} = 3 \text{ min}^{-1}, \quad \alpha_{\text{off}} = 0.2 \text{ min}^{-1}, \\ &\beta = 2 \text{ min}^{-1} \quad \text{and} \quad \gamma = 0.5 \text{ min}^{-1}. \end{aligned}$$

With these parameters the exact PSD expression becomes

$$S_{X_2}(\omega) = \frac{1.8\omega^4 + 36\omega^2 + 162.234}{\omega^6 + 20.25\omega^4 + 69\omega^2 + 16}. \tag{S80}$$

The order $\mathbf{p} = (2, 4)$ Padé approximant produced by our method is

$$G_{\mathbf{p}}(s) = \frac{2.334s^2 - 19.3391s - 15.978}{s^3 - 7.9023s^2 - 9.9196s - 3.3407}$$

and this yields the following expression for the PSD (see (S37))

$$\widehat{S}_{X_2}(\omega) = \frac{1.7898\omega^4 + 146.7412\omega^2 + 106.7541}{\omega^6 + 82.2862\omega^4 + 45.6011\omega^2 + 11.16}.$$

We validated the estimated Padé approximant $G_{\mathbf{p}}(s)$ and the result is shown in Figure S1(B). Note that even though this estimated PSD $\widehat{S}_{X_2}(\omega)$ appears to be quite different from the exact PSD (S80), their graphs for $\omega \in [0, \infty)$ are almost identical, as shown in Figure 2(B).

##### S4.1.3 Differentiation between adapting circuit topologies

We consider the simple three-node IFF and NFB topologies depicted in Figure 2(C,D) with stochastic kinetics. We use the results in Section S2.3 to provide analytical expressions for the PSD under the assumption that we can linearise the propensity functions capturing the repression mechanism. These analytical expressions inform us about qualitative differences in the PSD emanating from IFF and NFB topologies that are *structural*, in the sense that these differences hold regardless of the choice of reaction rate parameters. This shows that in the presence of stochastic dynamics, the PSD of single-cell trajectories serves as a key "response signature" that can differentiate between adapting circuit topologies.

We begin our analysis with the IFF topology whose reactions are given by

$$\mathbf{I} \xrightarrow{k_c I_0} \mathbf{I} + \mathbf{C}, \quad \mathbf{I} \xrightarrow{k_o I_0} \mathbf{I} + \mathbf{O}, \quad \mathbf{C} \xrightarrow{F_f(x_c)} \mathbf{C} + \mathbf{O}, \quad \mathbf{C} \xrightarrow{\gamma_c x_c} \emptyset \quad \text{and} \quad \mathbf{O} \xrightarrow{\gamma_o x_o} \emptyset, \tag{S81}$$

The reaction rates are stated above the reaction arrow, and $x_c$ (resp. $x_o$) denotes the copy-number of species $\mathbf{C}$ (resp. $\mathbf{O}$). The input abundance level $I_0$ is assumed to be constant, and $F_f(x_c)$ is a monotonically decreasing function representing the repression in the production of output species $\mathbf{O}$ by the controller species $\mathbf{C}$.

For our analysis we would need to *linearize* the function $F_f(x_c)$ as

$$F_f(x_c) = \beta_0 - \beta_{\text{ff}} x_c,$$

where $\beta_0$ and $\beta_{\text{ff}}$ are positive constants denoting the *basal* production rate and the strength of the incoherent feedforward respectively. Following the notation in Section S2.3, the $2 \times 2$ matrix $A$ and the $2 \times 1$ vector $b$ are given by

$$A = \begin{bmatrix} -\gamma_c & 0 \\ -\beta_{\text{ff}} & -\gamma_o \end{bmatrix} \quad \text{and} \quad b = \begin{bmatrix} k_c I_0 \\ \beta_0 + k_o I_0 \end{bmatrix}.$$

Note that since matrix $A$ is lower-triangular, its eigenvalues are the diagonal entries $-\gamma_c$ and $-\gamma_o$ and hence matrix $A$ is Hurwitz-stable for any choice of parameters. The stationary mean (S39) can be computed as

$$\overline{x}_c = \mathbb{E}_\pi(C) = \frac{k_c I_0}{\gamma_c} \qquad \text{and} \qquad \overline{x}_o = \mathbb{E}_\pi(O) = \frac{k_o I_0 + \beta_0}{\gamma_o} - \frac{\beta_{\text{ff}} k_c I_0}{\gamma_c \gamma_o}.$$

Observe that if $\beta_{\text{ff}} \approx k_o \gamma_c / k_c$, then the mean output value $\overline{x}_o \approx \beta_0/\gamma_o$ becomes insensitive to the input abundance level $I_0$. This shows the adaptation property of the IFF network.

Upon solving the Lyapunov equation (S40) we find that the second column of the stationary variance-covariance $\Sigma$ is given by

$$\Sigma_{21} = \text{Cov}_\pi(C, O) = -\frac{\beta_{\text{ff}} k_c I_0}{\gamma_c(\gamma_c + \gamma_o)}$$
$$\text{and} \quad \Sigma_{22} = \text{Var}_\pi(O) = \frac{\beta_0 + k_o I_0}{\gamma_o} - \frac{\beta_{\text{ff}} k_c I_0}{\gamma_c \gamma_o} + \frac{\beta_{\text{ff}}^2 k_c I_0}{\gamma_c \gamma_o (\gamma_c + \gamma_o)}.$$

Using (12) in the main paper and simplifying, we obtain the following expression for the PSD for the output trajectory

$$S_O(\omega) = \frac{2}{\gamma_o^2 + \omega^2} \left[ \beta_0 + k_o I_0 - \frac{\beta_{\text{ff}} k_c I_0}{\gamma_c} + \frac{\beta_{\text{ff}}^2 k_c I_0}{\gamma_c^2 + \omega^2} \right]. \tag{S82}$$

As the PSD $S_O(\omega)$ is a monotonically decreasing function of $\omega$, it does not have a local maximum at any positive value of $\omega$ *regardless of the IFF network parameters.*

We now turn our attention to the negative feedback (NFB) network depicted in Figure 2. Here the reactions are given by

$$\mathbf{I} \xrightarrow{I_0 F_b(x_o)} \mathbf{I} + \mathbf{C}, \quad \mathbf{C} \xrightarrow{k_o x_c} \mathbf{C} + \mathbf{O}, \quad \mathbf{C} \xrightarrow{\gamma_c x_c} \emptyset \quad \text{and} \quad \mathbf{O} \xrightarrow{\gamma_o x_o} \emptyset, \tag{S83}$$

where $x_c$ (resp. $x_o$) denotes the copy-number of species $\mathbf{C}$ (resp. $\mathbf{O}$) and $F_b(x_o)$ is a monotonically decreasing function representing the repression of the production of the controller species $\mathbf{C}$ by the output species $\mathbf{O}$. We *linearize* the function $F_b(x_o)$ as

$$F_b(x_o) = \beta_0 - \beta_{\text{fb}} x_o,$$

where $\beta_0$ is the basal production rate and $\beta_{\text{fb}}$ is the feedback strength. The $2 \times 2$ matrix $A$ and the $2 \times 1$ vector $b$ are given by

$$A = \begin{bmatrix} -\gamma_c & -\beta_{\text{fb}} I_0 \\ k_o & -\gamma_o \end{bmatrix} \quad \text{and} \quad b = \begin{bmatrix} \beta_0 I_0 \\ 0 \end{bmatrix}$$

and the stationary mean (S39) can be computed as

$$\overline{x}_c = \mathbb{E}_\pi(C) = \frac{\gamma_o \beta_0 I_0}{\gamma_c \gamma_o + k_o \beta_{\text{fb}} I_0} \qquad \text{and} \qquad \overline{x}_o = \mathbb{E}_\pi(O) = \frac{k_o \beta_0 I_0}{\gamma_c \gamma_o + k_o \beta_{\text{fb}} I_0}.$$

Observe that if the input abundance level $I_0$ is high, then mean output value $\overline{x}_o \approx \beta_0/\beta_{\text{fb}}$ only depends on the feedback function $F_b$ and it is insensitive to $I_0$, thereby demonstrating the adaptation property. On solving the Lyapunov equation (S40) we obtain the second column of the stationary variance-covariance $\Sigma$ as

$$\Sigma_{21} = \text{Cov}_\pi(C, O) = \frac{\gamma_c \gamma_o k_o \beta_0 I_0 (\gamma_o - \beta_{\text{fb}} I_0)}{(\gamma_c + \gamma_o)(\gamma_c \gamma_o + k_o \beta_{\text{fb}} I_0)^2}$$
$$\text{and} \quad \Sigma_{22} = \text{Var}_\pi(O) = \frac{\gamma_o k_o \beta_0 I_0 (\gamma_c^2 + \gamma_c \gamma_o + \gamma_c k_o + k_o \beta_{\text{fb}} I_0)}{(\gamma_c + \gamma_o)(\gamma_c \gamma_o + k_o \beta_{\text{fb}} I_0)^2}.$$

Using (12) in the main paper and simplifying, we arrive at the following expression for the PSD for the output trajectory

$$S_O(\omega) = \frac{2\gamma_o k_o \beta_0 I_0}{\gamma_c \gamma_o + k_o \beta_{\text{fb}} I_0} \left[ \frac{\gamma_c^2 + k_o \gamma_c + \omega^2}{(\gamma_c \gamma_o + k_o \beta_{\text{fb}} I_0)^2 + \omega^2 (\gamma_c^2 + \gamma_o^2 - 2 k_o \beta_{\text{fb}} I_0) + \omega^4} \right]. \tag{S84}$$

The next proposition shows that the mapping $\omega \mapsto S_O(\omega)$ has a positive local maximum if and only if

$$\beta_{\text{fb}} I_0 > \frac{\gamma_c^4 + \gamma_c^3 k_o + \gamma_o^2 \gamma_c k_o}{k_o \left( \sqrt{(\gamma_c \gamma_o + \gamma_c^2 + k_o \gamma_c)^2 + (\gamma_c^4 + \gamma_c^3 k_o + \gamma_o^2 \gamma_c k_o)} + \gamma_c \gamma_o + \gamma_c^2 + k_o \gamma_c \right)}. \tag{S85}$$

Moreover if this holds then the positive local maximum exists uniquely (hence it is also the global maximum) and its location is given by

$$\omega_{\max} = \sqrt{\sqrt{k_o(\gamma_c^3 + k_o \gamma_c^2 + k_o^2 \beta_{\text{fb}}^2 I_0^2 - \gamma_o^2 \gamma_c + 2\gamma_c \gamma_o \beta_{\text{fb}} I_0 + 2\gamma_c^2 \beta_{\text{fb}} I_0 + 2 k_o \gamma_c \beta_{\text{fb}} I_0)} - \gamma_c \gamma_o - \gamma_c^2}. \tag{S86}$$

Condition (S85) shows that *regardless of the NFB network parameters*, the output trajectories will exhibit oscillation if the input abundance level $I_0$ is high enough. This differentiates it from the IFF circuit which never exhibits oscillations. Note that high $I_0$ is precisely the condition for NFB to show adaptation and hence imposing this requirement is not very restrictive.

**Proposition S4.1 (Condition for oscillation)** *The PSD $S_O(\omega)$ for the NFB network achieves a local maximum at a non-zero frequency if and only if* (S85) *holds. Such a maximum is also the global maximum for $\omega \in [0, \infty)$ and its location is given by* (S86)*.*

**Proof.** Note that the PSD (S84) can be expressed as

$$S_O(\omega) = \frac{2\gamma_o k_o \beta_0 I_0}{\gamma_c\gamma_o + k_o\beta_{\text{fb}} I_0} g(x)$$

where $x = \omega^2$ and

$$g(x) = \frac{a + x}{b + cx + x^2}$$

with $a = (\gamma_c^2 + k_o\gamma_c)$, $b = (\gamma_c\gamma_o + k_o\beta_{\text{fb}}I_0)^2$ and $c = (\gamma_c^2 + \gamma_o^2 - 2k_o\beta_{\text{fb}}I_0)$. The derivative of $g$ w.r.t. $x$ is given by

$$g'(x) = \frac{b - ac - 2ax - x^2}{(b + cx + x^2)^2}.$$

It is immediate that the only positive solution to $g'(x) = 0$ can be

$$x = -a + \sqrt{a^2 + b - ac} \tag{S87}$$

and this positive solution exists if and only if we have

$$b > ac. \tag{S88}$$

One can check that for the aforementioned values of $a, b$ and $c$, condition (S88) simplifies to (S85) and letting $\omega_{\max} = \sqrt{x}$, expression (S87) yields (S86). Since there is at most one value of $x$ satisfying $g'(x) = 0$, the local maximum is also global maximum. This completes the proof of this proposition. □

We now apply our method Padé PSD and compare the estimated PSD with the PSDs obtained analytically and obtained with the averaged periodogram approach. For the numerical results shown in Figure 2(C,D) we set the parameters for the IFF network as

$$I_0 = 1, \quad k_c = 1 \text{ min}^{-1}, \quad k_o = 10 \text{ min}^{-1}, \quad \gamma_c = 1 \text{ min}^{-1}, \quad \gamma_o = 0.5 \text{ min}^{-1},$$
$$\beta_0 = 40 \text{ min}^{-1} \quad \text{and} \quad \beta_{\text{ff}} = 3 \text{ min}^{-1}$$

and for the NFB network as

$$I_0 = 1, \quad k_o = 2 \text{ min}^{-1}, \quad \gamma_c = 1 \text{ min}^{-1}, \quad \gamma_o = 0.5 \text{ min}^{-1},$$
$$\beta_0 = 50 \text{ min}^{-1} \quad \text{and} \quad \beta_{\text{fb}} = 0.5 \text{ min}^{-1}.$$

In order to prevent negativity of the feedforward and feedback functions we set them in the simulation as

$$F_f(x_c) = \max\{\beta_0 - \beta_{\text{ff}} x_c, 0\} \quad \text{and} \quad F_b(x_o) = \max\{\beta_0 - \beta_{\text{fb}} x_o, 0\}. \tag{S89}$$

This enforced positivity adds nonlinearities to the reaction propensities and hence the PSD analytical expressions derived above are no longer exact, but serve as accurate approximations for the chosen parameters. These analytical expressions along with PSDs estimated by our method are given in Table 1 in the main paper. For both these networks, we used the Padé approximant of order $\mathbf{p} = (0, 4)$. For IFF the estimated Padé approximant was

$$G_{\mathbf{p}}(s) = \frac{106.5561s + 113.9207}{s^2 + 1.5101s + 0.5027}$$

while for NFB the Padé approximant was

$$G_{\mathbf{p}}(s) = \frac{66.8159s + 66.6809}{s^2 + 1.497s + 1.4975}.$$

We validated these estimated Padé approximants and the results are shown in Figure S1(C-D).

We now extend our result on differentiating between adapting topologies beyond the three node topologies. Specifically we consider IFF and NFB topologies with arbitrary number of nodes as shown in Figure S2. We first analyse the generalised IFF topology whose reactions are:

$$\mathbf{I} \xrightarrow{k_{c_1} I_0} \mathbf{I} + \mathbf{C_1}, \quad \mathbf{I} \xrightarrow{k_o I_0} \mathbf{I} + \mathbf{O}, \quad \mathbf{C_m} \xrightarrow{F_f(x_{c_m})} \mathbf{C_m} + \mathbf{O}, \quad \mathbf{C_m} \xrightarrow{\gamma_{c_m} x_{c_m}} \emptyset, \quad \mathbf{O} \xrightarrow{\gamma_o x_o} \emptyset, \tag{S90}$$

$$\text{and} \quad \mathbf{C_j} \xrightarrow{k_{c_j} x_{c_j}} \mathbf{C_j} + \mathbf{C_{j+1}}, \qquad \mathbf{C_j} \xrightarrow{\gamma_{c_j} x_{c_j}} \emptyset \quad \text{for} \quad j = 1, \dots, m-1. \tag{S91}$$

Here the quantities above the reaction arrows denote reaction rates, and $x_{c_j}$ denotes the copy-number of species $\mathbf{C_j}$. Notice that the dynamics of $\mathbf{C_1}$ is simply birth-death with production rate $k_{c_1} I_0$ and degradation rate $\gamma_{c_1}$. Hence its steady-state mean is

$$\overline{x}_{c_1} = \mathbb{E}_\pi(C_1) = \frac{k_{c_1} I_0}{\gamma_{c_1}}$$

and its PSD is

$$S_{C_1}(\omega) = \frac{2k_{c_1} I_0}{\gamma_{c_1}^2 + \omega^2}.$$

Observe that for any $j = 1, \dots, m-1$ the production of $\mathbf{C_{j+1}}$ is stimulated by $\mathbf{C_j}$ via a zeroth-order reaction of the form (S43). The steady-state mean of $\mathbf{C_{j+1}}$ is given by

$$\overline{x}_{c_{j+1}} = \mathbb{E}_\pi(C_{j+1}) = \frac{k_{c_j}}{\gamma_{c_{j+1}}} \overline{x}_{c_j}$$

and applying Theorem 3.1 we can relate the PSD of $\mathbf{C_{j+1}}$ to the PSD of $\mathbf{C_j}$ as

$$S_{C_{j+1}}(\omega) = \frac{2k_{c_j} \overline{x}_{c_j}}{\gamma_{c_{j+1}}^2 + \omega^2} + \frac{k_{c_j}^2}{\gamma_{c_{j+1}}^2 + \omega^2} S_{C_j}(\omega).$$

This shows that if $S_{C_j}(\omega)$ is a nonnegative monotonically decreasing function then the same will be true for $S_{C_{j+1}}(\omega)$[4]. Under the assumption of linearity of the feedforward function $F_f$

[4] The class of positive monotonically decreasing functions is closed under multiplication and addition.

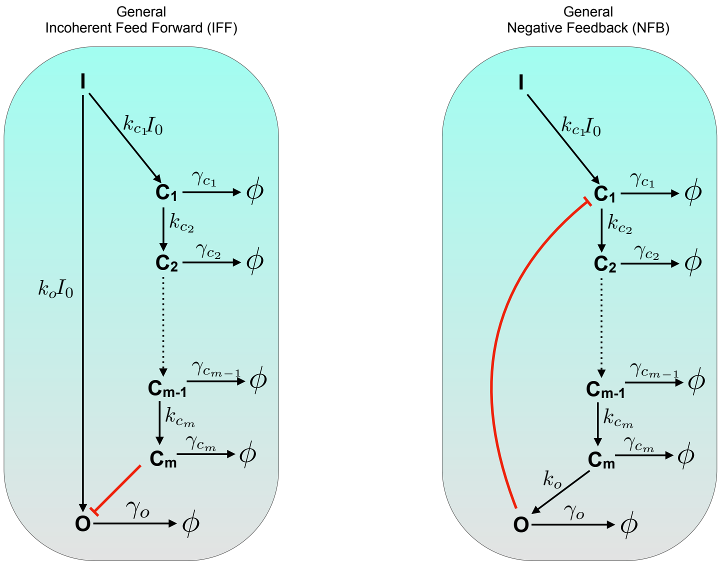

Supplementary Figure S2: IFF and NFB topologies with arbitrary number of nodes.

the stimulation of **O** by $\mathbf{C_m}$ can be viewed as zeroth order degradation. Applying Theorem 3.1 again we can evaluate the output PSD as

$$S_O(\omega) = \frac{2(k_o I_0 + \beta_0 - \beta_{\text{ff}} \overline{x}_{c_m})}{\gamma_o^2 + \omega^2} + \frac{\beta_{\text{ff}}^2}{\gamma_o^2 + \omega^2} S_{C_m}(\omega).$$

Since $S_{C_m}(\omega)$ is nonnegative and monotonically decreasing, we can conclude that $S_O(\omega)$ is also the same. Hence this generalised IFF network cannot exhibit oscillations for any choice of parameters. Of course for $m = 1$ this expression for $S_O(\omega)$ is identical to the expression (S82) derived using formula 12 in the main paper.

We now consider the generalised NFB network whose reactions are given by:

$$\mathbf{I} \overset{I_0 F_b(x_o)}{\longrightarrow} \mathbf{I} + \mathbf{C_1}, \quad \mathbf{C_m} \overset{k_o x_{c_m}}{\longrightarrow} \mathbf{C_m} + \mathbf{O}, \quad \mathbf{C_m} \overset{\gamma_{c_m} x_{c_m}}{\longrightarrow} \emptyset, \quad \mathbf{O} \overset{\gamma_o x_o}{\longrightarrow} \emptyset, \tag{S92}$$

$$\text{and} \quad \mathbf{C_j} \overset{k_{c_j} x_{c_j}}{\longrightarrow} \mathbf{C_j} + \mathbf{C_{j+1}}, \qquad \mathbf{C_j} \overset{\gamma_{c_j} x_{c_j}}{\longrightarrow} \emptyset \quad \text{for} \quad j = 1, \dots, m-1. \tag{S93}$$

As before the quantities above the reaction arrows denote reaction rates. What we shall now argue is that if the input level $I_0$ is high enough, then for this network, the PSD of the output will be non-monotonic which will differentiate it from the IFF network. To see this, note that under the linearisation of the feedback function introduced above, the $(m+1) \times (m+1)$ matrix $A$ is given by

$$A = \begin{bmatrix} -\gamma_{c_1} & 0 & 0 & 0 & -\beta_{\text{fb}} I_0 \\ k_{c_1} & -\gamma_{c_2} & 0 & 0 & 0 \\ \mathbf{0} & \cdot & \ddots & \mathbf{0} & \mathbf{0} \\ 0 & 0 & k_{c_{m-1}} & -\gamma_{c_m} & 0 \\ 0 & 0 & 0 & k_o & -\gamma_o \end{bmatrix}.$$

Therefore the determinant of the matrix $(s\mathbf{I} - A)$ can be expressed as

$$\mathcal{P}(s) = \det(s\mathbf{I} - A) = (s + \gamma_o) \prod_{j=1}^{m} (s + \gamma_{c_j}) + \beta_{\text{fb}} I_0 k_o \prod_{j=1}^{m-1} k_{c_j}.$$

Due to formula (12) and the Cramer's rule for matrix inverse we can see that the output PSD is a rational function whose denominator is given by the polynomial $\mathcal{P}(s)$ evaluated on the imaginary axis. Observe that if $I_0 = 0$ then $\mathcal{P}(s)$ is a polynomial with negative real roots at $-\gamma_o$ and $-\gamma_{c_j}$ for $j = 1, \ldots, m$. The standard root-locus argument [10] shows that as $I_0$ increases, there exists a pair of roots that approach the imaginary axis, and this would imply that the PSD value would approach infinity. This proves that the PSD will be non-monotonic when the input level $I_0$ is high enough. Note that unlike the $m = 1$ case considered before, the system can become unstable if the input $I_0$ exceeds a certain threshold.

#### S4.2 Nonlinear networks

##### S4.2.1 Improving oscillation strength for the *repressilator*

The *repressilator* [6] consists of three genes repressing each other in a cyclic fashion (see Figure 3(A)). We work with an adaptation of the stochastic model given in the Supplement of [22] to demonstrate that our method is able to accurately estimate the single-cell PSD and exhibit the sharpening of the PSD in the presence of sponge. Unsurprisingly this sharpening is also seen if the repressor binding mechanism is made more cooperative. However what is surprising is that with this more cooperative binding mechanism, the sponge may have an opposite effect, i.e. instead of sharpening the PSD it may actually broaden it. We now describe the stochastic model in detail.

In what follows, $p_1$, $p_2$ and $p_3$ denote the molecular counts of proteins TetR, cI and LacI. The dynamics of three repressor proteins is given by *bursty* gene-expression

$$p_{i,\text{tot}} \xrightarrow{\lambda_i(p_{j,\text{free}})} p_{i,\text{tot}} + 1 \quad \text{and} \quad p_{i,\text{tot}} \xrightarrow{1} p_{i,\text{tot}} - 1, \tag{S94}$$

for $(i, j) = (1, 3), (2, 1)$ and $(3, 2)$. Here

$$\lambda_i(p_{j,\text{free}}) = \lambda N_0 \frac{K_i^H}{K_i^H + p_{j,\text{free}}^H}$$

is the Hill function encoding the repression in the production of *total* protein count $p_i$ by the *free* (i.e. unbound to the plasmid) proteins $p_j$. The Hill coefficient $H$ represents the degree of cooperativity among promoter binding sites. The degradation rate of 1 gen.$^{-1}$ is assumed[5] for all the species, and it corresponds to dilution and cell-division.

Let $N_0$ be the number of repressilator plasmids and $N_t$ be the number of sponge plasmids. Their birth-death dynamics is given by

$$\emptyset \xrightarrow{\langle N_0 \rangle} N_0 \xrightarrow{1} \emptyset \quad \text{and} \quad \emptyset \xrightarrow{\langle N_t \rangle} N_t \xrightarrow{1} \emptyset, \tag{S95}$$

[5] Here gen. is abbreviation for generations.

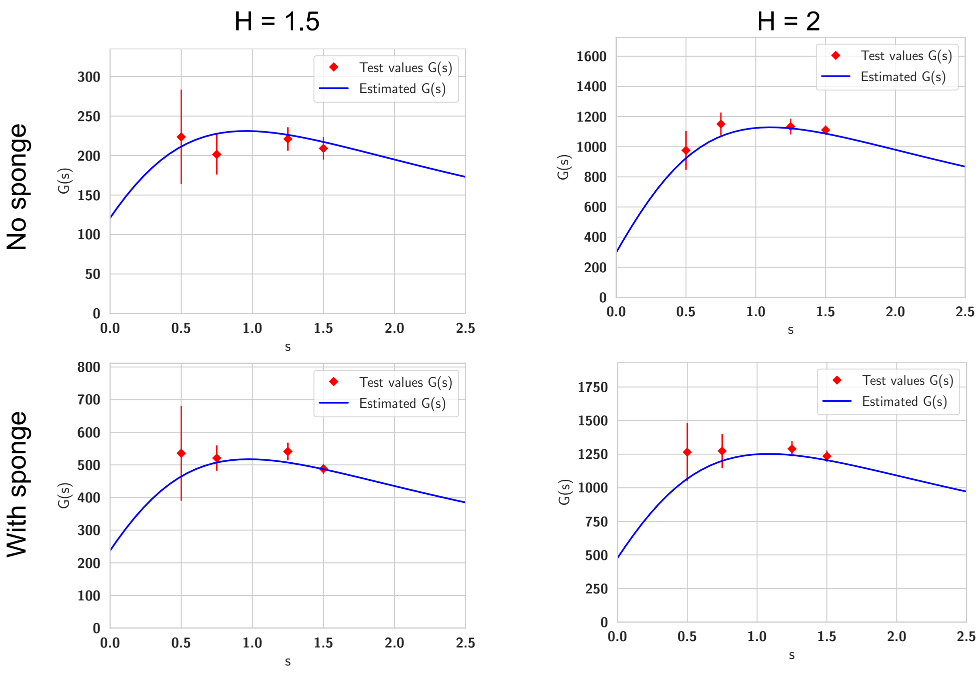

Supplementary Figure S3: Validation of the estimated Padé approximant $G_{\mathbf{p}}(s)$ for the repressilator network. Direct estimates for $G(s)$ are plotted in red for four test values $\mathbf{s} = (0.5, 0.75, 1.25, 1.5)$. The point denotes the estimator mean and the error bar denotes the symmetric one standard deviation interval. The blue curve denotes the estimated Padé approximant.

where $\langle N_0 \rangle$ and $\langle N_t \rangle$ are deterministic constants representing the mean number of repressilator and sponge plasmids respectively.

Let $N_{\text{pro},j}$ be the total number of promoters available for binding repressor protein $p_j$. Then

$$N_{\text{pro},j} = \begin{cases} N_0 + N_t & \text{for } j = 1 \ (tetR) \\ N_t & \text{for } j = 2, 3 \ (cI \text{ and } lacI) \end{cases}$$

and the number of bound molecules of repressor $p_j$ is proportional to

$$N_{\text{pro},j} \frac{p_{j,\text{free}}^H}{K^H + p_{j,\text{free}}^H}.$$

Assuming two binding sites per promoter, the number of free and total repressor protein molecules are related by

$$p_{j,\text{tot}} = p_{j,\text{free}} + 2N_{\text{pro},j} \frac{p_{j,\text{free}}^H}{K^H + p_{j,\text{free}}^H}. \tag{S96}$$

Using Gillespie's SSA we can simulate reactions (S94)-(S95) to generate a stochastic trajectory. Every time $p_{j,\text{tot}}$ changes due to a reaction in (S94), we update the free protein count by solving the nonlinear equation (S96) via the bisection method. We set the rate parameters as

$$\lambda = 30, \quad K_1 = 5, \quad K_2 = K_3 = 10, \quad H = 1.5, \quad \langle N_0 \rangle = 10 \quad \text{and} \qquad \text{(S97)}$$
$$\langle N_t \rangle = \begin{cases} 40 & \text{with sponge} \\ 0 & \text{without sponge.} \end{cases}$$

This choice of parameters is the same as in the Supplement of [22], except that we pick the mean burst size $\langle b \rangle$ to be 1 instead of 10 in order to keep the copy-numbers low and preserve the effects of intrinsic noise. The time unit for $\lambda$, $\langle N_0 \rangle$ and $\langle N_t \rangle$ is gen.$^{-1}$.

We use our method to estimate the PSD for the dynamics of the copy-numbers of the cI protein, and the results are reported in Figure 3. Next we increased the promoter co-operativity (i.e. the Hill coefficient) from $H = 1.5$ to $H = 2$ and repeated the same PSD estimation analysis. The estimated Padé approximants and the PSDs are given in Table S1 for both values of $H$.

| Hill coefficient | Case | Order $\mathbf{p} = (p_1, p_2)$ | Padé approximant $G_{\mathbf{p}}(s)$ | Estimated PSD $\widehat{S}_{\text{cI}}(\omega)$ |
|---|---|---|---|---|
| $H = 1.5$ | No sponge | $(0, 4)$ | $\frac{564.123s+232.0576}{s^2+0.5316s+1.916}$ | $\frac{135.6873\omega^2+889.2624}{\omega^4-3.5495\omega^2+3.6712}$ |
| | With sponge | $(0, 4)$ | $\frac{1228.3762s+418.7465}{s^2+0.4226s+1.7616}$ | $\frac{200.8428\omega^2+1475.3663}{\omega^4-3.3447\omega^2+3.1034}$ |
| $H = 2$ | No sponge | $(2, 2)$ | $\frac{2730.0273s+493.5059}{s^2+0.2106s+1.6555}$ | $\frac{162.899\omega^2+1633.9979}{\omega^4-3.2666\omega^2+2.7407}$ |
| | With sponge | $(2, 2)$ | $\frac{3119.1301s+905.1255}{s^2+0.3235s+1.8972}$ | $\frac{208.006\omega^2+3434.4757}{\omega^4-3.6898\omega^2+3.5995}$ |

Supplementary Table S1: Expressions for the Padé approximants and the PSDs estimated with the Padé PSD method for the repressilator network. The estimated Padé approximants were validated and the results are shown in Figure S3.

##### S4.2.2 Reducing single-cell oscillations due to the antithetic integral feedback controller

We start by describing the AIF controller in greater detail. It consists of two bio-molecular control species $\mathbf{Z_1}$ and $\mathbf{Z_2}$ and the following four elementary reactions with mass-action kinetics:

$$\emptyset \xrightarrow{\mu} \mathbf{Z_1} \quad \text{(reference)} \qquad \text{(S98)}$$
$$\mathbf{X}_\ell \xrightarrow{\theta} \mathbf{X}_\ell + \mathbf{Z_2} \quad \text{(output sensing)} \qquad \text{(S99)}$$
$$\mathbf{Z_1} + \mathbf{Z_2} \xrightarrow{\eta} \emptyset \quad \text{(annihilation)} \qquad \text{(S100)}$$
$$\mathbf{Z_1} \xrightarrow{k} \mathbf{Z_1} + \mathbf{X_1} \quad \text{(actuation).} \qquad \text{(S101)}$$

Here rate constants are indicated above the reaction arrows and $\mathbf{X_1}$ is the actuated species which, through an arbitrary set of reactions and intermediate species, positively affects the copy-numbers of the output species $\mathbf{X}_\ell$. The AIF controller robustly steers the mean copy-number level of $\mathbf{X}_\ell$ to the desired set-point $\mu/\theta$, where $\mu$ is the production rate of $\mathbf{Z_1}$ (see reaction (S98)) and $\theta$ is the reaction rate constant for the output sensing reaction (S99). The AIF affects the output by actuating the production of $\mathbf{X_1}$ (see reaction (S101)) and the feedback loop is closed by the annihilation reaction (S100) between $\mathbf{Z_1}$ and $\mathbf{Z_2}$.

In our example, the link between $\mathbf{X_1}$ and $\mathbf{X}_\ell$ (which forms the controlled network) is given by the classical gene-expression model considered in Section S4.1.1. Hence the actuated species $\mathbf{X_1}$ is the mRNA and the output species $\mathbf{X_2}$ is the protein. The schematics of the AIF connected to the gene-expression network is shown in Figure 4(A). Letting $z_1$ and $x_2$ denote the copy-numbers of $\mathbf{Z_1}$ and $\mathbf{X_2}$ respectively, we add the extra feedback by changing the rate of the actuation reaction (S101) from $kz_1$ to $(kz_1 + F_b(x_2))$ where $F_b$ is the feedback function which takes non-negative values and it is monotonically decreasing. We consider two types of feedback. The first is the *Hill* feedback

$$F_b(x_2) = \frac{4k_{\text{fb}}\left(\frac{\mu}{\theta}\right)^2}{\frac{\mu}{\theta} + x_2}$$

while the second is the *proportional* feedback

$$F_b(x_2) = k_{\text{fb}} \max\left\{\frac{3\mu}{\theta} - x_2, 0\right\}$$

which is based on the deviation of the output $x_2$ from the set-point $\mu/\theta$.

We set the controller parameters as

$$\mu = 10 \text{ min}^{-1}, \quad \theta = 1 \text{ min}^{-1}, \quad \eta = 100 \text{ min}^{-1} \quad \text{and} \quad k = 5 \text{ min}^{-1}$$

and the gene-expression network parameters as

$$k_r = 0 \text{ min}^{-1}, \quad k_p = 2 \text{ min}^{-1}, \quad \gamma_r = 2 \text{ min}^{-1} \quad \text{and} \quad \gamma_p = 1 \text{ min}^{-1}.$$

Setting parameter $k_r$ to 0 implies that all production of mRNA is due to the actuation reaction (S101) which ensures that any positive set-point is achievable in the absence of extra feedback. We use our method to estimate the PSD for the single-cell protein dynamics in the AIF regulated gene-expression network for both types of feedback and for different values of the gain parameter $k_{\text{fb}}$. All the estimated Padé approximants and the PSDs are given in Table S2.

##### S4.2.3 Inferring cooperativity in a self-regulated gene-expression network

Consider a self-regulated gene-expression system modelled as a single-species birth-death model

$$\emptyset \xrightarrow{\lambda_H(x)} \mathbf{X} \xrightarrow{\delta} \emptyset.$$

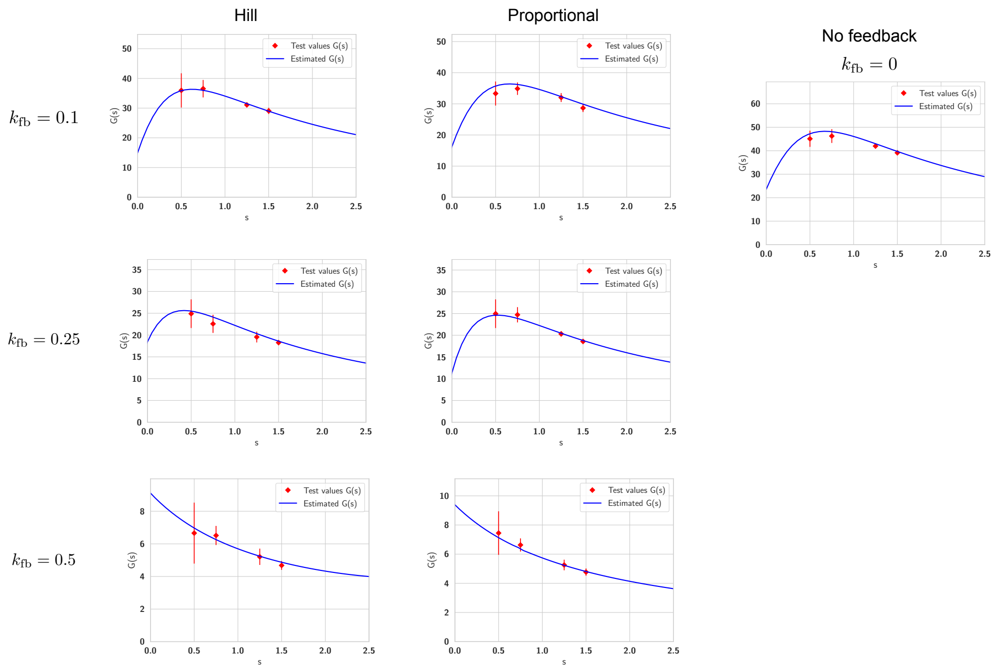

Supplementary Figure S4: Validation of the estimated Padé approximant $G_{\mathbf{p}}(s)$ for the antithetic integral feedback controller. Direct estimates for $G(s)$ are plotted in red for four test values $\mathbf{s} = (0.5, 0.75, 1.25, 1.5)$. The point denotes the estimator mean and the error bar denotes the symmetric one standard deviation interval. The blue curve denotes the estimated Padé approximant.

The production rate is set by the repressing Hill function

$$\lambda_H(x) = \frac{K_0}{K_1 + x^H}$$

and the degradation reaction has mass-action kinetics with rate constant $\gamma$. The Hill coefficient $H$ represents the degree of cooperativity in the binding of $\mathbf{X}$ molecules to the repressor sites. In our example we set $K_0 = K_1 = 10$ and $\gamma = 0.05$, and our aim is to infer the parameter $H$ from the normalised PSD, computed with 100 "experimental" trajectories that are obtained by simulating and discrete-sampling the reaction dynamics with $H = 1$.

The inference is based on simple comparison of the experimental PSD with the normalised PSDs obtained by Padé PSD for various values of $H$. The estimated Padé approximants and the PSDs are given in Table S3 for different values of $H$.

| Feedback type | Feedback parameter $k_{\text{fb}}(\text{min}^{-1})$ | Order $\mathbf{p} = (p_1, p_2)$ | Padé approximant $G_{\mathbf{p}}(s)$ | Estimated PSD $\widehat{S}_{\mathbf{X_2}}(\omega)$ |
|---|---|---|---|---|
| None | 0 | $(4, 0)$ | $\frac{88.4703s+20.8184}{s^2+0.4894s+0.8817}$ | $\frac{44.951\omega^2+36.7113}{\omega^4-1.5239\omega^2+0.7774}$ |
| Hill | 0.1 | $(4, 0)$ | $\frac{66.5605s+9.9109}{s^2+0.5839s+0.6618}$ | $\frac{57.911\omega^2+13.1174}{\omega^4-0.9826\omega^2+0.4379}$ |
| | 0.25 | $(4, 0)$ | $\frac{45.4898s+11.4944}{s^2+0.9424s+0.622}$ | $\frac{62.7484\omega^2+14.2998}{\omega^4-0.356\omega^2+0.3869}$ |
| | 0.5 | $(4, 0)$ | $\frac{12.7138s-57.2821}{s^2-2.5365s-6.2874}$ | $\frac{50.0681\omega^2+720.3121}{\omega^4+19.0085\omega^2+39.5315}$ |
| Proportional | 0.1 | $(4, 0)$ | $\frac{70.8937s+12.7146}{s^2+0.6333s+0.7812}$ | $\frac{64.3655\omega^2+19.8652}{\omega^4-1.1613\omega^2+0.6103}$ |
| | 0.25 | $(4, 0)$ | $\frac{46.977s+5.7793}{s^2+0.8655s+0.5062}$ | $\frac{69.7582\omega^2+5.8511}{\omega^4-0.2633\omega^2+0.2563}$ |
| | 0.5 | $(4, 0)$ | $\frac{14.8071s-11.1854}{s^2+0.8228s-1.1919}$ | $\frac{46.7375\omega^2+26.6647}{\omega^4+3.0609\omega^2+1.4207}$ |

Supplementary Table S2: Expressions for the Padé approximants and the PSDs estimated with the Padé PSD method for the antithetic integral feedback controller. The estimated Padé approximants were validated and the results are shown in Figure S4.

| Hill coefficient | Order $\mathbf{p} = (p_1, p_2)$ | Padé approximant $G_{\mathbf{p}}(s)$ | Estimated PSD $\widehat{S}_{\text{cI}}(\omega)$ |
|---|---|---|---|
| $H = 0$ | $(0, 2)$ | $\frac{18.0177}{s+0.0504}$ | $\frac{1.8149}{\omega^2+0.0025}$ |
| $H = 0.5$ | $(0, 4)$ | $\frac{13.1295s+6.1666}{s^2+0.525s+0.026}$ | $\frac{1.4526\omega^2+0.3206}{\omega^4+0.2236\omega^2+0.00068}$ |
| $H = 1$ | $(0, 4)$ | $\frac{6.7398s+1.1397}{s^2+0.2441s+0.0127}$ | $\frac{1.0116\omega^2+0.0289}{\omega^4+0.0343\omega^2+0.0002}$ |
| $H = 1.5$ | $(0, 4)$ | $\frac{3.5941s+0.7492}{s^2+0.3085s+0.0207}$ | $\frac{0.7187\omega^2+0.031}{\omega^4+0.0537\omega^2+0.0004}$ |
| $H = 2$ | $(0, 4)$ | $\frac{2.1647s+0.8615}{s^2+0.5251s+0.0499}$ | $\frac{0.5505\omega^2+0.0859}{\omega^4+0.176\omega^2+0.0025}$ |

Supplementary Table S3: Expressions for the Padé approximants and the PSDs estimated with the Padé PSD method for the self-regulated gene-expression network. The estimated Padé approximants were validated and the results are shown in Figure S6.

[2] S. Asmussen and P. W. Glynn. *Stochastic simulation: algorithms and analysis*, volume 57. Springer Science & Business Media, 2007.

[3] J. B. Conway. *A course in functional analysis.* Springer, 1985.

[4] J. W. Cooley and J. W. Tukey. An algorithm for the machine calculation of complex fourier series. *Mathematics of computation*, 19(90):297–301, 1965.

[5] T. Cormen, C. Leiserson, R. Rivest, and C. Stein. *Introduction to Algorithms*. McGraw-Hill Science/Engineering/Math, New York, second edition, 2003.

[6] M. B. Elowitz and S. Leibler. A synthetic oscillatory network of transcriptional regulators. *Nature*, 403(6767):335–338, 2000.

[7] M. B. Elowitz, A. J. Levine, E. D. Siggia, and P. S. Swain. Stochastic gene expression in a single cell. *Science*, 297(5584):1183–1186, 2002.

[8] S. Engelberg. *Digital signal processing: an experimental approach*. Springer Science & Business Media, 2008.

[9] S. N. Ethier and T. G. Kurtz. *Markov processes : Characterization and Convergence*. Wiley Series in Probability and Mathematical Statistics: Probability and Mathematical Statistics. John Wiley & Sons Inc., New York, 1986.

[10] G. F. Franklin, J. D. Powell, A. Emami-Naeini, and J. D. Powell. *Feedback control of dynamic systems*, volume 4. Prentice hall Upper Saddle River, 2002.

[11] D. T. Gillespie. Exact stochastic simulation of coupled chemical reactions. *The Journal of Physical Chemistry*, 81(25):2340–2361, 1977.

[12] A. Gupta, C. Briat, and M. Khammash. A scalable computational framework for establishing long-term behavior of stochastic reaction networks. *PLoS Comput Biol*, 10(6):e1003669, 06 2014.

[13] A. Gupta and M. Khammash. Computational identification of irreducible state-spaces for stochastic reaction networks. *SIAM Journal on Applied Dynamical Systems*, 17(2):1213–1266, 2018.

[14] A. G. Hart and R. L. Tweedie. Convergence of invariant measures of truncation approximations to markov processes. *Applied Mathematics*, 3(12):2205, 2012.

[15] T. Kato. *Perturbation theory for linear operators*, volume 132. Springer Science & Business Media, 2013.

[16] A. Khintchine. Korrelationstheorie der stationären stochastischen prozesse. *Mathematische Annalen*, 109(1):604–615, 1934.

[17] H. H. McAdams and A. Arkin. Stochastic mechanisms in gene expression. *Proc. Natl. Acad. Sci., Biochemistry*, 94:814–819, 1997.

[18] J. McCabe and J. Murphy. Continued fractions which correspond to power series expansions at two points. *IMA Journal of Applied Mathematics*, 17(2):233–247, 1976.

[19] S. P. Meyn and R. L. Tweedie. Stability of Markovian processes. III. Foster-Lyapunov criteria for continuous-time processes. *Adv. in Appl. Probab.*, 25(3):518–548, 1993.

[20] J. R. Norris. *Markov chains*, volume 2 of *Cambridge Series in Statistical and Probabilistic Mathematics*. Cambridge University Press, Cambridge, 1998. Reprint of 1997 original.

[21] H. Nyquist. Certain topics in telegraph transmission theory. *Transactions of the American Institute of Electrical Engineers*, 47(2):617–644, 1928.

[22] L. Potvin-Trottier, N. D. Lord, G. Vinnicombe, and J. Paulsson. Synchronous long-term oscillations in a synthetic gene circuit. *Nature*, 538(7626):514–517, 2016.

[23] A. Sidi. Some aspects of two-point padé approximants. *Journal of Computational and Applied Mathematics*, 6(1):9–17, 1980.

[24] M. Thattai and A. van Oudenaarden. Intrinsic noise in gene regulatory networks. *Proceedings of the National Academy of Sciences*, 98(15):8614–8619, 2001.

[25] K. Yosida. *Functional analysis*. Springer, 1995.

**Algorithm 5** Given the discrete-sampled augmented CTMC trajectories $\widehat{\mathcal{X}}_q = (\mathcal{X}_q(t_1), \ldots, \mathcal{X}_q(t_{N_f}))$ for $q = 1, \ldots, Q$, this method computes Monte Carlo estimates of the Padé derivatives $D_{m_1}^{(s_0)}$ and $D_{m_2}^{(\infty)}$ for $m_1 = 0, 1, \ldots, (p_1 - 1)$ and $m_2 = 0, 1, \ldots, (p_2 - 1)$, and the direct estimates $G(s_r)$ for $r = 1, \ldots, R$.

**function** MONTECARLOESTIMATOR($\widehat{\mathcal{X}}_1, \ldots, \widehat{\mathcal{X}}_Q$)
  **for** $q = 1, \ldots, Q$ **do**
    Set $\text{Mean}_q = D_{0,q}^{(\infty)} = \cdots = D_{p_2-1,q}^{(\infty)} = D_{0,q}^{(s_0)} = \cdots = D_{p_1-1,q}^{(s_0)} = G_{1,q} = \cdots = G_{R,q} = 0$
    **for** $j = 1, \ldots, N_f$ **do**
      Set $(x, y, z) = \mathcal{X}_q(t_j)$
      Update $\text{Mean}_q \leftarrow \text{Mean}_q + f(x)$ and $D_{0,q}^{(\infty)} \leftarrow D_{0,q}^{(\infty)} + (f(x))^2$
      **for** $m = 0, \ldots, (p_1 - 1)$ **do**
        Update $D_{m,q}^{(s_0)} \leftarrow D_{m,q}^{(s_0)} + f(x) y_{m+1}$
      **end for**
      **for** $m = 1, \ldots, (p_2 - 1)$ **do**
        Set $\psi =$ COMPUTE-PSI$(x, m)$
        Update $D_{m,q}^{(\infty)} \leftarrow D_{m,q}^{(\infty)} + \psi$
      **end for**
      **for** $r = 1, \ldots, R$ **do**
        Update $G_{r,q} \leftarrow G_{r,q} + f(x) z_r$
      **end for**
    **end for**
    Update $\text{Mean}_q \leftarrow \text{Mean}_q / N_f$
    Update $D_{0,q}^{(\infty)} \leftarrow D_{0,q}^{(\infty)} / N_f - (\text{Mean}_q)^2$
    **for** $r = 1, \ldots, R$ **do**
      Update $G_{r,q} \leftarrow (G_{r,q}/N_f - (\text{Mean}_q)^2)/s_r$
    **end for**
    **for** $m = 0, \ldots, (p_1 - 1)$ **do**
      Update $D_{m,q}^{(s_0)} \leftarrow (D_{m,q}^{(s_0)}/N_f - (\text{Mean}_q)^2)/s_0^{m+1}$
    **end for**
  **end for**
**return** Monte Carlo estimates

$$\widehat{D}_m^{(s_0)} := \frac{1}{Q} \sum_{q=1}^{Q} \widehat{D}_{m,q}^{(s_0)}, \quad \text{for} \quad m = 0, \ldots, (p_1 - 1)$$

$$\widehat{D}_m^{(\infty)} := \frac{1}{Q} \sum_{q=1}^{Q} \widehat{D}_{m,q}^{(\infty)} \quad \text{for} \quad m = 0, \ldots, (p_2 - 1)$$

$$\text{and} \quad \widehat{G}_r := \frac{1}{Q} \sum_{q=1}^{Q} G_{r,q} \quad \text{for} \quad r = 1, \ldots, R$$

**end function**

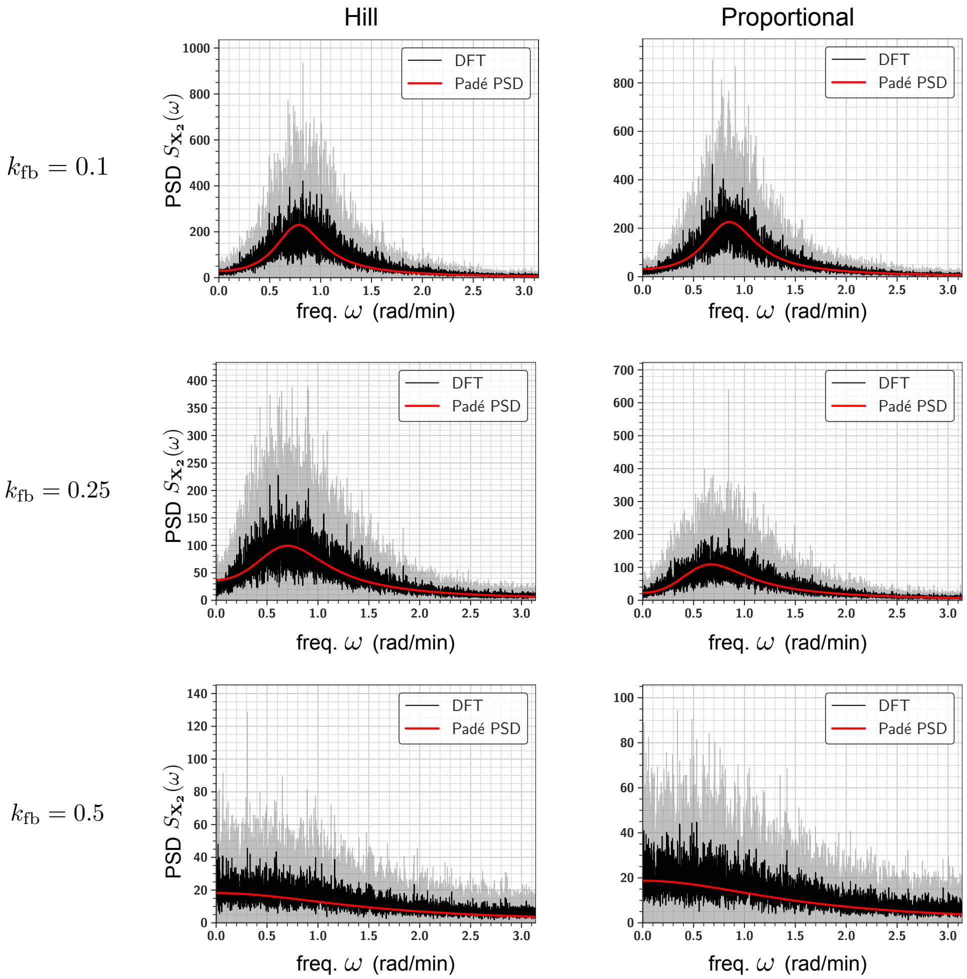

Supplementary Figure S5: Comparison of the PSDs estimated with the Padé PSD method and the DFT method for the Hill and proportional feedback for three choices of feedback parameter $k_{\mathrm{fb}}$.

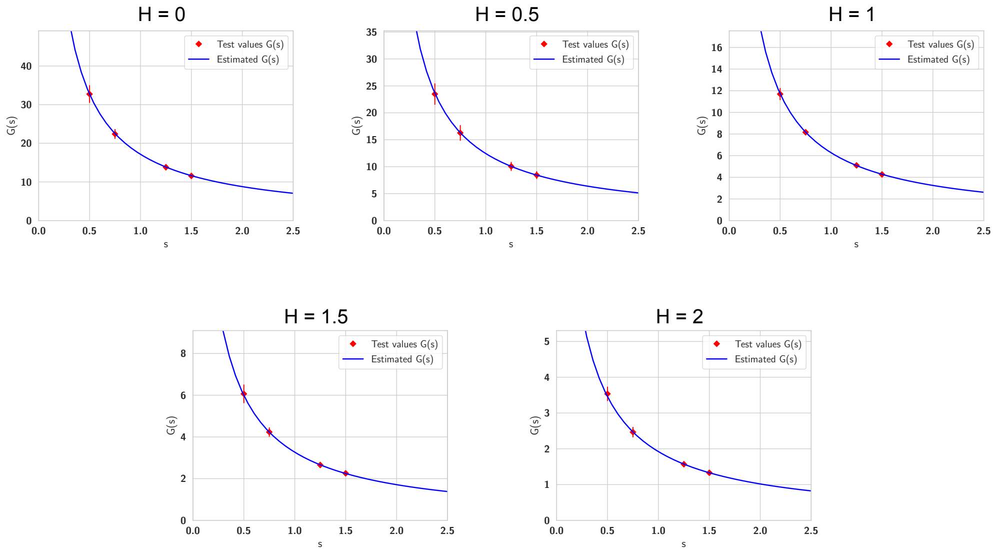

Supplementary Figure S6: Validation of the estimated Padé approximant $G_{\mathbf{p}}(s)$ for the self-regulated gene-expression network. Direct estimates for $G(s)$ are plotted in red for four test values $\mathbf{s} = (0.5, 0.75, 1.25, 1.5)$. The point denotes the estimator mean and the error bar denotes the symmetric one standard deviation interval. The blue curve denotes the estimated Padé approximant.
